# SuSiNE: Genetic fine-mapping with signed functional priors and multi-basin ensembling

**DOI:** 10.64898/2026.07.31.742084

**Authors:** Michael G. Callahan, Xiang Zhu

**Affiliations:** Department of Statistics, The Pennsylvania State University, University Park, Pennsylvania, United States of America

## Abstract

Genetic fine-mapping identifies causal variants within trait-associated loci, but linkage disequilibrium (LD) and wide datasets complicate this sparse variable-selection problem. SuSiE is popular for its fast variational inference, posterior inclusion probabilities (PIPs), and credible sets, yet a single fit can fail to resolve LD ambiguity, converge to a poor local optimum, or misrepresent uncertainty over competing configurations. We introduce SuSiNE (Sum of Single Non-central Effects), a SuSiE extension incorporating signed functional annotations through a prior-mean channel, ***µ***_0_ = *c****a***, while preserving effect conjugacy, credible sets, and summary-statistic sufficiency. The resulting single-effect Bayes factor self-gates on agreement between annotation sign and association direction, limiting annotation-noise influence. We show that the common final step of purity filtering can discard informative signal, and tends to hurt performance. We also introduce new effect-level diagnostics for concentration, accuracy, and fitted-basis movement, to provide deeper insights into model behavior. To explore and summarize multiple variational basins, we pair the model with grid-based ensembling and cluster-weight aggregation. In oligogenic simulations with annotations calibrated to AlphaGenome eQTL benchmarks, the ensemble raised pooled AUPRC for recovery of the largest-effect causal variants from a SuSiE-equivalent 0.2474 to 0.3130 (0.0656 delta, 95% paired-bootstrap CI [0.0591, 0.0722]). At 75% precision, recall rose from 11.9% to 19.3% (61.7% relative gain). AUPRC gains were robust across varying annotation quality and alternative sparse and diffuse architectures, while sufficiently strong null annotation-association alignment reversed the gains. In a GTEx Lung summary-statistic case study, SuSiNE placed nontrivial weight on annotation-informed fits at 7 of 20 loci and changed which variants received high PIP. *ARSA* showed the cleanest durable shift, whereas the large *YDJC* shift coincided with reference-LD discrepancy. An internal diagnostic found little evidence of strong annotation confounding in this panel. These analyses use reference rather than in-cohort LD, demonstrating method behavior rather than definitive variant-level discoveries.

**Author summary:** When a genetic study links part of the genome to a disease or to differences in gene expression, the next question is which variants are responsible. Answering this is hard, because nearby variants are usually inherited together and can look almost interchangeable in the data. We studied a widely used method, SuSiE, by asking where it breaks down. We found that a routine final cleanup step often discards real signal for nothing in return. A single run can also settle on one explanation without exploring alternatives that fit the data just as well. We introduce new checks that make both problems visible. We then developed SuSiNE, which lets the method use directional predictions from AI sequence models or other biological evidence. It runs many times across settings that encourage exploration, then combines the results into one summary. In calibrated simulations, SuSiNE found true causal variants substantially more often than the standard method. On real gene-expression data, it changed which variants look responsible at several locations. These results are limited, but they suggest AI sequence models are already good enough to offer competing explanations at well-studied genome locations, if we use them carefully.

## 1 Introduction

### 1.1 Background

Genome-wide association studies (GWAS), expression quantitative trait loci (eQTL) mapping, and other genetic association studies have identified thousands of genomic loci associated with complex traits and diseases [1–3]. Pinpointing the specific causal variants within these loci remains a significant challenge [4]. Fine-mapping regression models search a locus for causal variants, and the difficulty of that search depends heavily on the correlation structure of the candidates. The most widely appreciated obstacle is linkage disequilibrium (LD). Highly correlated variants can provide nearly interchangeable explanations under the available association evidence. A model may estimate their shared fitted signal *ŷ* accurately while remaining unable to attribute it confidently to one variant. This is fundamentally a limitation of the data, and resolving it requires external information.

Fine-mapping must also search a combinatorial space of causal configurations. Exact variable selection for regression problems is NP-hard [5], so practical methods can explore only a small slice of the total search space. However, NP-hardness does not by itself rule out useful and efficient approximations. When variants are uncorrelated, the variance-explained objective is modular, where each variant has an independent marginal contribution to *R*^2^, and greedy selection is exact. Under the weaker condition of submodularity, which formalizes diminishing returns, greedy selection still has a (1 − 1*/e*) approximation guarantee for the objective [6].

Statistical fine-mapping generally does not have either of these helpful structures. Sparse-regression *R*^2^ is submodular only under restrictive covariance conditions [7]. In high-dimensional genomic settings, the modest ratio of sample size to variant count can drive the submodularity ratio well below one [8]. The objective is then typically far from submodular in the approximate sense needed to support a useful greedy guarantee. Greedy procedures such as the IBSS algorithm introduced below can still be fast and effective heuristics, but there is no theoretical guarantee that they will select the best basin or the best causal configuration. It is important to note that this core difficulty of multimodal optimization in a combinatoric space would be present in fine-mapping even without the ambiguities introduced by linkage-disequilibrium.

A fine-mapping method must consequently do two things. It must search a large and poorly structured model space efficiently, and it must represent uncertainty when several variant sets are similarly plausible. This is the classical Bayesian variable-selection and model-averaging problem. Posterior mass is assigned to competing subsets, and per-variant posterior inclusion probabilities are obtained by marginalizing over those subsets [9, 10]. Earlier fine-mapping methods often represent a distribution over discrete causal configurations directly [11–14]. This representation is expressive, but enumerating or sampling the configurations becomes expensive as the locus grows.

The representational choice is easiest to understand through a small example: consider four variants arranged in two highly correlated pairs. Suppose one variant is causal within each pair, while the pairing itself is clear. The posterior can be written as AND{OR(1, 2), OR(3, 4)}. There are two local choices, and both choices must be made. We call this an *AND-of-ORs* representation. The same posterior could instead list four complete alternatives. They are AND(1, 3), AND(1, 4), AND(2, 3), and AND(2, 4). Taking the OR over these four alternatives gives what we call an *OR-of-ANDs* representation. It is longer in this simple case because it enumerates every combination of the two local choices.

This difference in representation matters when the choices require coordination. Suppose six variants admit only two plausible three-variant configurations, {1, 3, 5} and {2, 4, 6}. Their uncertainty is naturally written as OR{AND(1, 3, 5), AND(2, 4, 6)}. An AND-of-ORs representation that spreads mass independently within three components also creates mixed triplets such as {1, 4, 5} or {2, 3, 6}, even though those configurations may have no statistical support. Adding components does not repair this mismatch because it adds more independent choices and more opportunities to place mass on unsupported hybrids. OR-of-ANDs can represent any discrete collection of competing configurations, but it is combinatorially costly to explore. AND-of-ORs is compact when uncertainty decomposes into independent local choices, but it cannot retain every coordinated choice among whole configurations. S1 Appendix formalizes this distinction.

SuSiE’s unique approach takes the efficient AND-of-ORs route [15]. It decomposes the regression into *L* single-effect components and fits a product-form variational posterior with iterative Bayesian stepwise selection (IBSS). This continuous relaxation operates on a probability-weight matrix over components and variants, rather than on a long list of discrete variant sets. It yields per-variant PIPs and per-component credible sets, which has made SuSiE practical for large fine-mapping studies. Its factorized posterior represents independent choices within components. A single fit cannot, however, represent OR-of-ANDs uncertainty that requires coordinated choices across distinct configurations.

### 1.2 Problems

Despite these computational advantages, SuSiE has three distinct pitfalls that can limit its performance. Two concern the geometry of the variational fit, so we first introduce vocabulary that will be used throughout the paper. SuSiE fits its model by maximizing an evidence lower bound (ELBO) with IBSS. It is useful to separate two layers of the resulting fit. The first is the *explanatory basis*, which is the set of fitted signal directions **ŷ***_ℓ_* explained by the *L* per-effect components. It describes what signal in ℝ*^n^* the model has accounted for, regardless of which variant receives credit. The second is the *PIP vector*. Given an explanatory basis, the PIP vector describes how posterior inclusion mass is distributed among variants that could account for each fitted direction. We call one converged IBSS solution an *optimum*. It consists of one explanatory basis paired with one PIP vector and corresponds to an ELBO stationary point in SuSiE’s continuously relaxed variational space.

We use *basin* to describe a region of the ELBO landscape in which the explanatory basis is essentially fixed. Movement within a basin can redistribute PIP mass among correlated variants while leaving the fitted signal nearly unchanged. Movement across basins changes the explanatory basis and therefore changes the configuration of fitted directions. This distinction helps classify differences between multiple SuSiE fits: some changes are between which variants account for the same explanatory signal, while some changes are between which explanatory signals the model finds. The SuSiE model therefore places a bet about where configuration uncertainty lives: that most of it falls between variants sharing one explanatory basis, rather than between competing bases. If the bet is right, then the model gains computational efficiency without losing much accuracy. If the bet is wrong, then the model can fail to find the best basin or fail to represent uncertainty across basins.

The first pitfall SuSiE faces is LD ambiguity, which is a well documented phenomenon and not exclusive to the SuSiE method. The genotype data alone may not distinguish tightly correlated variants, so PIP mass moves within a basin even though the fitted signal is stable. The second is a *model-specification barrier*. Even at a well-optimized fit, one factorized optimum cannot express OR-of-ANDs uncertainty across distinct configurations in different basins. The third is an *optimizer barrier*. IBSS can converge to an inferior basin whose variants look attractive in the first marginal updates, even when another basin fits the data much better. The second and third barriers are SuSiE-specific. We demonstrate these failures in small toy examples and then look for their diagnostic signatures in realistic simulations and GTEx Lung eQTL data.

The credible-set summaries in common use can blur these different failures. Size, power, coverage, and purity are easy to interpret, but they treat important aspects of the posterior rather coarsely. Purity is the minimum pairwise absolute correlation among variants in a credible set, and a threshold of 0.50 is commonly used to filter fitted output [15, 16]. The rationale is sensible: very low-purity sets may reflect extra diffuse effects introduced when *L* is larger than the true number of signals [15]. However, low purity can also arise when a real fitted effect places concentrated mass on variants that are weakly correlated, but nonetheless indistinguishable under the realized association evidence. In the controlled example below, purity filtering removes such an effect. In the realistic baseline simulation, the same filter lowers causal-recovery AUPRC from 0.248 to 0.189.

Size, power, and coverage have a related limitation because they treat the credible set as a flat collection. Two alternative sets with identical membership receive the same scores, even if one of them concentrates almost all PIP mass on one variant and the other spreads mass evenly across the set. We therefore complement these credible set-level summaries with effect-level diagnostics for concentration, accuracy, and movement of the fitted basis.

### 1.3 Our contributions

High-LD ambiguity is a data problem, so resolving it requires external information that can distinguish variants whose genotypes cannot. The base SuSiE method exposes one channel for such information through the prior inclusion probabilities ***π***. PolyFun uses this channel for unsigned per-variant heritability estimates [17]. Related methods place functional annotations, supervised functional scores, or sequence-model predictions in the same type of inclusion prior [18–20]. This is a natural channel for unsigned information, but it cannot natively retain the direction of a signed annotation. A variant predicted to strongly increase expression receives the same inclusion prior as one predicted to decrease it by the same amount, even though the two predictions have opposite implications for an expression phenotype.

Directional regulatory information has already proved useful at the genome-wide annotation level [21], and modern sequence-to-function (S2F) models now produce signed variant-effect predictions. Enformer, Borzoi, Decima, and AlphaGenome represent successive advances in this model class [22–25] made in recent years. Their zero-shot accuracy improvements have direct implications for fine-mapping, and our later results show that the AlphaGenome eQTL predictions are already strong enough to provide a nontrivial directional signal for many loci.

We propose SuSiNE, which adds a second channel through a non-zero prior mean ***µ***_0_ = *c****a***. Here ***a*** is a signed effect-size annotation vector and *c* is a scalar that places the annotation in effect-size units. The prior mean enters the single-effect Bayes factor jointly with the observed association direction. An annotation that agrees in sign with the observed effect reinforces the evidence on average, while a disagreeing annotation is attenuated. We refer to this behavior as “self-gating.”

Self-gating does not make arbitrary annotations harmless. Instead, their harm depends on S2F model error and the association noise not being strongly aligned, which is an empirical condition that can be measured diagnostically. We test the expected model behavior under independent noise and also test its failure when annotations for null variants are deliberately aligned with marginal association noise. SuSiNE preserves conjugate single-effect updates, credible-set construction, monotone ELBO ascent with stationary accumulation points under the stated conditions, and summary-statistic inference. The signed ***µ***_0_ channel and the unsigned ***π*** channel are complementary, and a single fit can use both.

The optimizer and model-specification barriers require a different response, and we address them with paired ensembling and aggregation. The ensemble fits a grid of SuSiNE models under perturbed initializations and/or priors so that it can explore different regions of the ELBO landscape. The aggregation step then combines the PIP summaries from the individual ensemble fits using cluster-aware ELBO weights. We use the ELBO as an internal ordering of fit quality rather than as an exact posterior model probability, and we return to that approximation in the Discussion. Exploration and summarization go hand-in-hand: exploration without aggregation still forces the analyst to choose one fit, while aggregation without exploration has no alternative basins to summarize.

We make three linked methodological contributions. First, SuSiNE supplies a signed prior-mean channel that allows directional functional annotations to inform fine-mapping while preserving their sign. Second, the paired ensemble and cluster-weight workflow explores the variational landscape and summarizes across the optima found. Third, we reconsider SuSiE-family diagnostics, showing why purity should not be a default filter on fitted model effects, and introducing effect-level measures of posterior concentration, accuracy, and fitted-basis movement.

We evaluate these contributions through several complementary analyses. Pathology examples, realistic oligogenic simulations, and a GTEx Lung eQTL case study are all used to measure and distinguish the three failure modes of the SuSiE and SuSiNE methods. We test whether purity filtering discards useful effects, and compare the utility of the fixed annotations through the signed ***µ***_0_ channel against annotation-free SuSiE variants, functional-***π*** priors, and empirical-Bayes SuSiNE. We test ablation methods over the ensemble and aggregation procedures as well, to assess the most computationally efficient methods for exploring and summarizing the variational landscape. We run sensitivity tests to evaluate the robustness of SuSiNE and SuSiE performance to alternative genetic architectures, and to deliberately pathological annotations aligned with association noise. Lastly, we run real-data diagnostics to assess the realistic risk of annotation confounding in the GTEx Lung panel. The open-source SuSiNE R package we developed provides both individual-level and summary-statistic interfaces.

## 2 Results

### 2.1 SuSiE pathology results

We designed four toy datasets with *n* = 600 and *p* = 50 to isolate the three barriers described in the Introduction. In every dataset, variants {1, 3, 5} were causal, variants {2, 4, 6} were matched decoys, and the other 44 variants were null. We fit SuSiE with *L* = 5 from its default initialization, a truth-warm start, and a decoy-warm start. Scenario 1 was an independent-variant control. Scenario 2 introduced causal–decoy pairs and a low-correlation pair with tied association evidence. Scenario 3 created distinct explanatory bases for nearly the same fitted signal. Scenario 4 made the decoys attractive in marginal updates even though they had much lower joint explanatory power. Fig 1 summarizes the results. Complete constructions and numerical values are given in S1 Appendix Section 1 and Table A.

**Fig 1.**
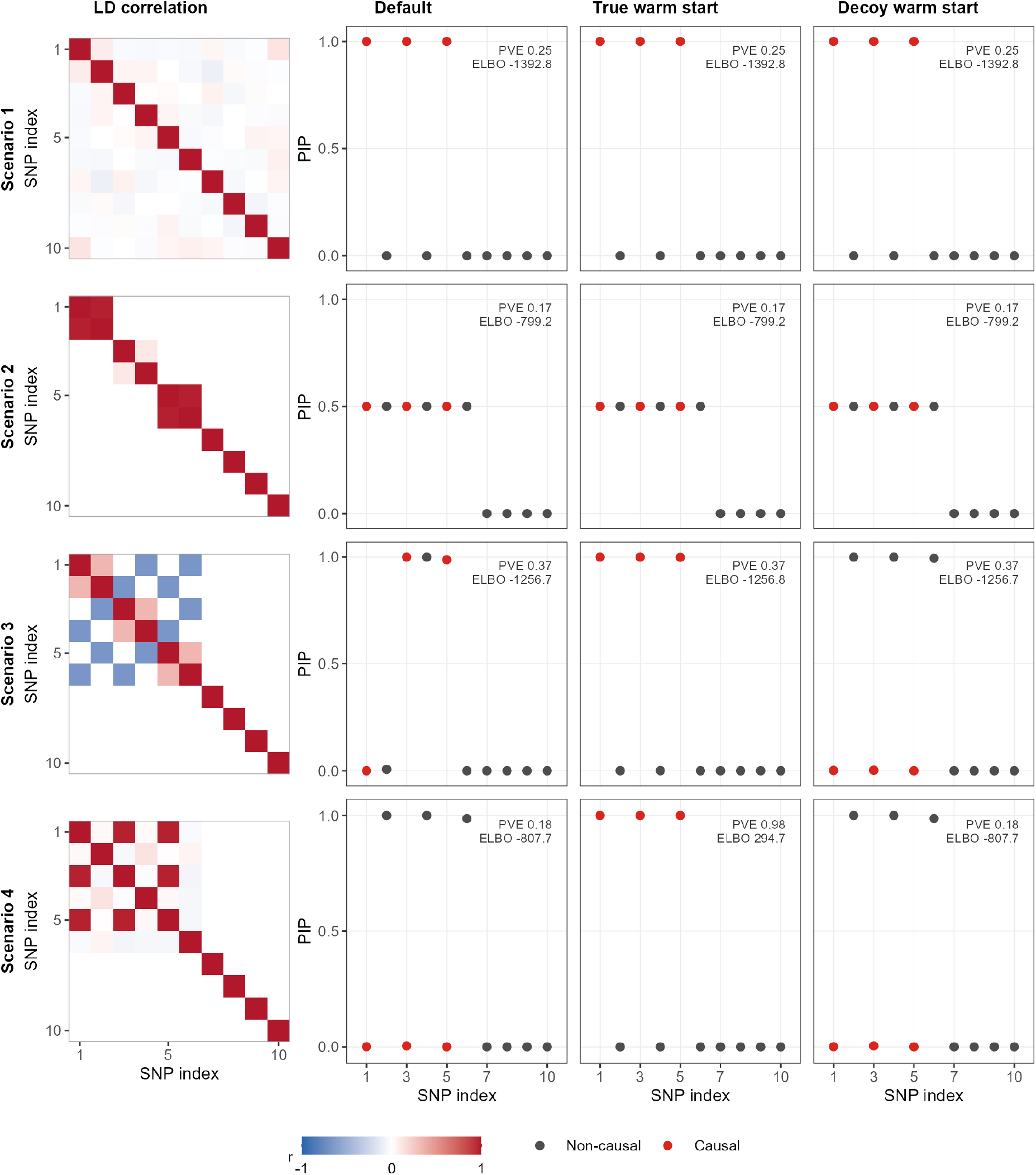
Toy examples isolate three distinct SuSiE failure modes. Each row is one scenario; columns show the genotype correlation matrix for the structured variants and PIPs from default, truth-warm, and decoy-warm initializations (*n* = 600, *p* = 50). Red points mark true causal variants {1, 3, 5}; decoys are {2, 4, 6}. Scenario 1 is a control. Scenario 2 shows honest LD ambiguity, including a low-purity but informative two-variant credible set at *r* = 0.10 whose removal lowers AUPRC. Scenario 3 shows a model-specification barrier: three near-equal-ELBO basins cannot be represented by one factorized fit. Scenario 4 shows an optimizer barrier: default IBSS remains in an inferior decoy basin, while the truth-warm fit reaches the high-PVE causal basin.

#### Scenario 1 (control)

All three initializations produced the same fit. PIP was approximately one on the three causal variants and approximately zero elsewhere. Every fit reached ELBO −1392.79 and fitted PVE about 0.25. This establishes the expected behavior when none of the three barriers binds.

#### Scenario 2 (LD ambiguity and purity)

All three starts again converged to the same fit at ELBO −799.21 and fitted PVE about 0.17. SuSiE split PIP (0.50, 0.50) within each causal–decoy pair. This is the behavior expected for an AND-of-ORs uncertainty pattern that the factorized posterior can represent. The middle pair is especially informative. Its variants have correlation *r* = 0.10 and marginal PVE about 6.6%, yet the realized association evidence does not distinguish them at the resolution of the fitted component. That component explains about 3.5% of phenotypic variance. Because the model uses *L* = 5, two components remain idle and could have represented additional effects if the data supported them.

The middle credible set has purity 0.10, far below the conventional 0.50 threshold, but it localizes a real high-effect signal to two variants. The unfiltered fit has causal-recovery AUPRC 0.50. Removing the two idle diffuse components leaves AUPRC unchanged at 0.50. Purity filtering instead removes the genuine middle component and lowers AUPRC to 0.35. In this example, purity describes the LD structure of the set but is not a safe automatic gate on the reported PIPs.

#### Scenario 3 (model-specification barrier)

The three initializations reached three different configurations. The truth-warm fit selected the causal set {1, 3, 5}, the decoy-warm fit selected {2, 4, 6}, and the default fit selected the mixed set {3, 4, 5}. All three configurations reproduce nearly the same fitted signal. Their ELBOs were −1256.82, −1256.70, and −1256.68, giving a total spread of only 0.14 nats and approximate ELBO-softmax weights (0.31, 0.34, 0.35). Each fit is confident in one configuration. No single factorized fit can retain the coordinated uncertainty across all three. Default empirical-Bayes prior-variance updating does not remove this limitation.

#### Scenario 4 (optimizer barrier)

Scenario 4 has a clearly dominant basin rather than a near tie. The default and decoy starts settle in the decoy basin with fitted PVE 0.18 at ELBO −807.72. The truth-warm start reaches the causal basin with fitted PVE 0.98 at ELBO 294.69. The gap exceeds 1,100 nats. The decoys win the first marginal screen, reaching |*r*(**y***, x_j_*)| as high as 0.34, while no causal variant exceeds 0.11. Greedy IBSS can therefore settle in a badly inferior basin even when a much better one exists. The truth-warm result is not deployable because it uses the causal configuration. It is an idealized sensitivity-to-initialization diagnostic.

#### From pathology to diagnostics

These examples motivate the comparisons used below. We ask whether ELBOs agree across starts, whether PIP summaries agree, whether aggregation differs from the best individual fit, and whether the fitted basis moves. In realistic simulations and the GTEx case study, nonzero values of these diagnostics show that the corresponding mechanisms are present. They should not be read as a literal additive decomposition of the observed performance gain.

### 2.2 SuSiNE theoretical results

We summarize the analytical properties of SuSiNE most relevant to practice. Each single-effect prior has mean ***µ***_0_ = *c****a***, while the conjugate Gaussian structure of SuSiE is retained. The directional prior therefore enters as an additive term in the Bayes factor. The resulting behavior is a trade-off: a self-gating mechanism limits the influence of miscalibrated annotations, while fundamental non-identifiability constrains how the annotation scale *c* can be set. Complete proofs and additional results are deferred to S1 Appendix.

#### Single-effect posterior

Let 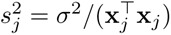 be the sampling variance of the OLS estimate *b̂_j_*, let 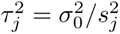 be the prior-to-noise variance ratio, and let 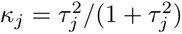 be the data shrinkage weight. Conditional on effect *ℓ* landing on variant *j*, the posterior on the effect size is 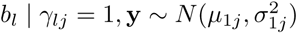 with

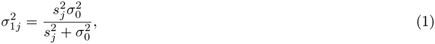

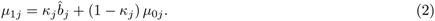

The posterior mean is a precision-weighted average of the OLS estimate and the prior mean: data dominate when *κ_j_* → 1, the annotation dominates when *κ_j_* → 0. Setting *µ*_0_*_j_* = 0 recovers the SuSiE update exactly. **Bayes-factor decomposition and self-gating.** Writing 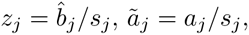 and 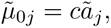 the log Bayes factor that drives the per-effect posterior inclusion weights decomposes as an approximate large-sample association Bayes factor [26]:

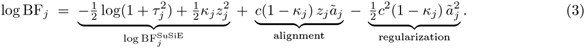

The first term is the standard SuSiE Bayes factor. The *alignment* term is positive when the marginal *z*-score and the annotation agree in sign, and negative when they disagree — this is the self-gating property previewed in Section 1.3: annotations that disagree with the data are automatically down-weighted, without any tuning. The *regularization* term is always negative, scaling with 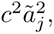, and provides built-in protection against unbounded inflation when *c* or |*ã_j_*| is large.

#### Properties inherited from SuSiE

Because SuSiNE only modifies the prior mean of an otherwise unchanged single-effect regression block, every SuSiE result that depends on conjugacy of the SER posterior or on the additive structure of the model carries over. Table 1 accounts for the seven properties most relevant to fine-mapping practice; each is verified in S1 Appendix with explicit reference to the original SuSiE statement (Wang et al. [15], propositions 1–3 and corollary 1; Zou et al. [16] for the RSS form). In particular, the summary-statistic extension follows because the signed prior mean changes the single-effect Bayes factor but does not change the RSS sufficient-statistic likelihood; S1 Appendix states this equivalence explicitly (Section 12, Remark 12.1). The practical significance is that SuSiNE preserves the structure and guarantees that make SuSiE attractive: the same per-effect tractability, monotone ELBO ascent with stationary accumulation points under the stated conditions, the same credible-set construction, and the same drop-in summary-statistic mode. It adds a second, signed channel for external information.

**Table 1.** Theoretical properties of SuSiE and their status in SuSiNE. Full statements and proofs are in S1 Appendix.

| SuSiE property | Status in SuSiNE | Why it matters for practice |
| --- | --- | --- |
| Variational factorization<br>$q = \prod_{\ell} q_{\ell}$ | Unchanged | Users still get per-effect PIPs and credible sets from a tractable posterior approximation. |
| Coordinate-ascent /<br>SER update | Unchanged | A SuSiNE update has the same order of computational cost as a SuSiE update, with the prior mean shifted by the annotation. |
| IBSS is coordinate<br>ascent on the ELBO | Unchanged | The fitting objective still improves monotonically under the usual exact IBSS updates. |
| Monotone ELBO ascent<br>and stationary<br>accumulation points | Verified for SuSiNE | The directional prior preserves the coordinate-ascent target under fixed positive variances and positive prior inclusion weights. |
| Exchangeability of<br>identical variants | Modified | Signed annotations can break ties among variants that the genotype data alone cannot distinguish, which is the intended use of the prior-mean channel. |
| Limiting equivalence to<br>BVSR as $p \rightarrow \infty$ | Unchanged | The directional prior changes how variants are scored, not the large-$p$ sparse-regression target inherited from SuSiE. |
| Summary-statistic<br>extension (RSS form) | Carries directly | Users can run the signed-prior workflow from GWAS or eQTL summary statistics and reference LD, which is essential when individual-level genotypes or expression data cannot be shared. |

#### Optimal annotation scale

The simplest non-trivial question is what value of *c* a practitioner ought to use. Under the stylized analysis in S1 Appendix, the expected log Bayes factor at a causal variant decomposes into a linear gain and a quadratic regularization cost. After centering the causal-variant effect and annotation distributions, the general maximizer is

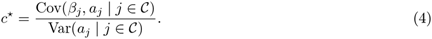

Thus *c^★^* is a regression slope and depends on the units of *a*. Only under the unit-loading generator *a_j_* = *β_j_* + *ɛ_j_* used in the supplement does this reduce to *c^★^* = *ϕ_a_*, where *ϕ_a_* = Corr(*a_j_, β_j_* | *j* ∈ C)^2^. Under the homogeneity assumptions of that analysis the maximizing slope is independent of the prior variance and sample size, although those quantities control the size of the gain. The equality *c^★^* = *ϕ_a_* should therefore be read as a property of the normalized generator, not as a scale-free recipe for an arbitrary annotation.

#### Non-identifiability of *ϕ_a_*

The target in Eq. 4 is conditional on the unknown causal set C. An all-variant statistic such as Corr(*a_j_, z_j_*) conflates causal signal, null noise, and the unknown causal fraction *ρ* = |C|*/p*. Without labeled causal variants, auxiliary restrictions on the causal and null distributions, or information pooled across loci, the causal-set regression slope is therefore not identified in general from one locus (S1 Appendix). This limits what a single fine-mapping problem can say about annotation scale; it does not imply that every estimator of *c* must fail, and comparisons of single-fit estimators with grid-based alternatives remain empirical.

#### Quadratic regret in *c*

A complementary result in S1 Appendix quantifies the cost of missing the optimum under the unit-loading generator. Writing 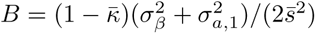 for its curvature constant, the expected-log-Bayes-factor regret is

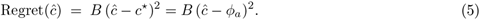

The objective is locally flat at the optimum and the penalty is symmetric in over- and under-estimation, so a coarse *c* grid is comparatively forgiving in the neighborhood of *ϕ_a_* — supporting the eight-point grid we use in the simulation and real-data ensembles. We note that absolute regret still scales with the locus-specific curvature constant *B*, which depends on unknown nuisance parameters; the claim is therefore relative (a coarse grid is acceptable in regret-per-unit-*B*), not absolute (regret is small in every regime).

#### Why ranking is harder than expected-ELBO improvement

We close by flagging a result that explains why positive expected gains in log Bayes factor at causal variants do not, in general, translate into proportional gains in AUPRC. The precision of any ranking is throttled by the causal fraction *ρ*. In particular, achieving precision *η* at recall *r* requires the null tail probability to satisfy *u*(*r*) ≤ *ρ*(1 − *η*)*/*((1 − *ρ*)*η*) · *r*, so in sparse regimes (*ρ* ≪ 1) even small null-side variance in the alignment term *z_j_ã_j_* can dominate the precision denominator. This suggests that performance varies with genetic architecture. Our oligogenic simulation results should therefore not be assumed to transfer unchanged to substantially sparser or more polygenic regimes. The sensitivity checks below support that caution: the C-CS gain remains positive under the tested sparse and diffuse architectures, but it is generally smaller than in the central oligogenic benchmark and high-precision recall can remain limited. The full statement and a Gaussian-shift corollary are in S1 Appendix.

### 2.3 Simulation study results — baseline

We benchmarked eight SuSiE-family specifications on 600 oligogenic datasets generated from 150 European-ancestry genotype matrices. Each dataset had *n* = 600, *p* ≈ 1000, 23 causal variants divided among strong, moderate, and weak effect tiers, and total heritability *h*^2^ = 0.25. We scored recovery of the eight largest-effect causal variants. Lower-effect causal variants were omitted from the precision–recall calculation rather than counted as negatives.

The annotation generator varied two different features which jointly determine annotation quality. The parameter *ϕ_a_* is the squared correlation between annotations and true effects at causal variants, and controls the accuracy of annotations at causal indices. The parameter *ν_a_* is the ratio of null to causal annotation variance, and controls the relative scale of the annotations at noncausal indices, which are by definition fully noised. Annotation-consuming specifications were evaluated across the full 3 × 3 (*ϕ_a_, ν_a_*) grid. The point (0.3, 0.9) was the nearest coarse-grid match to published AlphaGenome eQTL benchmarks. Table 2 summarizes the specifications. Fig 2 reports pooled AUPRC, and Fig 3 reports the baseline SuSiE purity-filter diagnostic.

**Fig 2.**
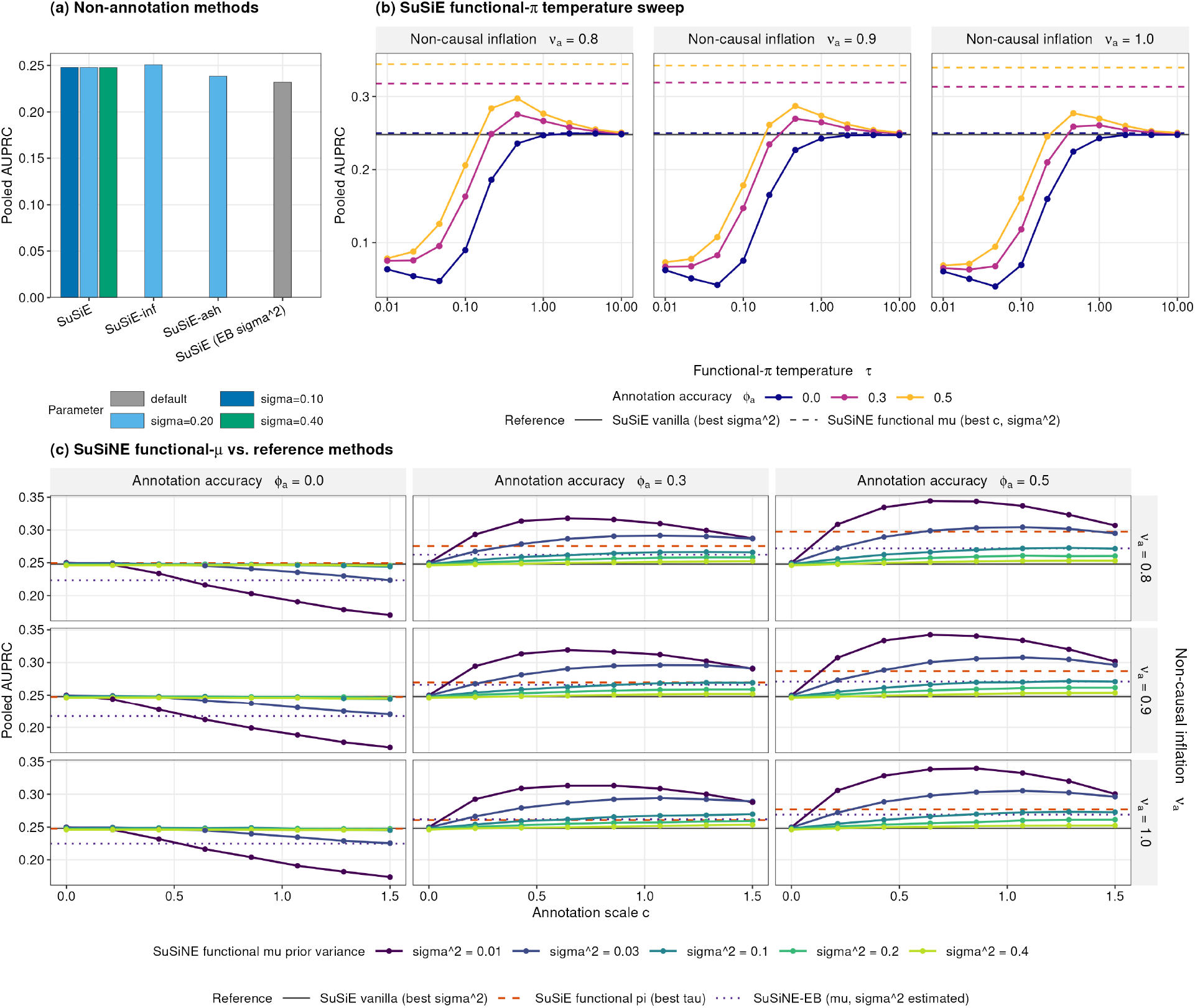
Signed prior means improve fine-mapping under calibrated annotations. Pooled AUPRC is shown across baseline single-fit specifications on the 600 oligogenic-architecture simulations. **(a)** Annotation-agnostic methods at their fixed-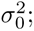 settings; SuSiE-inf is marginally strongest, but the spread within this family is small. **(b)** SuSiE-functional-*π* across the temperature grid *τ* on a log_10_ axis, faceted by *ν_a_* and colored by *ϕ_a_*. The solid black reference is SuSiE-vanilla at its best 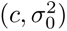; the colored dashed lines are *SuSiNE*-functional-*µ* at its best 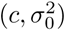 within each *ϕ_a_* stratum. Functional-*π* approaches the *SuSiNE*-functional-*µ* ceiling only at intermediate *τ* and degrades sharply at small *τ* when annotations are uninformative (*ϕ_a_* = 0), reflecting the absence of a self-gating property in the ***π*** channel. **(c)** *SuSiNE*-functional-*µ* AUPRC across the 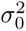 grid, faceted on *ϕ_a_* (columns) and *ν_a_* (rows). Reference lines: SuSiE-vanilla at best 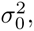 (solid grey), SuSiE-functional-*π* at best *τ* (dashed orange), *SuSiNE*-EB (dotted purple). The annotation regime (*ϕ_a_, ν_a_*) = (0.3, 0.9) is the named baseline-grid working point nearest the AlphaGenome calibration in S1 Appendix Fig A.

**Fig 3.**
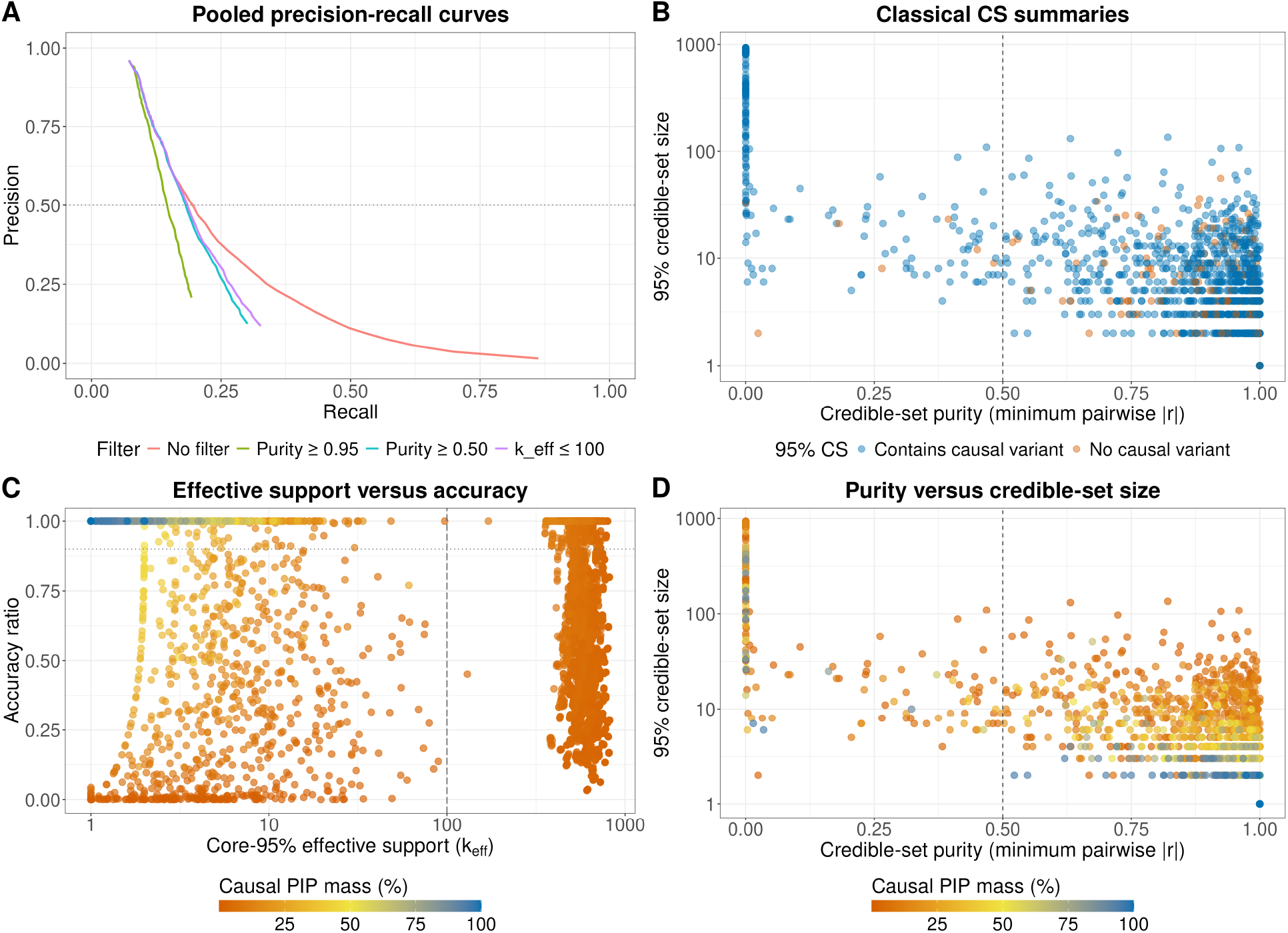
Purity filtering discards informative fine-mapping signal. Diagnostics are shown for the annotation-agnostic SuSiE baseline on the 600 oligogenic simulations. **(A)** Pooled precision–recall curves after recomputing PIPs from effects retained by each rule; all filters reduce AUPRC relative to the unfiltered fit (0.248). **(B)** Credible-set purity versus set size, colored by whether the set contains a causal variant; the dashed line marks the conventional purity-0.50 cutoff. **(C)** Core-95% effective support *K_ℓ_* versus effect accuracy ratio *A_ℓ_*, colored by the percentage of PIP mass on causal variants; the dashed vertical line marks *K_ℓ_* = 100. **(D)** Purity versus credible-set size, colored by causal PIP mass; the dashed line again marks purity 0.50. Panels B–D are simulation diagnostics because causal coverage, causal PIP mass, and *A_ℓ_* require known truth.

**Table 2.**
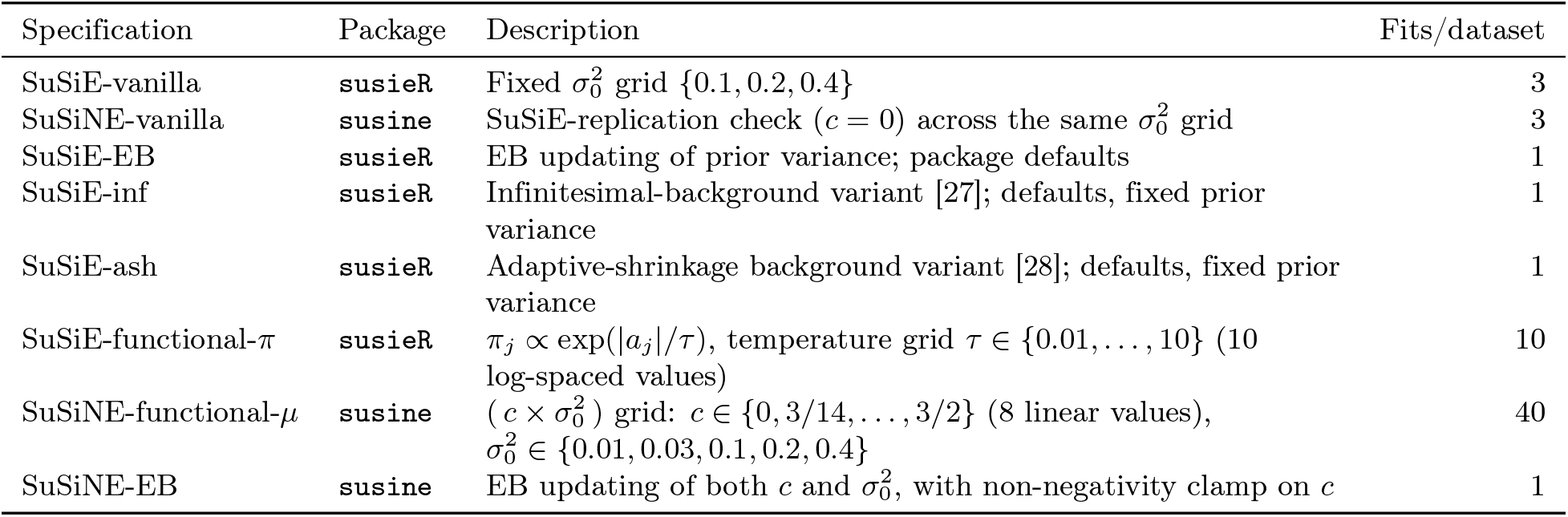
Baseline model specifications. “Fits/dataset” is per (dataset, annotation-setting) combination for annotation-consuming specifications and per dataset for annotation-agnostic specifications.

#### Annotation-agnostic baselines

Among the four annotation-agnostic specifications (SuSiE-vanilla at fixed 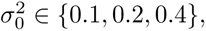 SuSiE-EB, SuSiE-inf, SuSiE-ash), pooled AUPRCs are relatively close but not identical (Fig 2a). SuSiE-inf is marginally strongest (AUPRC = 0.251), while SuSiE-vanilla is nearly indifferent to the fixed prior variance over the reasonable range tested (best AUPRC = 0.248). SuSiE-ash and SuSiE-EB trail at 0.238 and 0.232, respectively. The total spread across this family is 0.019 AUPRC units, which is small compared with the functional-*µ* gains below but not negligible in absolute terms. Under the oligogenic regime studied here, changing the annotation-agnostic SuSiE variant yields limited improvement, with SuSiE-inf helping slightly and SuSiE-ash and SuSiE-EB hurting modestly.

#### Annotation-consuming methods at the AlphaGenome calibration point

At the working calibration point (*ϕ_a_, ν_a_*) = (0.3, 0.9), *SuSiNE*-functional-*µ* at its best 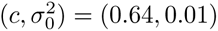 achieved AUPRC = 0.319. This exceeded the strongest annotation-agnostic baseline (SuSiE-inf) by 0.068 AUPRC units. SuSiE-functional-*π*, the closest existing method for incorporating functional annotations through SuSiE’s prior-inclusion channel, peaked at AUPRC = 0.270 at *τ* ≈ 0.46, recovering only 0.019 of the 0.068 AUPRC gain delivered by the ***µ***_0_ channel (Fig 2b). The empirical-Bayes specification *SuSiNE*-EB, which jointly estimates *c* and 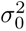 without any user grid search, achieved AUPRC = 0.266, trailing the grid-searched *SuSiNE*-functional-*µ* by 0.053 and landing near the functional-*π* operating point. This makes it a useful no-grid diagnostic, but not a substitute for the fixed-grid ***µ***_0_ workflow carried forward below.

#### Sensitivity across annotation quality

Both annotation channels are double-edged when annotations are poorly tuned or uninformative (Fig 2b–c). The relevant baseline-screen question is therefore not whether a single annotated fit is uniformly robust, but how much signal can be extracted from annotations of plausible quality and how sensitive that extraction is to *τ*, *c*, and 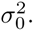 Under the central calibration point and the higher-quality *ϕ_a_* = 0.5 regimes, the best tuned ***µ***_0_ model clearly exceeds the best tuned functional-*π* model. The advantage is also not confined to one exact value of *c*: the ***µ***_0_ heatmap shows a broad high-performing region around the selected scale and small prior variance. Under null or low-quality annotations, both channels contain settings that limit damage, including *c* = 0 for ***µ***_0_ and weak functional-*π* weighting. But badly tuned annotations can still hurt. We therefore treat single-fit grids as a measurement of the achievable gain from functional annotations, while the ensemble analysis below is where we ask whether grid aggregation can make the workflow more tolerant of unknown annotation quality.

#### Credible-set purity filtering

We used two effect-level diagnostics to interpret these comparisons. The effective top-95% support size *K_ℓ_* measures diffuseness as the entropy-equivalent number of variants in the top 95% of an effect’s PIP mass. The accuracy ratio *A_ℓ_* compares the largest causal-variant PIP with the largest PIP in the effect. It equals one when a causal variant is top-ranked. Because *A_ℓ_* uses the known causal set, it is a simulation diagnostic.

The baseline SuSiE purity diagnostic confirms that the toy pathology in Section 2.1 also appears in realistic oligogenic simulations (Fig 3). Recomputing combined PIPs after dropping effects, the unfiltered baseline achieved pooled AUPRC 0.248. The conventional purity-0.50 filter reduced AUPRC to 0.189, a loss of 0.059. Stricter purity filters lost more signal, with AUPRC 0.164 at purity 0.90 and 0.147 at purity 0.95. A direct diffuseness filter, retaining only effects with *K_ℓ_* ≤ 100, also reduced AUPRC to 0.196. Thus every hard filter tested was below the unfiltered analysis (Fig 3A). The unfiltered precision-recall curve dominates the filtered curves over the shared recall range, rather than trading lower recall for higher precision. The separation is already visible near the 50% precision region and widens through the 25% precision region, where a method would still be useful as a variant-prioritization screen.

One possible concern is that this result could be an artifact of a rank-based metric. If filtering only removes very low-PIP tail variants, then unfiltered PIPs might score better by ranking irrelevant tail mass without improving the part of the posterior that practitioners use. We therefore examined per-effect causal PIP mass, the total posterior mass that an effect assigns to true causal variants. This response does not reward a metric for ordering tiny PIP values well. It asks whether the effect places meaningful posterior mass on causal variants. Across 6,000 active effects, causal PIP mass was right-skewed, with median 0.051 and mean 0.159.

Panel B asks whether purity separates causal from non-causal effects once effect accuracy is shown directly. It does not. Non-causal effects are not confined to the low-purity region; many appear above the conventional purity threshold, while many low-purity effects still cover a causal variant. Thus purity is not a reliable quality score even when the plotted response is a truth-aware diagnostic.

Panel C compares 95% credible-set size with the core-95% effective support *K_ℓ_*. The two are related, but not interchangeable: a large credible set can still have a concentrated posterior core, while a smaller set can remain diffuse across its members. This is why *K_ℓ_* is a cleaner concentration measure than raw credible-set size.

Panel D directly compares purity with *K_ℓ_*. Apart from a diffuse low-purity band, low purity is not simply a proxy for posterior diffuseness. The count summaries make the point sharper: among 4,428 effects below purity 0.50, only 0.2% cover no causal variant, whereas among 1,572 effects at or above purity 0.50, 10.1% cover no causal variant. The standard purity gate therefore removes many low-purity effects that still carry causal signal. We therefore retain purity as an LD summary, but do not use it as a default gate on reported PIPs.

#### Effect drift in the high-quality regime

In the high-quality annotation regime (*ϕ_a_, ν_a_*) = (0.5, 0.9) we used a 5-arm refit chain to separate how the ***µ***_0_ and ***π*** channels reshape the fit. Each cold annotated fit was followed by an annotation-free vanilla refit initialized from its solution. This warm refit tested whether an annotation-discovered basin remained stable after the annotation was removed. Pooled precision-recall improved most for annotated ***µ***_0_. Its AUPRC rose from 0.25 for cold SuSiE to 0.35, compared with 0.29 for annotated ***π***. The gain also fell in a practically actionable part of the curve. At a fixed 50% precision, the ***µ***_0_ channel recovered 64% more top-eight causal variants than cold SuSiE (31% versus 19%). Decomposing the gain on the per-effect (*K_ℓ_, A_ℓ_*) plane and against the truth-basis-divergence metric shows that annotations act through two mechanisms. The first is a prior-dependent within-basin sharpening that re-centers and concentrates effects, but this sharpening is largely erased when the prior is removed by warm refit. The second is a smaller but durable between-basin movement toward the true causal basis, and this movement survives prior removal. The ***µ***_0_ channel is more effective than ***π*** at both. The durable between-basin component is the realistic-locus analogue of the optimizer barrier of Section 2.1, and it foreshadows the real-data results of Section 2.6. Full methods, pooled precision-recall curves, per-effect density shifts, and the truth-basis-divergence decomposition are reported in S1 Appendix (Section 13.1, Figs B–C).

#### Decision for the ensemble study

On these baseline benchmarks the ***µ***_0_ channel dominates the ***π*** channel under matched signed annotations, with a wider operating window in 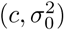 than functional-*π* has in *τ*. We therefore carry *SuSiNE*-functional-*µ*, both grid-searched and EB-updated, into the ensemble study below. We discontinue the SuSiE-ash, SuSiE-inf, and functional-*π* specifications from further evaluation.

### 2.4 Simulation study results — ensemble

We evaluated ensembles that varied initialization, refinement, prior-mean scale, and prior variance. Ensemble labels record the model family and the exploration axes. Prefix C denotes fixed-variance functional-*µ* fits, while suffixes C and S denote grids over annotation scale and prior variance. The primary C-CS ensemble crossed eight values of each parameter, giving 64 fits per dataset. C-CSR began with those same fits and then removed the annotation prior in a warm refit. It therefore measures whether annotation-discovered basins remain stable under the annotation-free objective.

We compared four deployable summaries of the same fits: max-ELBO selection, uniform averaging, ELBO-softmax averaging, and cluster-weight aggregation. Cluster weighting first merges near-duplicate PIP summaries and then weights the resulting clusters by the best ELBO in each cluster. This prevents a frequently reached basin from gaining weight only because it was found many times. We also report a truth-aware best-member benchmark. It uses causal status and is not deployable.

At the primary annotation setting (*ϕ_a_, ν_a_*) = (0.3, 0.95), C-CS with cluster-weight aggregation was the strongest deployable workflow (Fig 4). Relative to the zero-prior-mean SuSiE-equivalent baseline (AUPRC 0.2474), it reached AUPRC 0.3130. This improved pooled top-eight causal recovery by an absolute ΔAUPRC of 0.0656 (95% paired-bootstrap CI [0.0591, 0.0722], or 26.5% relative improvement). This annotation-agnostic SuSiE baseline appears as 0.2474 here (ensemble study, pooled confusion bins, (*ϕ_a_, ν_a_*) = (0.3, 0.95)), as 0.248 in the baseline screen (Section 2.3, same pooled-bin convention but the (0.3, 0.9) grid point), and as ≈ 0.25 in the drift chain (S1 Appendix Fig B, continuous-threshold integration). These differ only by annotation-grid point and AUPRC computation convention, not by underlying data. An idealized truth-warm initialization benchmark within vanilla SuSiE reached AUPRC 0.2851, a +0.0376 gain computed from unrounded results. Because it requires the true configuration, this is a sensitivity-to-initialization benchmark rather than a deployable exploration gain. The C-CS max-ELBO fit reached AUPRC 0.3090, another +0.0239 above truth-warm vanilla. This is a net contrast, not a pure functional-prior estimand. Cluster-weight aggregation added +0.0040 beyond max-ELBO, giving the headline 0.3130. The C-CS truth-aware best-member benchmark reached AUPRC 0.3364 (ΔAUPRC = 0.0890, 95% CI [0.0821, 0.0962]). The observed 0.0234 gap from cluster weighting to this benchmark measures headroom relative to choosing one member with ground truth, rather than a true bound on an optimal aggregation rule. C-CSR with cluster-weight aggregation reached AUPRC 0.2568, only +0.0094 over baseline (+3.8% relative to baseline and 14.3% of the headline absolute gain), showing that most of the cold-fit annotation gain was prior-dependent. C-CSR remains an idealized durability diagnostic because candidate basins were discovered under annotated priors before the prior was removed. Multimodality diagnostics show that C-CS explores more distinct PIP summaries than the annotation-agnostic and empirical-Bayes refinement ensembles (S1 Appendix, Fig D). Table 3 summarizes this diagnostic ladder without treating its non-nested contrasts as an additive causal decomposition.

**Fig 4.**
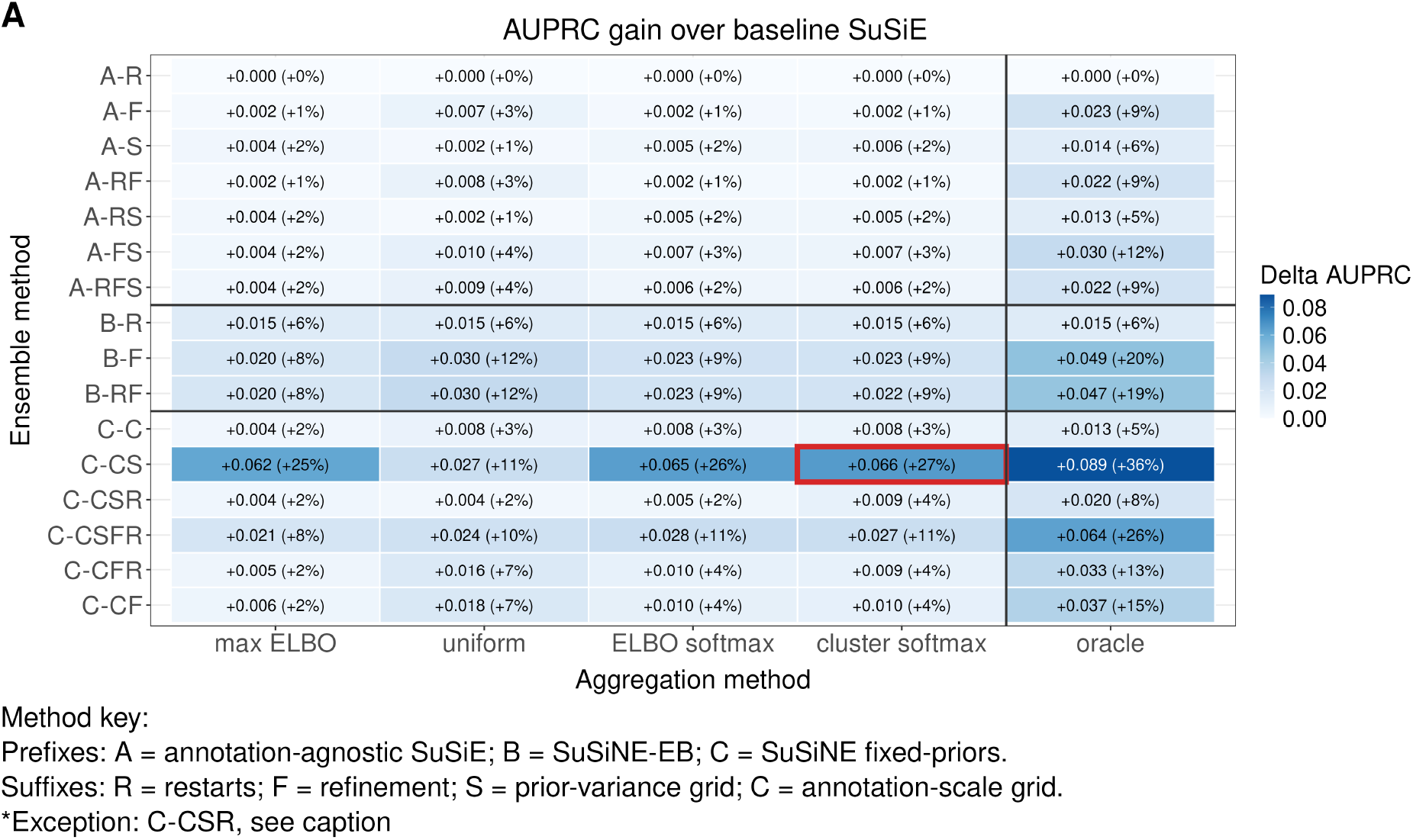
Cluster-weight aggregation identifies the strongest deployable ensemble. Pooled ΔAUPRC is shown versus the zero-prior-mean SuSiE-equivalent baseline for top-eight causal recovery on the 600 phenotype datasets from 150 genotype matrices at (*ϕ_a_, ν_a_*) = (0.3, 0.95). Rows are ensemble strategies grouped by model family (A: annotation-agnostic; B: EB functional-*µ*; C: fixed-variance functional-*µ*) and suffixes R, F, S, and C for restarts, refinement, prior-variance grids, and annotation-scale grids. Columns are aggregation rules. “Cluster softmax” is the primary cluster-weight rule: complete-linkage clustering of PIP vectors under the credible-shift distance at cut 0.05, cluster weights proportional to each cluster’s maximum ELBO, and within-cluster weights split by ELBO-softmax. C-CSR reruns C-CS source fits under the default zero-prior model to test durability after prior removal. The outlined C-CS cluster-softmax cell is the primary deployable workflow.

**Table 3.** Diagnostic ladder for ensemble AUPRC at the AlphaGenome calibration point (*ϕ*_a_*, ν*_a_) = (0.3, 0.95). The contrasts are not an additive causal decomposition because the interventions change both search and prior information. Deployable rows use no ground truth and select no single grid cell; the truth-warm and best-member rows are references that require ground truth. The separate single-fit *ceiling* comparison of the *µ*_0_ versus *π* channels is reported in Section 2.3.

| Configuration | AUPRC | $\Delta$ vs. baseline | Role |
| --- | --- | --- | --- |
| SuSiE-equivalent baseline | 0.2474 | — | deployable reference |
| Truth-warm vanilla SuSiE | 0.2851 | +0.0376 | initialization-sensitivity benchmark (not deployable) |
| C-CS, max-ELBO | 0.3090 | +0.0616 | deployable: exploration + functional prior |
| C-CS, cluster-weight | 0.3130 | +0.0656 | deployable: + aggregation (**recommended**) |
| C-CSR (prior removed) | 0.2568 | +0.0094 | durable exploration gain after annotation removal |
| C-CS best-member benchmark | 0.3364 | +0.0890 | truth-aware single-member selection (not deployable) |

Annotation accuracy controlled the size of the ensemble gain, but the response was asymmetric and nonlinear (Fig 5A–C). With uninformative annotations (*ϕ_a_* = 0, *ν_a_* = 0.95), C-CS cluster-weight aggregation was close to baseline but slightly lower (AUPRC 0.2415, −0.0060 or −2.4%). At the central setting (*ϕ_a_, ν_a_*) = (0.3, 0.95), the same workflow reached AUPRC 0.3130, and at *ϕ_a_* = 0.5 it rose to 0.3316 (+34.0% over baseline). At 75% precision, baseline SuSiE recovered 572 of 4800 target causal variants (recall 0.11917), whereas C-CS recovered 543 at *ϕ_a_* = 0 (recall 0.11313), 925 at *ϕ_a_* = 0.3 (recall 0.19271), and 1049 at *ϕ_a_* = 0.5 (recall 0.21854). The central-setting relative recall gain is 61.7% from the unrounded counts. C-CS PIPs were also better calibrated in high-PIP bins (Fig 5B), while estimated local genetic variance was modestly more conservative than baseline (panel D). The full 64-run grid took 29.83 seconds per dataset on average, or 62.5× the 0.477-second baseline fit. The compute-cost 4 × 4 subgrid, *c* ∈ {0, 0.5, 1.0, 1.5} crossed with 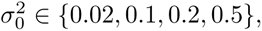 reached AUPRC 0.3137 in 7.41 seconds (15.5× baseline). Its slight advantage over the full grid is an empirical result, not a guarantee that fewer runs dominate.

**Fig 5.**
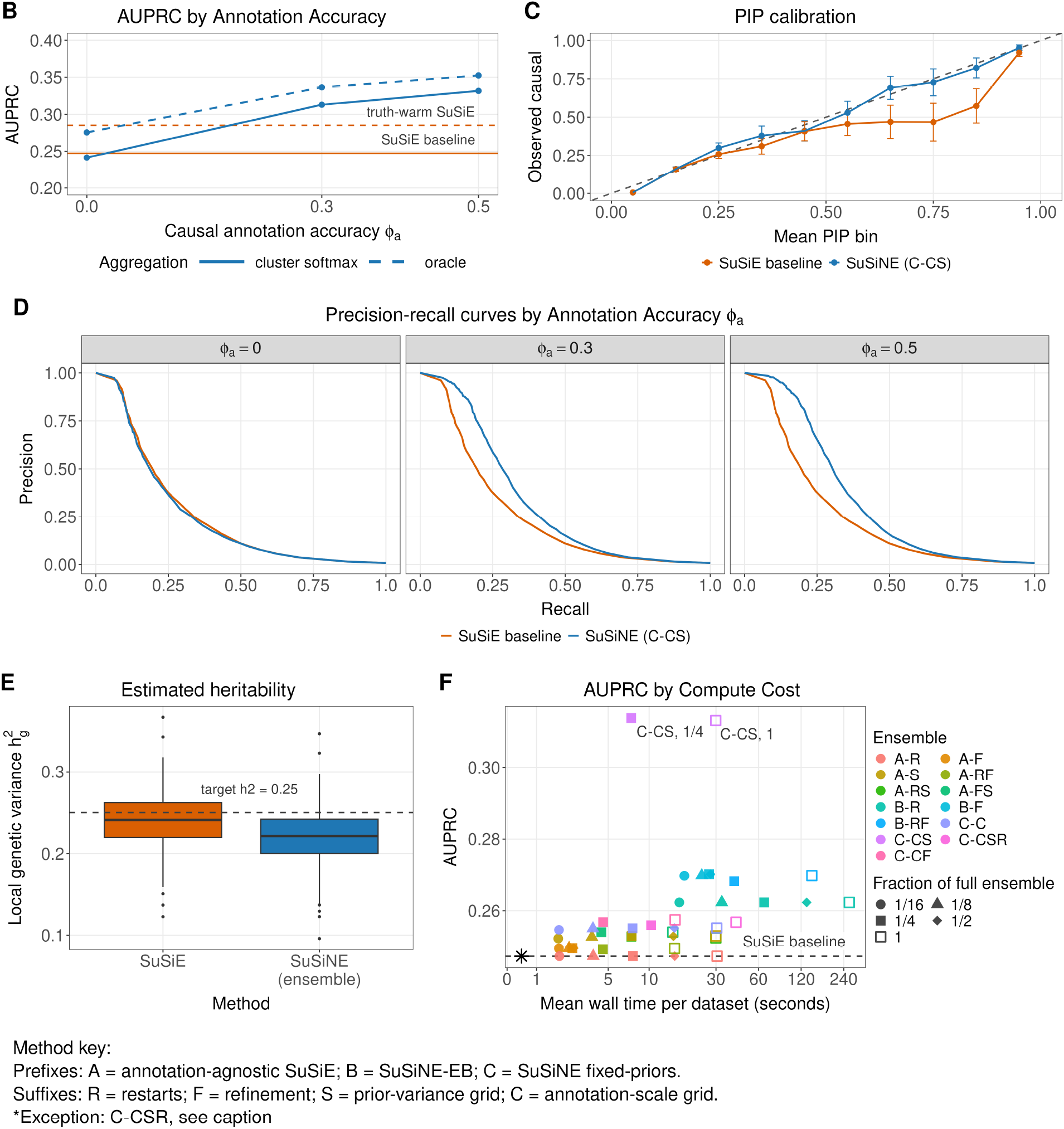
C-CS improves recovery, calibration, and computational trade-offs. The highlighted C-CS workflow is an 8 × 8 signed functional-*µ* grid over annotation scale *c* and prior variance 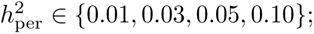 summarized by the credible-shift cluster-weight rule from Fig 4. **(A)** AUPRC across annotation accuracies *ϕ_a_* ∈ {0, 0.3, 0.5} at *ν_a_* = 0.95. **(B)** PIP calibration. **(C)** Precision–recall for top-eight causal recovery. **(D)** Posterior expected local genetic variance, E[var(**X*β***) | *y*]*/*var(*y*), with the design target *h*^2^ = 0.25 dashed. **(E)** AUPRC versus mean wall time per dataset for C-CS subgrids and the full grid.

### 2.5 Sensitivity check results

We first tested the recommended cluster-weight C-CS workflow under sparse and diffuse architectures using a smaller 4 × 4 parameter grid. The sparse arm used one, three, or five causal variants across four per-causal heritability levels. The diffuse arm used 100 causal variants across three levels of total local heritability. As in the central benchmark, the diffuse analysis scored recovery of the eight largest effects.

The architecture sensitivity checks support the main conclusion, but with smaller gains than in the central oligogenic benchmark (Fig 6). In the sparse arm, C-CS improved pooled AUPRC over the matched annotation-free baseline in all 12 settings. Absolute gains ranged from +0.0047 to +0.0500 AUPRC, with relative gains from 2.5% to 18.2%. The gain was largest at intermediate signal strengths and smallest when the sparse effects were either extremely weak, where both methods had little power, or very strong, where baseline SuSiE was already close to the attainable ranking. Recall at 75% precision followed the same pattern less uniformly: it was essentially unchanged in the weakest sparse settings, but increased by as much as 0.064 in the stronger settings. Thus sparse architectures throttle the available improvement, as predicted by the ranking theory, but the signed-prior ensemble did not reverse the baseline ordering.

**Fig 6.**
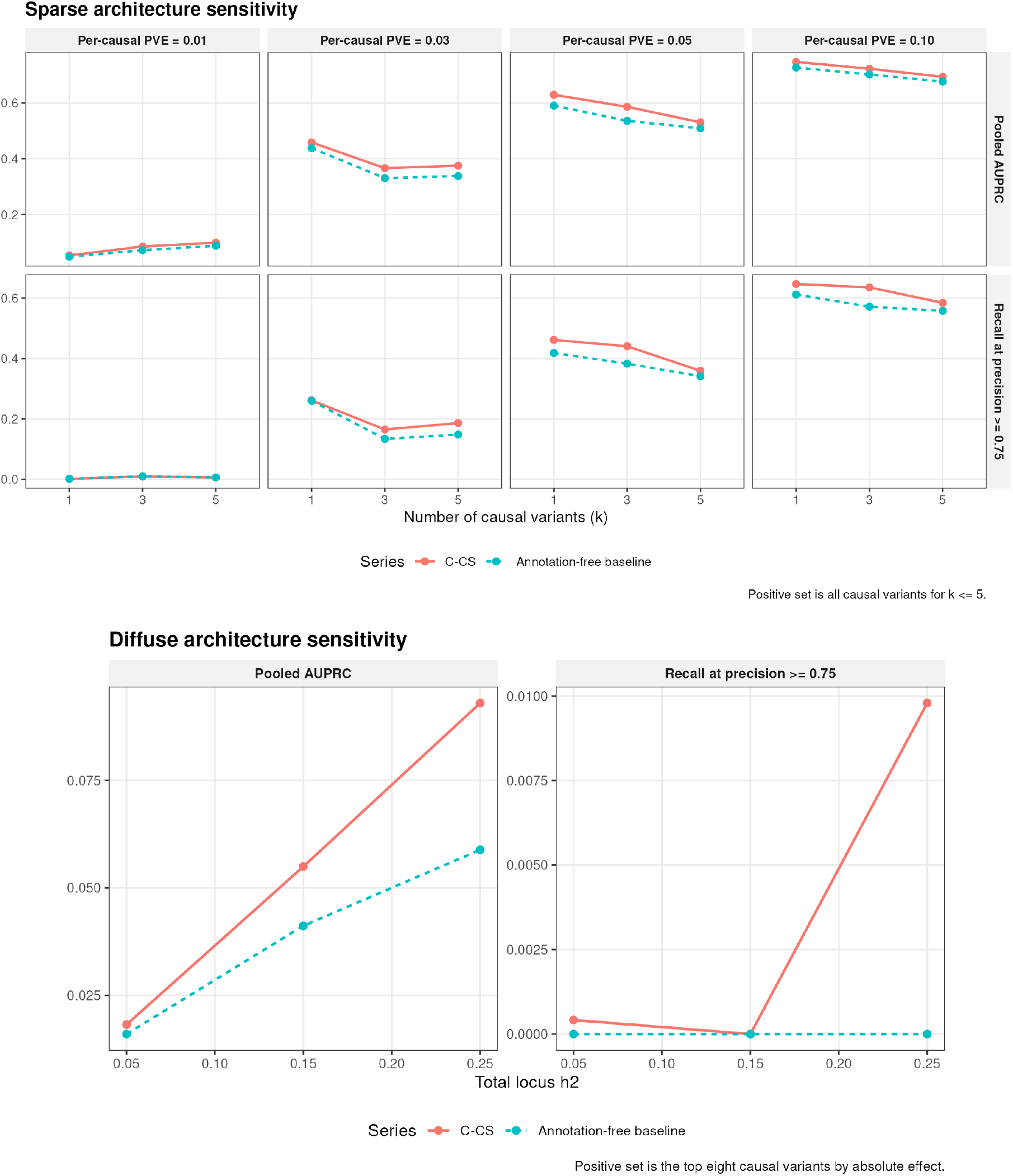
C-CS gains persist across sparse and diffuse genetic architectures. The recommended workflow is evaluated on 600 datasets per setting and compared with a matched annotation-free SuSiE-equivalent baseline. **(A)** Sparse architectures with *p^★^* ∈ {1, 3, 5} causal variants and per-causal heritability 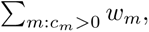 all causal variants are positives. **(B)** Diffuse architectures with 100 causal variants and total local heritability *h*^2^ ∈ {0.05, 0.15, 0.25}; positives are the top eight causal variants by absolute effect. C-CS improves pooled AUPRC in every architecture setting, but the absolute high-precision gain is smaller than in the central oligogenic benchmark and can be limited when signal is weak or very diffuse.

The diffuse arm is a harder high-precision recovery task because the data contain 100 causal variants and the metric scores only the top eight by absolute effect. C-CS again improved pooled AUPRC at all three total-heritability settings: 0.0182 versus 0.0160 at *h*^2^ = 0.05, 0.0550 versus 0.0412 at *h*^2^ = 0.15, and 0.0930 versus 0.0588 at *h*^2^ = 0.25. The relative gains increased with signal strength, from 13.9% to 58.0%, but absolute AUPRC remained low and recall at 75% precision remained under 1% even at *h*^2^ = 0.25. This is a useful boundary case: the annotation channel improves ranking, but a diffuse weak-effect architecture can still leave few discoveries at a stringent precision threshold.

We next blended the simulated annotation with marginal *z*-scores to test a failure of self-gating. The mixing weight *λ* ranged from zero to one. One arm changed annotations only at causal variants and served as a positive control. The other changed annotations only at null variants and tested the effect of aligning annotation error with association noise.

The annotation-contamination results show where the self-gating argument breaks (Fig 7). At *λ* = 0, the 4 × 4 C-CS sensitivity grid reached AUPRC 0.3159 and recall 0.2023 at 75% precision, compared with the annotation-free baseline AUPRC 0.2474 and recall 0.1192. In the causal positive-control arm, increasing *λ* strengthened alignment between annotations and marginal associations only at true causal variants. AUPRC improved through *λ* = 0.7 and then leveled off, reaching 0.3722 at *λ* = 0.7 and 0.3702 at *λ* = 1, while recall continued to rise to 0.2571. This confirms that the pipeline and prior channel respond in the expected direction when the added alignment is real causal signal.

**Fig 7.**
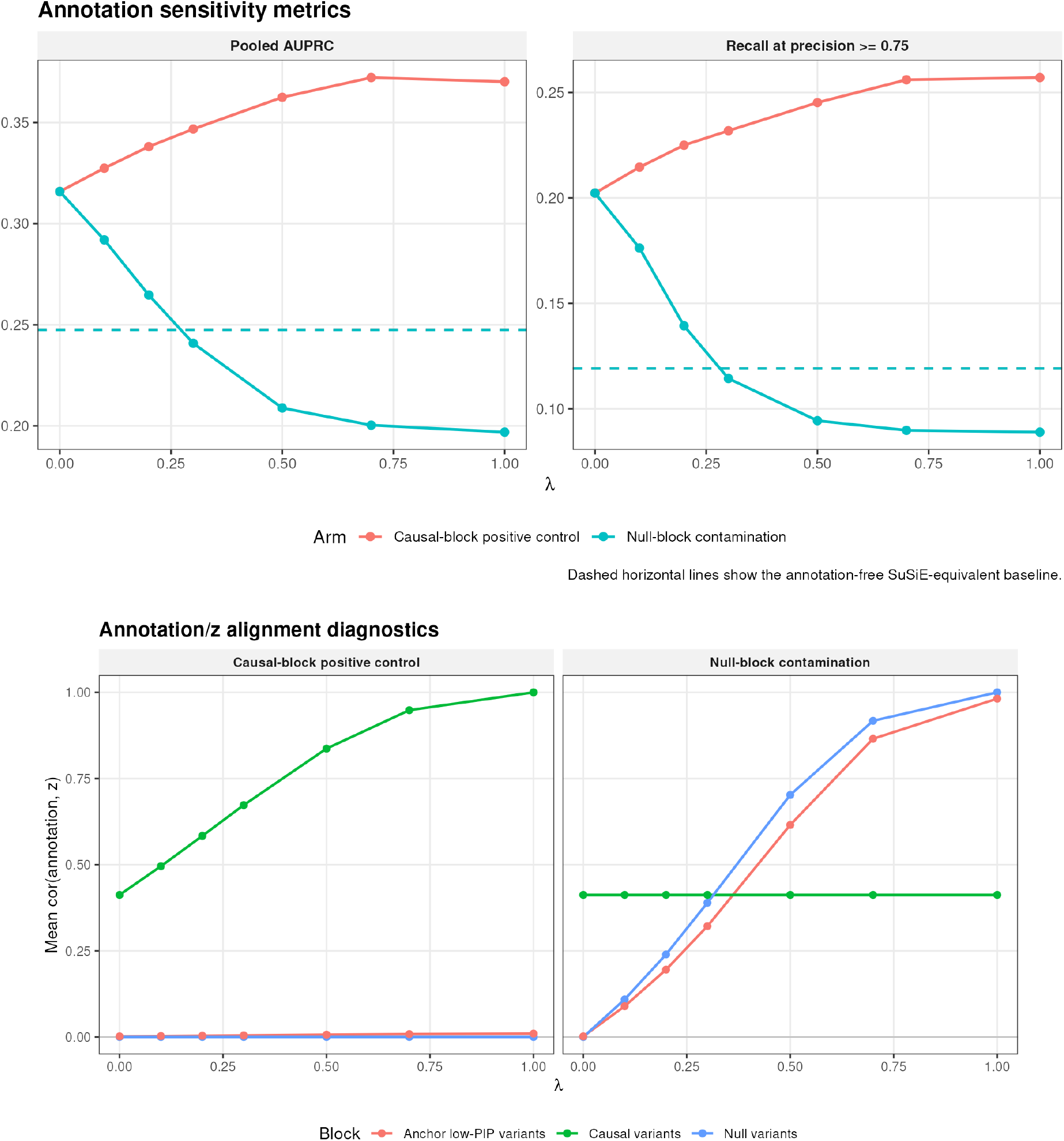
Strong null alignment can reverse annotation-informed gains. Sensitivity of the recommended C-CS workflow is shown on the central oligogenic simulator. **(A)** Pooled AUPRC and recall at precision ≥ 0.75 as the annotation vector is blended with marginal *z*-scores on either true causal variants (positive-control arm) or true null variants (null-contamination arm). Dashed horizontal lines mark the annotation-free baseline. **(B)** Realized correlations between annotations and marginal *z*-scores in the causal block, null block, and low-anchor-PIP variants. Causal-block alignment improves performance, while strong null-block alignment erases the C-CS gain.

The null-contamination arm is the stress test. When only non-causal annotations are blended toward marginal *z*-scores, C-CS remains above the annotation-free baseline at *λ* = 0.1 and 0.2 (AUPRC 0.2920 and 0.2647), but falls below baseline by *λ* = 0.3 (AUPRC 0.2409) and degrades further at stronger contamination (AUPRC 0.1969 at *λ* = 1). The corresponding diagnostic correlations show that this degradation occurs only after substantial alignment is introduced into the null block. The real-data internal null-alignment diagnostic is much smaller: among anchor-low-PIP variants in the 20 GTEx Lung loci, the median annotation–*z* correlation was 0.0124, the median absolute correlation was 0.0174, and the largest absolute locus value was 0.0807 (Fig F in S1 Appendix). These values sit below the anchor-low-PIP correlation seen at the mildest simulated null-contamination setting (*λ* = 0.1, correlation ≈ 0.089), where C-CS still retained a clear AUPRC gain. The result does not prove that AlphaGenome errors and association errors are independent in all real settings, but it argues against strong genome-local null contamination in this case-study panel.

This stress test also provides direct evidence for a future annotation-leakage concern. If sequence-to-function models are trained or calibrated on SuSiE-derived labels and their predictions later inform SuSiNE, their errors can become aligned with the association-derived evidence. The null-contamination arm establishes that this alignment can erase the benefit of the signed prior and eventually make performance worse than annotation-free SuSiE. It does not show that the current AlphaGenome annotations have this problem. It establishes the failure condition that a future circular training and fine-mapping pipeline would need to avoid.

### 2.6 Real data study results

We applied SuSiNE to GTEx v10 Lung eQTL summary statistics using 1000 Genomes EUR reference LD. The panel contained 20 protein-coding gene loci selected before AlphaGenome scoring to span a wide range of baseline fine-mapping difficulty. Each locus received signed AlphaGenome annotations. We fit the same 8 × 8 C-CS grid used in simulation and combined its 64 PIP vectors by cluster-weight aggregation.

We compared the ensemble with a cold annotation-agnostic SuSiE anchor and with an annotation-free warm refit. The warm refit began from the highest-weight ensemble member, which we call the “source run.” The distance *D*_ens_ is the PIP-vector L2 distance from the ensemble to the anchor. The analogous source and warm-refit distances are *D*_src_ and *D*_warm_. The matched-component drift *D*_MC_ aligns single-effect components and measures their movement under the reference LD matrix, weighted by the amount of fitted signal. It can reveal effect reallocation that cancels in the total fitted predictor.

Fig 8 summarizes four diagnostics across the panel, and Table 4 gives the locus-level values. A six-locus variant-level zoom is provided in S1 Appendix (Fig G). Across the panel, the SuSiE anchor assigned PIP *>* 0.5 to 17 variant–locus entries and the SuSiNE ensemble assigned PIP *>* 0.5 to 19. Eleven high-PIP entries overlapped. The ensemble therefore changed which variants were highlighted more than it changed the total number of high-PIP calls.

**Fig 8.**
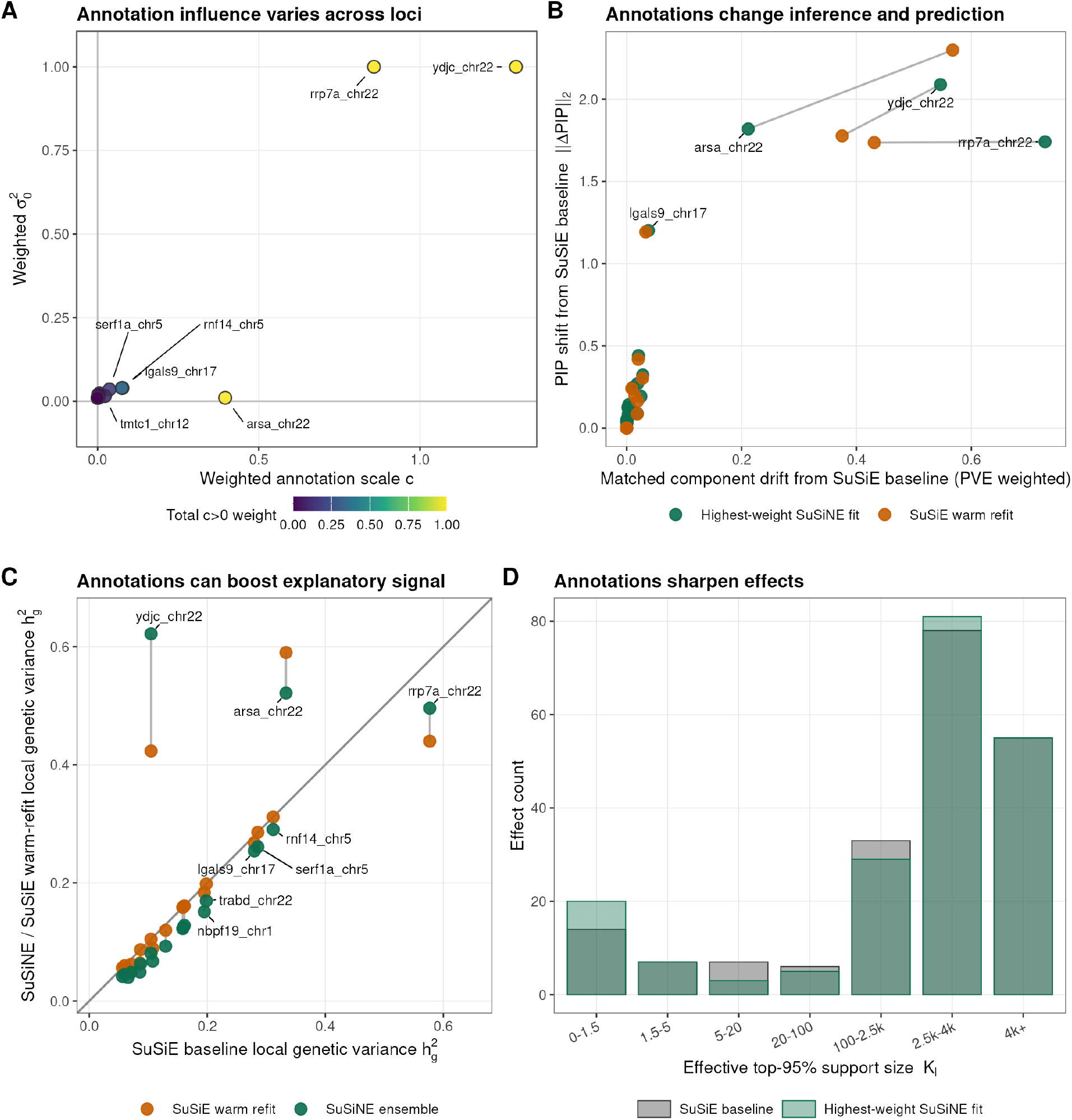
Annotation uptake and fitted-basis movement vary across GTEx Lung loci. **(A)** Annotation uptake: ELBO-weighted prior-mean scale versus weighted prior variance; color shows the aggregate off-zero weight 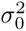 and point size shows the largest single-run weight. **(B)** Drift from the SuSiE anchor for the highest-weight source run and its warm-vanilla refit. The vertical axis is PIP L2 distance; the horizontal axis is PVE-weighted matched-component drift *D*_MC_, as described in Methods. **(C)** Posterior expected local genetic variance for the SuSiE anchor, SuSiNE ensemble, and warm refit. **(D)** Per-effect diffuseness *K_ℓ_* for the anchor and highest-weight source; lower bins indicate sharper effect posteriors. Negligible-mass effects are excluded.

**Table 4.**
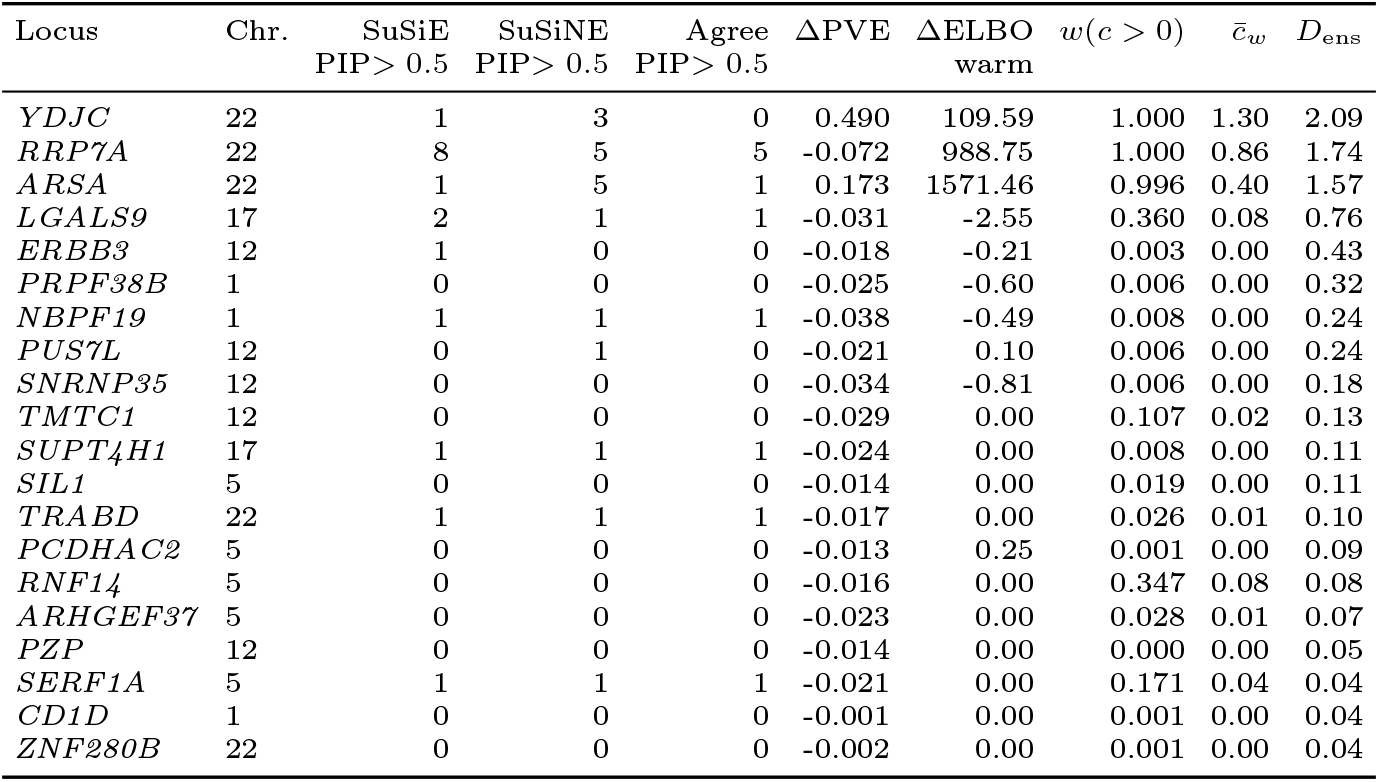
Per-locus summary of the GTEx Lung real-data study, ordered by ensemble-to-anchor PIP distance *D*_ens_. PIP columns count variants with PIP *>* 0.5 under the anchor, the ensemble, and both fits. ΔPVE is the ensemble minus anchor local-genetic-variance diagnostic used in Fig 8c. ΔELBO warm is the annotation-free warm-refit ELBO minus the cold SuSiE anchor ELBO. *w*(*c >* 0) is the total aggregate weight assigned to annotation-informed ensemble members, and *c̄_w_* is the ELBO-weighted prior-mean scale.

#### Annotation influence across the panel

Annotation influence was strongly locus-specific (Fig 8a). Three loci placed near-complete aggregate weight on annotation-informed fits, with total aggregation weights across all off-zero (*c >* 0) models greater than 0.99: *YDJC*, *ARSA*, and *RRP7A*. Four additional loci formed an intermediate-response group, *LGALS9*, *RNF14*, *SERF1A*, and *TMTC1*, with total off-zero weight between 0.10 and 0.36. From the unrounded weights, thirteen of the twenty loci placed less than 0.1 weight on annotated fits and ten placed less than 0.01, consistent with near-zero annotation uptake when the signed channel was not supported by the locus-level association evidence. The strongest responders did not all use the same part of the grid: *YDJC* and *RRP7A* selected the largest prior variance 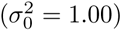when annotations were influential, whereas *ARSA* showed near-complete annotation response in the tight-prior regime 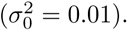

#### Reference-LD consistency of the responder movements

Because the real-data fits use a 1000 Genomes reference panel rather than in-sample GTEx LD, we use this case study to interpret model behavior rather than to make definitive biological calls about individual variants. We screened every responder locus for *z*–LD inconsistency before reading any individual variant as signal. We computed *estimate_s_rss* per locus, ran *kriging_rss* to compare each variant’s observed *z* against the value its LD neighbors predict, and audited the genetic variants with high PIP movement for strand ambiguity (Table B in S1 Appendix). We first observe that the responder loci carry more *z*–LD inconsistency than the non-responders (median subset *s* 0.14 versus 0.07; Table C in S1 Appendix), which is expected: a directional prior moves the posterior where the likelihood is flat, and reference-panel mismatch is one of the data patterns that can flatten it. Second, within each high-responding locus, the genetic variants with high PIP movement span a spectrum from LD-consistent to LD-inconsistent. We therefore read the panel as evidence about the overall mechanism instead of as an inventory of biological calls on individual variants.

#### PIP drift, matched-component drift, and durability

Fig 8b reports source-run-to-anchor distance *D*_src_, warm-refit-to-anchor distance *D*_warm_, and the corresponding matched-component drift. *ARSA* is the cleanest durable basin-discovery case: *D*_src_ = 1.82 and *D*_warm_ = 2.30, matched-component drift increases from 0.212 to 0.567, ensemble ΔPVE is 0.173, and the annotation-free warm refit improves ELBO by 1571.46 over the cold anchor. *YDJC* also shifts durably (*D*_src_ = 2.09, *D*_warm_ = 1.78; matched-component drift 0.546 and 0.375; ΔPVE 0.490; warm-refit ΔELBO 109.59), but its key variant has a 7.9-SD kriging discrepancy within a near-perfectly correlated pair. We therefore treat *YDJC* as a reference-LD inconsistency illustrating mechanism, not as a clean annotation-driven result. Warm-refit durability shows that annotation-free IBSS can remain in an alternative fitted basin once initialized there; it does not establish robustness to LD misspecification because all fits use the same 1000 Genomes matrix.

*RRP7A* shows why the matched-component diagnostic is needed. The annotated source and warm refit remain separated from the anchor (source PIP L2 = 1.74; warm-refit PIP L2 = 1.74) and have the largest component-level rearrangement in the panel (*D*_MC_ = 0.728 for the source and 0.431 after warm refit). This is major effect reallocation, not just PIP reshuffling or discovery of additional fitted signal: the ensemble does not increase fitted signal (ΔPVE = −0.072), even though its annotation-free warm-refit ELBO is +988.75 above its cold anchor. By contrast, if components are summed before comparison, the corresponding total-predictor reference-LD drift is only 0.134 for the source and 0.130 after warm refit. *RRP7A* is therefore a direct example of component-level basis movement that would be understated by a total fitted-value diagnostic alone. The intermediate-response loci show less durable movement after prior removal. *RNF14* and *SERF1A* have nonzero annotation uptake (*w*(*c >* 0) = 0.35 and 0.17), but their displayed PVE changes are small and negative. *LGALS9* remains ambiguous, with intermediate off-zero support (0.36), persistent warm-refit PIP drift, modest matched-component drift (0.038 at the source and 0.033 after warm refit), a modest negative ΔPVE, and a warm-refit ELBO below the cold anchor. Most other loci show little fitted-basis movement, consistent with within-basin reweighting or nonresponse rather than durable basin discovery.

#### Aggregation and model-specification uncertainty

The intermediate-response loci also show why exploration alone is insufficient. *LGALS9* is the clearest example: three PIP clusters carry essentially all of the cluster-weight aggregate mass, with the largest cluster receiving less than half of the weight. These clusters include both annotation-free and annotation-informed fits (aggregate weights 0.64 at *c* = 0 and 0.36 at *c >* 0). The substantial weight on *c* = 0 clusters shows that this multimodality is not only a consequence of nonzero prior means: even annotation-free perturbations can reveal alternative summaries that the cold anchor did not report. Nevertheless, their PIP summaries are not interchangeable: the highest-weight source has PIP L2 = 1.20 from the anchor and the warm refit remains similarly shifted (PIP L2 = 1.19), while matched-component drift remains modest (*D*_MC_ = 0.038 at the source and 0.033 after warm refit). This is the real-data analogue of the model-specification barrier in Scenario 3 of Fig 1: several comparably supported configurations explain nearly the same fitted signal, but a single SuSiE-family fit can report only one of them. Cluster-weight aggregation preserves this OR-of-ANDs-like uncertainty instead of forcing the analysis to choose the top source run.

The real-data multimodality benchmark in Fig E in S1 Appendix supports the same conclusion at the panel level. When the 20 GTEx loci are overlaid on the C-CS simulation background using the same credible-shift cluster count and maximum credible-shift distance diagnostics as the ensemble simulations, the real loci span a broad range of multimodality: some, such as *ZNF280B*, sit near the low-cluster, low-distance corner, whereas *RRP7A*, *NBPF19*, *TRABD*, and *TMTC1* occupy the high-cluster or high-distance tail. Thus the multimodality that motivates cluster-aware aggregation is not an artifact of the simulated benchmark alone.

#### Effect diffuseness

At the study level, the highest-weight source fits sharpen some effects but do not uniformly reduce diffuse uncertainty (Fig 8d). Here *K_ℓ_* is the effective top-95% support size defined above. The sharpest support bin, *K_ℓ_ <* 1.5, increases from 14 to 20 effects, while the 1.5–5 bin remains at 7 effects. Intermediate bins mostly decrease: 5–20 drops from 7 to 3, 20–100 from 6 to 5, and 100–2500 from 33 to 29. The most diffuse tail is mixed rather than uniformly reduced, with 2500–4000 increasing from 78 to 81 and 4000+ unchanged at 55. Thus, the real-data behavior mirrors the simulation mechanism qualitatively: annotations can sharpen specific effects and expose alternative basins, but the locus panel remains heterogeneous rather than a uniform shift toward low diffuseness.

#### Selected-locus zoom

A six-locus, variant-level zoom across the full annotation grid is provided in S1 Appendix (Fig G). At the responder loci *YDJC*, *ARSA*, *RRP7A*, and *LGALS9*, PIP drift is banded rather than gradual. Large slices of annotation settings snap IBSS into a small number of tight, deep alternative-basin optima, with maximum adjacent-grid jumps in ∥***p***_run_ − ***p***_anchor_∥_2_ exceeding 1.4, rather than shifting the optimum slightly within one basin. Read against the LD-consistency screen (Table B in S1 Appendix), the cleanest variant-level evidence is at *ARSA*. There, *chr22_50628562_C_T_b38* rises from anchor PIP 0.020 to 0.857 in the ensemble and 0.997 in the warm refit, while sitting within its LD context (kriging deviation 1.7 SD, unflagged). *YDJC* is the cautionary case. Its two near-perfectly correlated variants (*r* = 0.997) take large opposite-sign conditional effects that match opposite-sign annotations, but one has an observed *z* = 0.11 against a kriging-predicted −3.31 (a 7.9-SD discrepancy). The prior is therefore resolving a reference-LD inconsistency rather than a clean biological call. *RRP7A* shows the same discontinuous grid behavior, but with effect reallocation rather than fitted-signal gain. *LGALS9* remains less decisive under warm refit, and its separated L2 bands match the multimodality result above.

## 3 Discussion

SuSiNE addresses three distinct problems that can look alike in a single SuSiE fit, but require different responses. LD ambiguity is a limitation of the genotype data. Selecting causal variant configurations between correlated candidates is non-convex optimization over a combinatorially large search space; and uncertainty between multiple posterior basins is unrepresentable in a single factorized variational summary. The signed prior-mean channel supplies directional external information for the first problem; the ensemble explores alternative fits for the second; cluster-aware weighting summarizes the alternatives for the third. The simulation results support the practical value of this combination for prioritizing the eight largest-effect causal variants in PIP mass. The results also showcase the utility of well-designed diagnostics: purity remains a useful description of LD, but automatic purity filtering discards informative signal in both the toy example and the oligogenic benchmark and should not gate reported PIPs by default.

While the evidence provided for how annotations *may* improve fine-mapping is clear, the ground truth on genetic fine-mapping causality remains largely unmeasured directly, which means the ground-truth on S2F model annotation behavior is fuzzy. For our simulations, the AlphaGenome calibration bridges published GTEx eQTL benchmarks to a simple signed-Gaussian generator. It does not reproduce the structured errors of AlphaGenome, Enformer, Borzoi, or Decima [22–25]. Performance can vary with the training data, assay, tissue, distance to the transcription start site, and the distinction between statistical fine-mapping labels and biological ground truth. The simulator likely overstates unsigned rank information, which may favor functional-*π*. At the same time, its signed-Gaussian form may favor the prior-mean channel. The observed advantages of the prior-mean channel, and its simulation performance in general, are therefore informative, but it is not a fully generator-independent channel comparison. Held-out tissues, sign errors, magnitude miscalibration, TSS-distance structure, and cross-variant error dependence may be important next simulation targets.

The signed channel self-gates when annotation error is independent of association noise. Alignment between an incorrect annotation and the observed association then tends to average away, and the regularization term prevents unbounded inflation. That protection does not apply when the two error sources align. In the null-contamination sensitivity analysis, deliberately aligning annotations with marginal association noise among null variants erased the gain by *λ* = 0.3. The GTEx low-PIP analysis is best described as an internal null-alignment diagnostic, not a ground-truth negative control. Its near-zero correlations argue against strong genome-local null alignment in this panel, but they do not firmly establish independence.

This sensitivity result also identifies a concrete future circularity risk. Sequence-to-function models are often trained or evaluated using fine-mapping-derived labels. If a model is trained on SuSiE-derived labels and its predictions are then supplied to SuSiNE as annotations, later labels can partly inherit the same sequence-model signal that is being evaluated. The resulting dependence can align annotation error with association-derived evidence. This is precisely the condition under which the aligned-null sensitivity analysis shows that self-gating fails. This is not evidence that the present AlphaGenome annotations are contaminated in this way, but it is direct evidence that the proposed train-label-use cycle could create a failure mode once annotation-informed fine-mapping becomes part of the S2F training labels. The risk is likely smaller for high-PIP causality labels that are strongly data-dominated and larger for more ambiguous inferences. Future S2F evaluations should preserve annotation-free fine-mapping labels, maintain strict lineage separation between training annotations and evaluation labels, or explicitly model the induced dependence.

Several mechanisms could create correlated error even without this S2F-to-fine-mapping circularity. Duplicated regulatory architectures under epistatic selection in paralogous genome regions can make a sequence model and an eQTL analysis favor the same wrong variants. Distance to the TSS is another measurable channel. Annotation accuracy and cis-eQTL discoverability can both vary with distance, so their errors need not be independent where those trends align. Paralog-aware diagnostics, distance-stratified audits, tissue-specific annotation-error models, and explicit down-weighting of the prior-mean channel in high-risk regions are natural safeguards to explore in future work.

The proposed ensemble does not fully solve exploration or aggregation in simulations. Most of the C-CS gain in AUPRC disappeared after the functional prior was removed. C-CSR, which corresponds to the simple SuSiE warm-start on the SuSiNE C-CS fit, retained only 0.0094 of the 0.0656 absolute AUPRC gain, compared to a 0.038 absolute gain for the warm-starting SuSiE on the true configuration. This indicates that the gain from baseline SuSiE to SuSiNE C-CS is mostly due to the reduced LD-ambiguity, rather than fully saturating the exploration of alternative basins. In addition, the truth-aware best-fit aggregation method on C-CS scored 0.089 gain over baseline, which is substantially higher than the credible-weighted aggregation (0.0656), indicating that the credible-weighted aggregation does not fully solve the aggregation problem. While cluster-aware ELBO weighting performed best among the summary-statistic aggregation rules tested here, stacking on held-out predictive performance would be preferable when individual-level data are available [29, 30].

There is also no universally correct locus-level value of *c*. The general target, *c^★^* = Cov(*β, a* | C)*/* Var(*a* | C), conditions on the unobserved causal set and depends on annotation units. The special equality *c^★^* = *ϕ_a_* holds only for the normalized generator used in S1 Appendix. A single locus therefore cannot generally identify the desired annotation-quality scale without extra assumptions. This motivates the fixed grid and aggregation. SuSiNE-EB remains useful as a fast diagnostic, but not as a replacement for the recommended grid-plus-aggregate workflow. Direct scale estimation should pool information across many loci.

The GTEx Lung analysis demonstrates method behavior, not definitive variant discovery. It combines GTEx summary statistics with 1000 Genomes EUR reference LD. LD–*z* inconsistency can therefore create confident conditional effects even after allele matching and residual-variance screening [3, 16, 31]. Heterogeneous effective sample sizes remain a further limitation [32]. The warm refit uses the same reference LD and cannot resolve shared LD misspecification. Of the durable responders, *ARSA* is the cleanest LD-consistent annotation-driven shift. *YDJC* coincides with a 7.9-SD kriging discrepancy in a near-perfectly correlated pair, and *RRP7A* reallocates components without increasing fitted signal. The biological impact is therefore limited to a single clean locus, while mechanistic impact is observed across multiple loci. In-cohort LD is the appropriate standard for actionable variant interpretation. Tissue and ancestry generalization remain untested.

The central benchmark uses one oligogenic architecture with *h*^2^ = 0.25, *n* = 600, *p* ≈ 1000, and European samples, while varying annotation quality. The architecture checks support the consistent sign of improvement under the tested sparse and diffuse settings, but the magnitude varies: gains to recall at high precision can remain small in weak or diffuse architectures. The full C-CS workflow costs 64 mutually independent fits per locus. A 4 × 4 subgrid recovered similar AUPRC at roughly one quarter of the serial wall time in this benchmark, and the grid is embarrassingly parallel. Because most GTEx loci placed little aggregate weight on annotation-informed fits, a two-stage protocol may be useful in future work. One could run a single-fit anchor broadly and reserve the ensemble for loci with evidence of annotation uptake or credible-shift multimodality. The performance of such a screening rule would need to be established before deployment.

The pathology results clarify why new diagnostics matter. A converged, highly confident SuSiE fit is not necessarily a faithful readout. Scenario 3 shows that the default SuSiE method can still fully commit to just one of several nearly equal configurations. Scenario 4 shows that a clearly better basin can still be missed, due to default initialization and greedy optimization. Level of PVE, ELBO, PIP, and fitted-basis agreement across different starts are therefore useful diagnostics. The contrasts among default SuSiE, truth-warm SuSiE, C-CS max-ELBO, C-CS aggregation, C-CSR, and the truth-aware single-fit oracle form a diagnostic ladder. They are not an additive causal attribution because the interventions are not nested counterfactuals. The same restraint applies to purity. It is an LD summary, not a default quality gate. We retain a purity threshold only in the refinement branch generator to mimic the established SuSiE refinement strategy, though effect diffuseness or basin movement may ultimately be better branch triggers.

The diagnostics also have different domains of use. Effect diffuseness *K_ℓ_* requires only fitted posterior components. The accuracy ratio and truth-basis divergence require the true causal set, so they are simulation-evaluation measures rather than real-locus outputs. Matched-component drift is the corresponding truth-free real-data diagnostic. It can reveal component-level movement that is hidden when fitted effects are added together. This distinction complements the paper’s representational argument, that SuSiE’s AND-of-ORs factorization is efficient, but it cannot itself hold mutually exclusive OR-of-ANDs configurations. Exploration and summarization are therefore separate tasks. Similar ideas may be useful for other mean-field variational approximations to multimodal targets [33, 34]. They remain distinct from multi-trait extensions such as mvSuSiE, which share information across correlated traits [35].

We do not require a direct PolyFun benchmark to establish this mechanism. PolyFun supplies unsigned per-variant heritability estimates through the prior-inclusion channel ***π*** [17]. SuSiNE supplies signed annotations through the prior-mean channel ***µ***_0_. We show in simulations that coercing the signed source into an unsigned score discards valuable information for model inference. The opposite comparison, passing unsigned heritability estimates into a signed prior mean, is not meaningful because the sign is undefined. PolyFun is therefore complementary rather than redundant, and combining heritability-based ***π*** with signed ***µ***_0_ is a natural extension, but would not form the basis of a direct benchmark.

The next steps follow directly from these limitations. They include empirically characterizing correlated annotation and association errors, building a pooled cross-locus model for annotation quality, combining the ***π*** and ***µ***_0_ channels, using stacking when predictive densities are available, propagating calibrated sequence-to-function uncertainty into the prior mean, and developing search procedures that are less dependent on greedy IBSS initialization. These directions are particularly relevant for cTWAS-style gene-level analyses and regulatory-network priors, where LD-driven confounding makes the separation of evidence sources essential [36, 37].

In summary, signed annotations and multi-fit summaries improved high-effect causal-variant prioritization in the evaluated regime. The sensitivity checks identify clear boundaries on that gain. The signed prior-mean channel preserves SuSiE’s tractable single-effect and summary-statistic structure. The broader contribution is also diagnostic and expository, as LD ambiguity, optimizer failure, and cross-configuration uncertainty leave different signatures and call for different responses. A single variational fit can be useful, but it does not by itself establish a faithful causal readout.

## 4 Materials and methods

### 4.1 SuSiE pathology

The four engineered examples obey SuSiE’s sparse linear Gaussian assumptions but are not intended to resemble typical eQTL loci. We fit each dataset with *L* = 5 from the package default, a truth-warm start, and a decoy-warm start. Residual and prior variances used SuSiE’s usual empirical-Bayes updates. Complete constructions, parameters, and numerical results are in S1 Appendix Section 1 and Table A.

### 4.2 SuSiNE model specification

The SuSiNE model extends SuSiE by allowing non-zero prior means on variant effects. For *n* individuals and *p* variants with genotype matrix **X** ∈ ℝ*^n×p^* and phenotype **y** ∈ ℝ*^n^*:

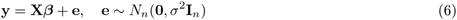

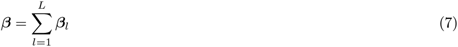

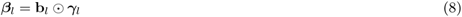

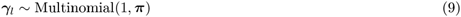

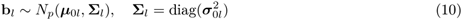

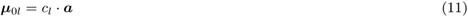

Here ⊙ denotes elementwise multiplication, ***γ****_l_* ∈ {0, 1}*^p^* is a one-hot vector indicating which variant carries the *l*-th effect, and **b***_l_* ∈ R*^p^* contains effect sizes drawn from a multivariate normal with mean ***µ***_0_*_l_* and diagonal covariance **Σ***_l_*. The prior mean ***µ***_0_*_l_* is the product of a scalar *c_l_* and a vector derived from functional annotations, ***a***. The prior inclusion probabilities ***π*** are shared across effects and can optionally incorporate functional information as in PolyFun [17], but in this work we focus on uniform ***π*** to isolate the contribution of the new ***µ***_0_ channel, unless otherwise specified.

Empirical Bayes updating of *c_l_* and 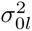 is supported because these parameters do not vary by variant, preserving coordinate-ascent tractability (S1 Appendix Sections 5 and 11). This is the same high-level role empirical Bayes often plays in high-dimensional prediction, learning prior and tuning parameters from many related variables, especially when external co-data are available [38].

### 4.3 Ensembling and aggregation

For a single dataset, an *ensemble* contains fits obtained by changing the initialization, prior scale, prior variance, or refinement path. Each fit *m* returns a PIP vector ***p̂****_m_* and final ELBO L*_m_*. The aggregation step combines these fits into a per-variant summary.

We evaluate several aggregation rules, using the same ensemble members as input. The *max-ELBO* rule selects the PIP vector from the fit with largest L*_m_*. The *uniform* rule averages with equal weighting PIPs across all ensemble members. The *ELBO-softmax* rule weights each fit in proportion to exp(L*_m_* − max*_m_*′ L*_m_′*). Our primary rule, *cluster-weight* aggregation, first clusters fitted PIP vectors with complete-linkage hierarchical clustering under a credible-set-emphasizing PIP distance *d*(***p***, ***q***) = max*_j_* max(*p_j_, q_j_*) |*p_j_* − *q_j_*|, cutting the tree at *d* = 0.05; this *credible-shift* metric weights each per-variant disagreement by the larger of the two PIPs, so clusters split only when fits disagree about high-confidence variants rather than about negligible tail mass. Clusters are then aggregated, such that each cluster receives a weight proportional to 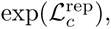, where 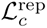 is the largest ELBO in the cluster, and that weight is distributed within the cluster across its member fits by ELBO-softmax (at unit temperature). The cut *d* = 0.05 and the softmax temperature were chosen from ablations in which aggregation performance was locally flat in both; results are insensitive to either within that region, so we fix these values rather than report the sweeps. This is what makes cluster-weight frequency-agnostic, and it is the key difference from plain ELBO-softmax. Under ELBO-softmax, a basin that many ensemble members reach as near-duplicate fits gets counted once per fit, so its weight inflates with the number of runs that happen to land there. Cluster-weight merges those near-duplicates first and scores each basin by the evidence of its best member, so a higher-ELBO posterior optimum found by only one or two fits is not drowned out by a crowd of near-identical fits at a lower-ELBO optimum. Note that this clustering operates on PIP vectors rather than on the explanatory basis, so its clusters often approximate basins, but can also subdivide one: a directional prior that reweights within-group PIP mass without moving the fitted **ŷ** stays in the same basin, yet may form a distinct PIP cluster. In simulation only, we also report a truth-aware best-member benchmark, labeled *oracle* in legacy figure and output labels, that selects the single ensemble member with the best AUPRC. This benchmark requires ground truth and is not a deployable aggregator or an optimized mixture. The three ELBO-based deployable rules, max-ELBO, ELBO-softmax, and cluster-weight, use the per-fit ELBO L*_m_* as a tractable stand-in for the log marginal likelihood of fit *m*. Uniform averaging does not use ELBO. We acknowledge that this is the maximized variational bound, not the evidence itself. Using the bound this way is standard for variational model comparison [39, 40], but it is an approximation. Its limitations and the stacking-based alternative available with individual-level data are discussed above.

### 4.4 Simulation studies

#### Overview

The baseline screen and ensemble study use the same genotype matrices, phenotype seeds, and causal architectures. This paired design isolates differences due to the fitted models and workflows. The annotation-quality settings are given separately for each experiment.

#### Genotype data

Genotype matrices are selected from a simulated EUR genotype dataset of 120,000 individuals [41]: 150 loci of *n* = 600 individuals and *p* ≈ 1000 SNPs each. The selected matrices are stratified across the available range of a locus-level LD-summary statistic before keeping 150 matrices for the study. For each locus, SNPs are filtered to MAF ≥ 0.01 within the 1000 variants closest to the gene TSS. The same 150 genotype matrices are reused across the simulation experiments, so baseline and ensemble comparisons are paired on locus-level LD structure.

#### Phenotype simulation

We adopted the oligogenic architecture from SuSiE 2.0 [28] (their Supplementary Notes S4), designed to mirror eQTL effect structure while constraining total compute to a single realistic architecture. For each locus, we sampled 23 causal variants uniformly without replacement and partitioned them into three fixed tiers: 3 sparse/strong effects, 5 oligogenic/moderate effects, and 15 polygenic-background effects. Raw effects were drawn independently by tier: sparse effects from *N* (0, 1); oligogenic effects from a two-component Gaussian mixture with probability 0.35 on *N* (0, 0.8^2^) and probability 0.65 on *N* (0, 0.25^2^); and polygenic-background effects from *N* (0, 0.08^2^). The three tiers were then rescaled so that they contributed 50%, 35%, and 15%, respectively, of the total squared-effect energy 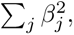 preserving the intended sparse-to-polygenic hierarchy before the phenotype-level heritability calibration. Phenotypes were generated from the linear predictor *η* = **X*β*** with target total heritability *h*^2^ = 0.25. Specifically, the residual variance was set to

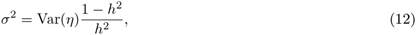

and responses were drawn as **y** = *η* + ***ε*** with *ε_i_* ∼ *N* (0*, σ*^2^). Four independent phenotype seeds per genotype matrix yield 150 × 4 = 600 simulated datasets. Seed management was deterministic and indexed by both the genotype matrix and phenotype replicate, making effect and noise draws reproducible while preventing accidental reuse of identical causal-effect draws across loci.

#### Annotation simulation

To assess SuSiNE’s sensitivity to annotations of varying quality, we generate simulated annotations parameterized by two interpretable knobs: *ϕ_a_*, the squared correlation between the annotation vector and the true causal effects restricted to causal positions, and *ν_a_*, the ratio of non-causal to causal annotation variance. Denoting the causal support by *T* ⊆ {1*, …, p*} and its complement by *T ^c^*, draws are constructed such that

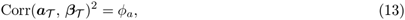

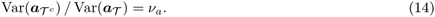

At causal positions, annotations are generated as a Gaussian signal-noise mixture around the true effects. The centered RMS of the causal annotation block is fixed relative to the causal-effect RMS, while *ϕ_a_* changes the signal-to-noise mixture within that block. At non-causal positions, annotations are centered Gaussian noise whose second moment is set relative to the causal block by *ν_a_*. The total RMS across causal and null variants is not held fixed when *ν_a_* changes. Thus *ϕ_a_* controls directional accuracy on causal variants, while *ν_a_* controls how much null annotation magnitude can compete with causal annotation magnitude. The regime ranges from fully informative at (*ϕ_a_, ν_a_*) = (1, 0) to fully uninformative at (0, 1). We restrict *ν_a_* ≤ 1 because larger values would create an anti-informative regime in which null annotations are systematically larger than causal annotations.

#### Annotation-quality calibration against AlphaGenome

Informative priors can sharpen inference when the annotations are accurate and mislead it when they are not. We therefore used AlphaGenome’s published GTEx eQTL benchmarks to place the simulator in a realistic quality range. AlphaGenome reports a zero-shot auROC of 0.71 for distinguishing high-confidence fine-mapped variants from distance-matched controls. It also reports signed correlations near 0.50 between its scores and SuSiE posterior effect sizes, with a tissue-weighted mean of 0.49. The corresponding unsigned correlation is only 0.10.

The simulator parameters do not map directly to any one of these quantities. We constructed matched metric analogues over an 11 × 11 grid using all 600 phenotype datasets. The causality analogue scores the three largest true causal effects against true null variants. The coefficient analogue ranks all 23 causal effects.

Because the causality AUPRC is measured at very low prevalence, we convert it to a balanced-auROC equivalent under an equal-variance Gaussian score model. Matching the causality and signed-coefficient anchors gives *ν_a_* ≈ 0.95 and *ϕ_a_* ≈ 0.3. The baseline screen uses the nearest coarse-grid point (0.3, 0.9), while the ensemble study fixes *ν_a_* = 0.95.

This calibration is a bridge between evaluation settings rather than a claim that the generator reproduces AlphaGenome’s full error distribution. Matching the signed benchmark produces more unsigned rank information than AlphaGenome reports, which may favor the functional-*π* comparator. The generator’s signed-Gaussian form may also favor the prior-mean channel. Full definitions, the AUPRC-to-balanced-auROC conversion, and the calibration figure are in S1 Appendix Section 2 and Fig A.

The empirical best single-fit baseline-grid value *c* = 0.642857 *…* is not an estimate of the dimensionless *ϕ_a_* = 0.3. The coefficient *c* depends on the units of the annotation. The simulator fixes those units relative to causal-effect RMS, while the real-data workflow uses a probit transform, within-locus RMS normalization, and a locus-specific effect-scale anchor. Rescaling is therefore the primary reason the empirical coefficient need not equal the unit-loading special case *c^★^* = *ϕ_a_*. PIP coupling, LD competition, AUPRC optimization, and interaction with 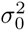 provide additional differences between the theorem and the empirical grid. S1 Appendix Section 2 gives the full interpretation.

#### 4.4.1 Baseline single-fit screen

The screen compared eight specifications (Table 2). Annotation-consuming specifications were run across the full 3 × 3 annotation-quality grid. Annotation-agnostic specifications were fit once per dataset. SuSiE specifications used *susieR* [15], while SuSiNE specifications used the *susine* R package developed for this work.

The SuSiNE-vanilla specification serves as a replication check, fitting SuSiNE at *c* = 0 and confirming agreement with *susieR* across all datasets. SuSiE-functional-*π* supplies annotation-informed priors through the inclusion-probability channel via a temperature softmax on the absolute annotation values,

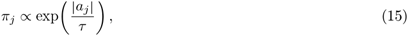

with *τ* as the temperature hyperparameter. We sweep 10 log-spaced *τ* values from 0.01 to 10. Small *τ* concentrates prior mass on high-|*a*| variants, while large *τ* approaches a uniform prior. Because this formulation uses |*a_j_*|, the *π* channel discards the sign of the annotation by construction. This is the key distinction from the ***µ***_0_ channel in SuSiNE. The SuSiNE-functional-*µ* grid sweeps eight equally spaced values of *c* and the five values of 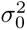 reported above, for 40 fits per dataset.

##### Inference settings

All fits use *L* = 10 maximum effects and a cap of 100 IBSS iterations. Credible sets are constructed at cumulative PIP threshold *ρ* = 0.95. Primary PIP scoring and manuscript-facing summaries use the unfiltered fitted *α* matrices, while purity, credible-set size, and effect diffuseness are retained as diagnostics; this departs from common purity-filtered reporting in light of the metric-brittleness analysis in Section 2.1.

##### Effect-level diagnostics

To diagnose *why* functionally informed priors do or do not improve fine-mapping, and to study the behavior of functional-*π* and functional-*µ*_0_ channels, we introduce two effect-level metrics that decompose effect quality into orthogonal diffuseness and accuracy axes. For each fitted single-effect component *ℓ* with PIP vector ***α****_ℓ_*, let ***α***^(95)^ denote the restriction of ***α****_ℓ_* to the smallest set of variants whose cumulative mass reaches 0.95, renormalized to sum to 1. We then define the *effective top-95% support size K_ℓ_* and the *accuracy ratio A_ℓ_*:

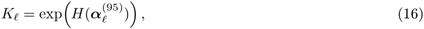

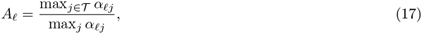

where *H*(·) denotes Shannon entropy and T is the set of true causal variant indices. *K_ℓ_* is the number of variants that would reproduce the observed top-95% entropy under a uniform distribution, quantifying how *diffuse* the top of the effect posterior is; small values indicate a peaked effect and large values indicate mass spread across many variants. *A_ℓ_* is the ratio of the largest PIP among true-positive variants to the largest PIP overall, quantifying how *accurate* the effect is in pointing toward a true causal variant: *A_ℓ_* = 1 means the top-ranked variant has a PIP which is tied with the causal variant, and *A_ℓ_ <* 1 means some non-causal variant outranks every causal variant.

For the purity-filter diagnostic in the baseline SuSiE analysis, we used the annotation-agnostic SuSiE-vanilla fit with *c* = 0 on the same 600 oligogenic phenotype datasets. The analysis unit was the fitted single-effect component. After excluding inactive components with empty credible sets, this gave 6,000 active effects across the 600 datasets. We recomputed combined per-variant PIPs after dropping effects under four rules: purity ≥ 0.50, purity ≥ 0.90, purity ≥ 0.95, and *K_ℓ_* ≤ 100. The unfiltered fit was kept as the reference. For each rule we pooled precision-recall bins across datasets and scored recovery of the top eight causal variants per dataset, matching the baseline-screen AUPRC convention.

We also used this minibatch to compare classical credible-set summaries with the effect-level diagnostics. The truth-agnostic summaries were purity, credible-set size, and *K_ℓ_*. The truth-aware summaries were causal coverage, *A_ℓ_*, and causal PIP mass, defined as the total posterior mass that an effect assigns to the true causal variants. Causal PIP mass was used only as a simulation diagnostic of effect quality. To quantify how well each metric described this response, we fit quasibinomial-logit generalized additive models with REML using *mgcv*. Heavy-tailed size and *K_ℓ_* predictors entered on the log_10_ scale. One-dimensional models used s(x, k = 5), and two-dimensional models used te(x, y, k = 5). All models used the same analysis frame. Five-fold cross-validation was assigned at the genotype-matrix level: all four phenotype datasets generated from the same matrix remained in the same fold, so the folds were genotype-disjoint. We report out-of-sample *R*^2^, held-out Spearman correlation, Spearman correlations among metrics, and *mgcv* concurvity for the two paired planes.

Because the *L* component indices are arbitrary and effects migrate across fits, we compare effect quality across specifications at the level of the *distribution* of (*K_ℓ_, A_ℓ_*) rather than by tracking individually paired components. We report variance-weighted two-dimensional density differences (destination minus source) on the (*K_ℓ_, A_ℓ_*) plane at the high-annotation-quality condition (*ϕ_a_, ν_a_*) = (0.5, 0.9). To separate optimizer-barrier from model-specification-barrier contributions, we run a 5-arm refit chain. In this chain, each cold annotated fit (***µ***_0_ or ***π***) is followed by an annotation-free vanilla warm refit. The warm refit asks whether the basin discovered under the prior is durable under the annotation-free objective. The chain specification and the variance-weighted density comparison are detailed in S1 Appendix (Section 13.1).

##### Truth-basis divergence

Whereas *K_ℓ_* and *A_ℓ_* grade effect quality within a single fit, we also track whether a treatment moves the fitted explanatory basis toward the known data-generating basis. The *truth-basis divergence D*_truth_(*F*) is a variance-weighted 1 − *r*^2^ distance between a fit’s single-effect fitted directions 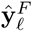 and a synthetic truth basis built from the top-23 causal effects, with components matched one-to-one by the Hungarian algorithm on fitted-**ŷ** correlations. Its per-dataset *reduction* under a treatment is the mechanism-level analogue of per-dataset ΔAUPRC. It measures whether the treatment closed the gap to truth in basis space, independent of any per-variant PIP-ranking effects. The full construction and equation are given in S1 Appendix (Section 13.1).

We use two paper-facing drift summaries because the simulation and real-data analyses have different ground truth. In simulation, truth-basis divergence measures movement toward the known causal basis. In real data the true causal basis is unavailable and the study is run from summary statistics and reference LD, so the paper-facing movement diagnostic is a truth-free matched-component drift, defined below, that aligns fitted components between two models and measures how much their signed component-level fitted effects move under the reference LD matrix while down-weighting loci with little fitted signal.

#### 4.4.2 Ensemble scaling study

The ensemble scaling study uses the same 600 oligogenic phenotype datasets derived from 150 genotype matrices and therefore inherits the same genotype matrices, phenotype seeds, causal architecture, fitted effect count (*L* = 10), and IBSS iteration cap (100 iterations). Its purpose is different from the baseline screen: rather than comparing many single-fit model specifications, it asks which exploration axes and aggregation rules best recover the performance lost to the optimizer and model-specification barriers identified in Section 2.1. Following the baseline results, we carried forward only three model families: an annotation-agnostic zero-prior-mean SuSiE-equivalent fit implemented through the SuSiNE backend, an empirical-Bayes SuSiNE model with signed functional prior means, and a fixed-variance functional-*µ* SuSiNE model with ***µ***_0_ = *c****a***.

##### Annotation settings

Annotation-consuming ensembles were evaluated on a one-dimensional accuracy sweep, with *ϕ_a_* ∈ {0, 0.3, 0.5} and the non-causal annotation scale fixed at *ν_a_* = 0.95. Thus *ϕ_a_* is the target squared correlation between annotations and true effects on causal variants, while the non-causal annotation variance is held near the causal annotation scale. This is a compute-saving specialization of the baseline grid, which swept both *ϕ_a_* and *ν_a_*. The main ensemble figures focus on *ϕ_a_* = 0.3, the AlphaGenome-calibrated accuracy value, with *ϕ_a_* = 0 and *ϕ_a_* = 0.5 included to measure the response to uninformative and stronger annotations.

##### Ensemble construction

We varied four exploration axes. Random-restart ensembles perturb the initial prior inclusion weights ***π*** for each effect by drawing from a symmetric Dirichlet distribution with all concentration parameters set to 0.001, run those perturbed-prior fits, and then refit the converged solutions under uniform prior inclusion weights. The default fit and the annotation-free refits, rather than the perturbed-prior fits themselves, are used as ensemble members.

Refinement ensembles follow the spirit of the *susieR* refinement procedure [15, 16], using credible sets to define local exclusions that can force IBSS to search for alternative basins. Starting from a fit, the procedure identifies effects with a 95% credible set passing the refinement purity threshold (0.95), then creates one branch per eligible effect. In each branch, all variants in that effect’s 95% credible set are temporarily blocked by setting their prior inclusion weights to zero and renormalizing the remaining variants to uniform weights. The model is refit under this perturbed prior, then refit again from that solution with the prior inclusion weights returned to uniform. Branching is repeated recursively until no eligible pure effect remains or the maximum ensemble size is reached. This use of purity is pragmatic and inherited from the SuSiE refinement workflow; it is not an endorsement of purity as a default manuscript-facing filter on reported PIPs.

Prior-mean-scale ensembles vary *c* in ***µ***_0_ = *c****a*** over linear grids from 0 to 1.5. Prior-variance ensembles vary the fixed prior variance 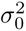 over grids spanning 0.01 to 1.0, always including the baseline value 0.2.

Ensemble labels used in the results concatenate the model family and exploration axes. Prefix A denotes annotation-agnostic zero-prior-mean fits, prefix B denotes empirical-Bayes functional-*µ* SuSiNE fits, and prefix C denotes fixed-variance functional-*µ* SuSiNE fits. The suffixes R, F, C, and S denote random restarts, refinement, prior-mean-scale grids, and prior-variance grids, respectively. For example, C-CS is the fixed-variance functional-*µ* ensemble crossing eight equally spaced *c* values from 0 to 1.5 with 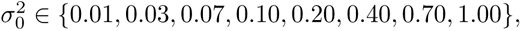 for 64 fits per dataset; this is the primary ensemble highlighted in the results. C-CSR uses the same C-CS search to identify basins, then refits each converged solution under the default zero-prior model 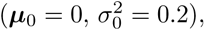 testing whether annotation-discovered basins remain stable after the functional prior is removed.

##### Aggregation and evaluation

Within each dataset and ensemble group, PIPs were aggregated using the max-ELBO, uniform, ELBO-softmax, and cluster-weight rules described above. We also report an “oracle” (truth-aware) best-member aggregator, defined by selecting the individual ensemble member with the highest per-dataset AUPRC before pooling results. This benchmark uses ground truth and is therefore not a deployable method or an aggregation rule; it measures headroom relative to truth-aware selection of one explored fit.

Performance was computed from PIP confusion bins pooled across datasets. For classification-style precision–recall, AUPRC, and PIP-calibration analyses, positives were defined as the top eight true causal variants per dataset ranked by |***β***|; the remaining lower-effect causal variants were omitted rather than counted as negatives. This scoring target matches the sparse and moderate-effect tiers of the oligogenic architecture and avoids treating weak polygenic-background effects as equally recoverable. PIP calibration was assessed by binning the retained variants according to reported PIP and comparing mean PIP with the empirical frequency of membership in this top-eight target. For ensemble-size analyses, we evaluated deterministic sub-ensembles at smaller grid resolutions (for example, 4 × 4 and 8 × 8 C-CS grids) and used observed wall-clock time, averaged per dataset, as the compute-cost measure.

For bootstrap stability checks on the headline ensemble gain, the paper-prep workflow uses a paired bootstrap over the 600 shared phenotype-dataset identifiers. The comparison is paired because the SuSiE-equivalent baseline and each ensemble summary are evaluated on the same datasets. Within each of *B* = 2000 replicates (RNG seed 20260505), phenotype datasets are sampled with replacement, PIP confusion-bin counts are weighted by sampling multiplicity, and pooled AUPRC is recomputed for both methods before taking the absolute difference ΔAUPRC = AUPRC_ensemble_ − AUPRC_baseline_. We report percentile 95% intervals for this absolute gain. The four phenotype datasets that share a genotype matrix are separate bootstrap units.

#### 4.4.3 Sensitivity checks

We ran three additional checks on the recommended cluster-weight C-CS workflow. All sensitivity simulations reuse the same 150 genotype matrices and four phenotype seeds per matrix, giving 600 datasets per grid setting, and use the same *L* = 10 cap, IBSS iteration cap of 100, credible-set level *ρ* = 0.95, and pooled confusion-bin scoring as the ensemble study. To reduce compute, the sensitivity C-CS grid uses a 4 × 4 crossing of *c* ∈ {0, 0.43, 1.07, 1.50} and *σ*^2^ ∈ {0.01, 0.10, 0.40, 1.00}, summarized by the same cluster-weight rule. Each setting is compared with an annotation-free single-fit SuSiE-equivalent baseline run on the same datasets.

The genetic-architecture sensitivity check asks whether the headline gain depends on the oligogenic architecture used above. The sparse arm uses *p^★^* ∈ {1, 3, 5} causal variants and per-causal heritability 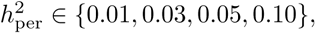 with all causal variants treated as positives. Since sparse effects are constant and positive in this construction, annotations are generated in a sign-prior mode rather than by centering heterogeneous effects. The diffuse arm uses 100 causal variants with total local heritability *h*^2^ ∈ {0.05, 0.15, 0.25}; precision-recall metrics score the top eight causal variants by absolute effect so that the endpoint is not dominated by many very weak effects. Both arms use the AlphaGenome-calibrated annotation setting *ϕ_a_* = 0.3, *ν_a_* = 0.95.

The annotation-contamination sensitivity check probes a failure mode of the self-gating argument. Starting from the same oligogenic simulator and annotation setting as the main ensemble study, we form a contaminated annotation vector by blending the simulated annotation with the marginal *z*-score vector on either the null block or the causal block. For the active block *B*, the marginal *z_B_* values are rescaled to match the RMS of the original annotation block. If 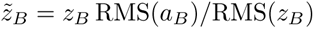 and 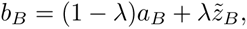 the contaminated block is 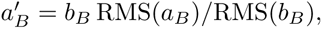 with *λ* ∈ {0, 0.1, 0.2, 0.3, 0.5, 0.7, 1.0}. The inactive block is copied exactly, and diagnostic gates verify that the active-block RMS is preserved and that the inactive block is unchanged. The null arm contaminates only non-causal variants and tests whether spurious alignment between annotation noise and association noise can mislead SuSiNE. The causal arm contaminates only true causal variants and serves as a positive control, since stronger causal annotation–association alignment should help.

Finally, for the GTEx Lung case study we computed a post-hoc internal null-alignment diagnostic that does not refit any model. At each locus, we took the annotation-free SuSiE anchor, selected variants with anchor PIP *<* 0.01, and computed the correlation between the AlphaGenome annotation *a_j_* and the marginal association statistic *z_j_* among those low-PIP variants. This asks whether the real-data annotations are aligned with association noise in the region where the baseline fine-mapper sees little evidence for causality; it is a diagnostic subset, not an external negative control.

### 4.5 Real data study

#### Marginal association statistics

For each gene, we extract per-variant slope *β̂_j_*, standard error se(*β̂_j_*), allele frequency af*_j_*, and minor-allele count ma_count*_j_* from the GTEx v10 Lung *allpairs* parquet file for the gene’s chromosome [3, 42]. Variants are filtered to 0.01 ≤ af*_j_* ≤ 0.99 and effective sample size *n_j_* = ma_count*_j_/*(2 af*_j_*) *>* 50. The marginal z-score is *z_j_* = *β̂_j_/*se(*β̂_j_*), and the standardized marginal effect, used both for the RSS likelihood and for the per-locus annotation scale described below, is

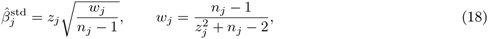

with per-variant standard error 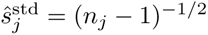 and *w_j_* a bias-correction factor distinct from the annotation *a_j_* defined below.

#### LD reference panel

LD is computed from the 1000 Genomes Project Phase 3 release on GRCh38 [31], restricted to the EUR samples. Genotypes are matched to GTEx variants on chromosome, position, and reference/alternate alleles. Dosages are centered and scaled, with missing calls handled by mean imputation before computing the locus-level Pearson correlation matrix **R**. Because these are reference-panel rather than in-sample LD matrices, we treat LD mismatch as a central caveat rather than as fully solved by preprocessing. As a post-hoc diagnostic, we compute the method-of-moments residual variance estimate *σ̂*^2^ = 1 − *z^⊤^*(**R** + 10*^−^*^4^**I**)*^−^*^1^*z/n* when numerically available and run kriging-style LD–*z* consistency checks for the loci emphasized in the results.

#### Locus selection

Candidate genes are protein-coding genes drawn from the GTEx Lung allpairs files across five chromosomes (chr1, chr5, chr12, chr17, chr22), sampled per chromosome from GENCODE-annotated gene models; for each candidate we retain the gene’s cis variants passing the filters above. Because the study is restricted to a single tissue, many loci are uninformative for method evaluation for opposite reasons: some have little marginal signal to fine-map, while others are so sharply resolved that a signed annotation channel has little room to matter. Selecting loci at random would therefore yield a panel dominated by degenerate cases, so we instead assemble a panel of 20 loci engineered to span the full observed range of fine-mapping difficulty, scored entirely from the annotation-agnostic data.

We summarize each candidate by two coordinates. The first is its *observed* difficulty: the candidate is fit with a vanilla SuSiE RSS baseline screen [15] (*L* = 10, residual variance estimated), and the model-wide PIP vector ***p*** is reduced to an effective support size *k*_eff_ = exp(*H*(***p̄***)), where ***p̄*** = ***p****/ Σ_j_ p_j_* and *H* is Shannon entropy. This is the effective number of variants over which posterior inclusion mass is spread. The second is a *predicted* difficulty, regressed from metrics computable before any fit: three moments of the off-diagonal LD distribution, 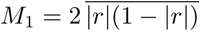 (a “mid-LD energy” maximized at |*r*| = 0.5), 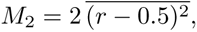 and 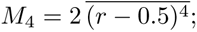 two marginal-signal summaries, the participation ratio 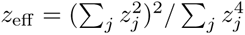 (an effective number of strong marginal signals) and the count of variants with |*z_j_*| *>* 3; and a variant-count funnel (raw, post-MAF, post-sample-size, exported, master, and 1000G-matched/unmatched variant counts). Count-valued features are log(1 + ·)-transformed and all features are robustly standardized by median and IQR. Predicted *k*_eff_ is the held-out prediction of a five-fold cross-validated principal-component regression of log(1 + *k*_eff_) on the leading eight principal components of these features.

We then select the panel by farthest-point (maximin) sampling in a two-dimensional difficulty plane. Its coordinates are log(1 + *k*_eff_) for predicted difficulty and log(1 + *k̂*_eff_) for observed difficulty, with both axes robustly standardized. Starting from a fixed anchor locus for reproducibility, we repeatedly add the candidate whose minimum Euclidean distance to the already-selected set is largest, until 20 loci are chosen, without imposing region quotas. This spreads the panel across the difficulty plane rather than concentrating it in any single regime. The selection is locked prior to AlphaGenome scoring, making it independent of the annotation channel under evaluation. We emphasize that the predicted-difficulty axis serves only as a sampling device: across the 372 screened candidate loci its cross-validated predictions recover only a coarse ordering of observed *k*_eff_ (Spearman *ρ* ≈ 0.64, with large absolute error), so we use it as a screening heuristic for assembling a difficulty-spanning panel and make no claim that fine-mapping difficulty is reliably predictable from dataset-observable metrics.

#### AlphaGenome annotation generation

For each selected locus we score every retained variant with AlphaGenome [25] using the public API under elevated access granted by the AlphaGenome team. Each request submits a 1 Mb sequence interval centered on the gene’s transcription start site when both alleles fit within the canonical TSS-centered window; otherwise the interval is recentered on the midpoint of the gene’s exon span, again subject to both alleles fitting inside a 1 Mb interval. Variants that fit neither window are logged and excluded from the annotation channel. We use the RNA-seq output head together with AlphaGenome’s *GeneMaskLFCScorer*, which returns a per-variant log-fold-change score on the target gene’s expression, and restrict the returned rows to source *gtex* and tissue *Lung* to match the GTEx Lung cis-eQTL context. The pipeline handoff provides a single signed bounded AlphaGenome quantile score *q_j_* ∈ (−1, 1) per variant, which is used as the input to the prior-mean transformation below. This same annotation-quality regime corresponds to the calibration point used in the simulation study (Section 4.4, S1 Appendix Fig A): a balanced-case causality auROC of approximately 0.71 and signed-coefficient Spearman *ρ* ≈ 0.49 on the AlphaGenome-reported GTEx benchmarks.

#### Annotation transformation and prior-mean assembly

The locus-aligned annotation vector ***a*** is constructed by mapping the bounded quantile score *q_j_* to a real-valued effect-size proxy under a probit transform, clipping for numerical stability, and RMS-normalizing within the locus:

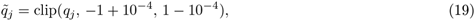

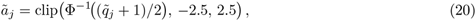

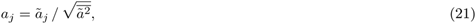

where Φ*^−^*^1^ is the standard-normal quantile function and 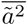 is the locus mean of squared annotations. RMS normalization (rather than *z*-score standardization) is deliberate: *a_j_* = 0 preserves its semantic meaning of *no signed AlphaGenome evidence at variant j*. Variants for which AlphaGenome returned no score (e.g. excluded by the window-eligibility filter) are assigned *a_j_* = 0 and flagged for diagnostics.

To set the per-locus prior-mean scale on the same units as the standardized marginal effects, we anchor a baseline scalar *c*^base^ to two locus-internal references and take the smaller of the two:

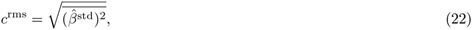

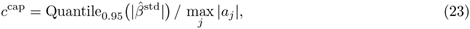

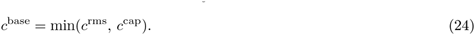

The RMS reference matches the typical standardized marginal effect, and the cap prevents pathological loci with a single very large |*a_j_*| from inducing prior means that exceed the largest plausible standardized effect. The grid of prior-mean scales used in the ensemble is expressed in these units, with ***µ***_0_*_,j_* = *c* · *c*^base^ · *a_j_* at each grid point.

#### Fitting protocol

At each locus we fit a deterministic ensemble of SuSiNE RSS models on the (*z,* **R***, n*) triple, following the C-CS specification carried forward from the simulation study (Section 4.4.2). The ensemble is an 8 × 8 grid crossing eight values of the prior-mean scale,

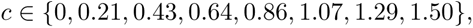

with eight values of the prior variance,

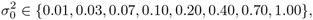

for 64 fits per locus. All fits use *L* = 10 maximum effects, an IBSS iteration cap of 100, convergence tolerance 10*^−^*^5^ on the ELBO, a fixed residual variance, and prior updating disabled so that *c* and 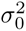 are exactly the values fixed by the grid. The locus-level sample size passed to the RSS likelihood is the median per-variant effective sample size *ñ_l_*; this median is robust to a small number of per-variant QC drop-outs but masks the across-variant heterogeneity in *n_j_* that exists when sample availability is uneven, one of several reasons we treat the real-data analysis as a case study rather than a clinically actionable result (see Discussion). Alongside the ensemble, we fit a single annotation-agnostic anchor using *susieR::susie_rss* [15] at 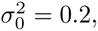 with prior variance estimation off, residual variance fixed at 1, *L* = 10, and the same iteration and tolerance settings; the anchor is the reference fit against which all annotation-driven movement is measured.

### Aggregation and warm refit

Within each locus, the 64 ensemble PIP vectors are aggregated with the same cluster-weight rule carried forward from the simulation study (Section 4.4.2). Complete-linkage hierarchical clustering uses the credible-shift PIP distance

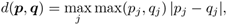

cut at *d* = 0.05. Each cluster is weighted by exp of its largest member ELBO, and the cluster weight is split within the cluster across member fits by ELBO-softmax at temperature 1. The aggregated locus-level PIP vector is the resulting weighted mixture, and the same weights induce the run-level weighting *w*_run_ that drives the annotation-influence diagnostics reported in Section 2.6. Credible sets are built per effect by sorting normalized ***α****_ℓ_* until the cumulative mass reaches 0.95; we report unfiltered sets and compute purity (minimum pairwise |*r*| within set) for diagnostic purposes only.

To distinguish within-basin prior-driven PIP changes from basin movement, we additionally identify each locus’s *highest-weight source run* — the ensemble member with the largest *w*_run_, breaking ties by ELBO and then by run id — and refit it under the annotation-agnostic prior (***µ***_0_ = **0**, 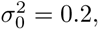 uniform ***π***), warm-started from the source fit’s posterior approximation. This refit asks whether the basin discovered by the annotated fit survives prior removal: if vanilla IBSS, given the annotated basin as a starting point, holds that location without the prior, the basin is durable and the annotation gain represents an exploration win; if vanilla slides back toward the cold anchor, the gain was dependent on the active prior. The construction is the real-data analogue of the warm-vanilla refit used in the high-quality simulation regime (Section 2.3; S1 Appendix, Figs B–C).

#### Diagnostics reported

For each locus we record the standard SuSiE fit-quality outputs (final ELBO, residual-variance trace, credible-set list with sizes and purities), per-variant aggregated PIPs, and per-effect posterior summaries. We also compute four paper-facing diagnostics. *Annotation influence* reports the weighted mean prior-mean scale Σ*_m_ w_m_c_m_* and the off-zero weight share 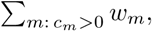 showing whether the ensemble actually uses annotation-informed fits or places its weight back on *c* = 0. *PIP L2 drift versus matched-component drift* compares the highest-weight source and warm refit to the SuSiE anchor, separating changes in variant ranking from movement of the fitted component basis. *Local genetic variance* compares the SuSiE anchor, SuSiNE ensemble, and warm refit using the posterior expected genetic variance, E[var(**X*β***) | *y*]*/*var(*y*), clipped to [0, 1], so that annotation-driven gains in apparent explained signal can be distinguished from pure PIP reshuffling. Finally, *effect diffuseness* bins *K_ℓ_* from Eq. 16 across loci, showing whether annotations sharpen or diffuse individual single-effect components. The selected-locus zoom (Section 2.6) then drills into six loci that span the observed annotation-influence range, reporting six variants per locus selected by absolute baseline-to-ensemble posterior-mean shift alongside each ensemble member’s PIP-vector L2 distance from the anchor as a function of *c*.

#### 4.5.1 Real-data PIP and matched-component drift diagnostics

We give distinct names to the three PIP-vector distances used in the real-data analysis. The ensemble-to-anchor distance is *D*_ens_ = ∥**p**^ensemble^ − **p**^anchor^∥_2_ and appears in Table 4. Fig 8b instead reports the source-run-to-anchor distance *D*_src_ and warm-refit-to-anchor distance *D*_warm_. Both are instances of

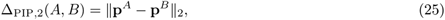

with *B* the anchor and *A* respectively the highest-weight source run or its annotation-free warm refit. We use the same PIP L2 scale for the selected-locus grid traces, so panel-wide and locus-level movements remain comparable.

The horizontal axis is a PVE-weighted matched-component drift. Its purpose is to detect effect-basis movement in real data, where the truth-basis divergence used in simulation cannot be computed. For fit *F*, define each effect’s posterior-mean coefficient vector

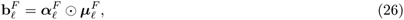

where 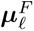 is the posterior mean conditional on inclusion, and define the reference-LD norm 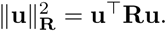

For two fits *A* and *B* at the same locus, components are matched one-to-one by the Hungarian algorithm using the absolute fitted-effect correlation

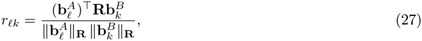

with zero-variance padding as needed. If *σ*(*ℓ*) is the component of *B* matched to component *ℓ* of *A*, the relative signed component drift is

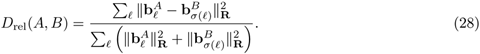

This signed distance catches changes in orientation, sign, and amplitude among matched components, including changes that can cancel when all effects are summed into one total posterior-mean predictor. To keep the paper-facing scale tied to the amount of signal involved, we multiply by the larger whole-model fitted variance of the two compared fits,

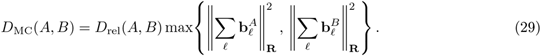

Because the RSS fits use standardized genotypes and standardized phenotype scale, *D*_MC_ is in approximate PVE units. It is deliberately different from both the simulation truth-basis divergence and the older total fitted-value drift. The former requires a known causal basis; the latter can miss component-level rearrangements that cancel in aggregate. In Fig 8b, *A* is either the highest-weight source run or its warm refit, and *B* is the SuSiE anchor.

### 4.6 Ethics statement

This study performed secondary analyses of aggregate GTEx v10 Lung *cis*-eQTL summary statistics and publicly available, pseudonymized 1000 Genomes Project reference-panel genotypes. To estimate linkage disequilibrium, we accessed per-sample genotype calls for EUR reference-panel participants and converted them to locus-level correlation matrices; individual-level genotypes and sample identifiers were not retained in the downstream analysis outputs or redistributed. We did not access individual-level GTEx genotype or expression data, interact with participants, obtain biospecimens, or access directly identifying information. Separate method-evaluation experiments used simulated genotype and phenotype data. The GTEx and 1000 Genomes data were originally collected under the consent and ethics procedures of their source studies. Consistent with Penn State policy for activities that do not meet the definition of human-subjects research, no separate institutional review board submission was required for these analyses.

### 4.7 Use of artificial intelligence tools

The authors used Claude Opus 4.8 (Anthropic) and ChatGPT 5.5 (OpenAI) to assist with code generation, prose revision, and LaTeX formatting. M.G.C. manually reviewed and, where necessary, revised all AI-assisted code, text, and formatting. The authors verified the resulting analyses and manuscript content and take full responsibility for the final work.

## Data availability

All third-party data used in this study are publicly available. GTEx v10 Lung *cis*-eQTL summary statistics were obtained from the GTEx Portal (https://gtexportal.org; dbGaP accession phs000424) [3, 42].

Linkage-disequilibrium reference panels were computed from the 1000 Genomes Project Phase 3 release on GRCh38, restricted to the EUR samples [31]. The simulated multi-ancestry genotypes underlying the simulation studies were obtained from Harvard Dataverse (https://doi.org/10.7910/DVN/COXHAP) [41]. Signed variant-effect annotations were generated with AlphaGenome through its public API [25]; access to the AlphaGenome model and API is as described by its developers. The exact remote AlphaGenome release/build served during annotation generation was not recorded in the archived run metadata. We therefore archive the derived signed-annotation prior-mean tables (below) so that all downstream fits are reproducible even if the served model changes.

The SuSiNE and *test.susine* R packages and the *eQTL-annotations-for-susine* data-preparation code are available on GitHub (https://github.com/Michael-G-Callahan/susine, https://github.com/Michael-G-Callahan/test-susine, https://github.com/Michael-G-Callahan/eQTL-annotations-for-susine) and each is archived at citable, version-specific Zenodo DOIs: SuSiNE software v0.1.0 (https://doi.org/10.5281/zenodo.21687306), the analysis and reproducibility harness v0.1.0 (https://doi.org/10.5281/zenodo.21687310), and the GTEx/AlphaGenome annotation pipeline v0.1.1 (https://doi.org/10.5281/zenodo.21690319). The derived data required to reproduce every figure and reported number are archived separately in a citable Zenodo deposit (DOI: https://doi.org/10.5281/zenodo.21539486). The derived-data archive contains the consolidated simulation outputs (baseline, ensemble-scaling, drift, and sensitivity analyses), the real-data collector outputs for the 20-locus GTEx Lung case study, the per-locus summary-statistic bundles (*z*, LD matrix *R*, and effective sample size *n*), and the AlphaGenome-derived signed-annotation prior-mean (***µ***_0_) tables. Raw genotypes and GTEx summary statistics are not redistributed; they are available from the sources cited above. The derived data are released under CC-BY-4.0 and the software under the MIT License. The analyses depend on *susieR*, used unmodified from its public upstream repository (https://github.com/stephenslab/susieR): the SuSiE-pathology exhibits use commit *34a4a52* (v0.15.51) and all other analyses use tag *0.15.53*.

## Acknowledgments

This research used data from the Genotype-Tissue Expression (GTEx) Project. It also used data from the 1000 Genomes Project. We thank the AlphaGenome team at Google DeepMind for providing API access that enabled the generation of signed variant-effect annotations used in this study. Computational analyses were performed on the Roar Collab high-performance computing cluster at The Pennsylvania State University.

## Supporting information

**S1 Appendix. Supplementary theory, diagnostics, and LD-consistency tables.** Mathematical derivations and proofs for SuSiNE, supplementary real-data diagnostic figures, and LD-consistency tables for the GTEx Lung case study.

## References

1. Visscher PM, Wray NR, Zhang Q, Sklar P, McCarthy MI, Brown MA, et al. 10 Years of GWAS Discovery: Biology, Function, and Translation. American Journal of Human Genetics. 2017;101(1):5–22. doi:10.1016/j.ajhg.2017.06.005.

2. Buniello A, MacArthur JAL, Cerezo M, Harris LW, Hayhurst J, Malangone C, et al. The NHGRI-EBI GWAS Catalog of Published Genome-Wide Association Studies, Targeted Arrays and Summary Statistics 2019. Nucleic Acids Research. 2019;47(D1):D1005–12. doi:10.1093/nar/gky1120.

3. GTEx Consortium. The GTEx Consortium Atlas of Genetic Regulatory Effects across Human Tissues. Science. 2020;369(6509):1318–30. doi:10.1126/science.aaz1776.

4. Schaid DJ, Chen W, Larson NB. From Genome-Wide Associations to Candidate Causal Variants by Statistical Fine-Mapping. Nature Reviews Genetics. 2018;19(8):491–504. doi:10.1038/s41576-018-0016-z.

5. Natarajan BK. Sparse Approximate Solutions to Linear Systems. SIAM Journal on Computing. 1995;24(2):227–34. doi:10.1137/S0097539792240406.

6. Nemhauser GL, Wolsey LA, Fisher ML. An Analysis of Approximations for Maximizing Submodular Set Functions–I. Mathematical Programming. 1978;14(1):265–94. doi:10.1007/BF01588971.

7. Das A, Kempe D. Algorithms for Subset Selection in Linear Regression. In: Proceedings of the 40th Annual ACM Symposium on Theory of Computing; 2008. p. 45–54. doi:10.1145/1374376.1374384.

8. Das A, Kempe D. Approximate Submodularity and Its Applications: Subset Selection, Sparse Approximation and Dictionary Selection. Journal of Machine Learning Research. 2018;19:1–34.

9. O’Hara RB, Sillanpaa MJ. A Review of Bayesian Variable Selection Methods: What, How and Which. Bayesian Analysis. 2009;4(1):85–117. doi:10.1214/09-BA403.

10. Hoeting JA, Madigan D, Raftery AE, Volinsky CT. Bayesian Model Averaging: A Tutorial. Statistical Science. 1999;14(4):382–417. doi:10.1214/ss/1009212519.

11. Hormozdiari F, Kostem E, Kang EY, Pasaniuc B, Eskin E. Identifying Causal Variants at Loci with Multiple Signals of Association. Genetics. 2014;198(2):497–508. doi:10.1534/genetics.114.167908.

12. Benner C, Spencer CCA, Havulinna AS, Salomaa V, Ripatti S, Pirinen M. FINEMAP: Efficient Variable Selection Using Summary Data from Genome-Wide Association Studies. Bioinformatics. 2016;32(10):1493–501. doi:10.1093/bioinformatics/btw018.

13. Wen X, Lee Y, Luca F, Pique-Regi R. Efficient Integrative Multi-SNP Association Analysis via Deterministic Approximation of Posteriors. American Journal of Human Genetics. 2016;98(6):1114–29. doi:10.1016/j.ajhg.2016.03.029.

14. Yang Z, Wang C, Liu L, Khan A, Lee A, Vardarajan B, et al. CARMA Is a New Bayesian Model for Fine-Mapping in Genome-Wide Association Meta-Analyses. Nature Genetics. 2023;55:1057–65. doi:10.1038/s41588-023-01392-0.

15. Wang G, Sarkar A, Carbonetto P, Stephens M. A Simple New Approach to Variable Selection in Regression, with Application to Genetic Fine Mapping. Journal of the Royal Statistical Society: Series B. 2020;82(5):1273–300. doi:10.1111/rssb.12388.

16. Zou Y, Carbonetto P, Wang G, Stephens M. Fine-Mapping from Summary Data with the “Sum of Single Effects” Model. PLoS Genetics. 2022;18(7):e1010299. doi:10.1371/journal.pgen.1010299.

17. Weissbrod O, Hormozdiari F, Benner C, Cui R, Ulirsch J, Gazal S, et al. Functionally Informed Fine-Mapping and Polygenic Localization of Complex Trait Heritability. Nature Genetics. 2020;52(12):1355–63. doi:10.1038/s41588-020-00735-5.

18. Jiang J, Cole JB, Freebern E, Da Y, VanRaden PM, Ma L. Functional Annotation and Bayesian Fine-Mapping Reveals Candidate Genes for Important Agronomic Traits in Holstein Bulls. Communications Biology. 2019;2:212. doi:10.1038/s42003-019-0454-y.

19. Wang QS, Kelley DR, Ulirsch J, Kanai M, Sadhuka S, Cui R, et al. Leveraging Supervised Learning for Functionally Informed Fine-Mapping of cis-eQTLs Identifies an Additional 20,913 Putative Causal eQTLs. Nature Communications. 2021;12:3394. doi:10.1038/s41467-021-23134-8.

20. Srivastava D, Korsakova A, Wang Q, Ruiz L, Yuan H, Kelley DR. Borzoi-Informed Fine Mapping Improves Causal Variant Prioritization in Complex Trait GWAS; 2025. BioRxiv preprint. doi:10.1101/2025.07.09.663936.

21. Reshef YA, Finucane HK, Kelley DR, Gusev A, Kotliar D, Ulirsch JC, et al. Detecting Genome-Wide Directional Effects of Transcription Factor Binding on Polygenic Disease Risk. Nature Genetics. 2018;50:1483–93. doi:10.1038/s41588-018-0196-7.

22. Avsec Z, Agarwal V, Visentin D, Ledsam JR, Grabska-Barwinska A, Taylor KR, et al. Effective Gene Expression Prediction from Sequence by Integrating Long-Range Interactions. Nature Methods. 2021;18(10):1196–203. doi:10.1038/s41592-021-01252-x.

23. Linder J, Srivastava D, Yuan H, Agarwal V, Kelley DR. Predicting RNA-Seq Coverage from DNA Sequence as a Unifying Model of Gene Regulation. Nature Genetics. 2025;57:949–61. doi:10.1038/s41588-024-02053-6.

24. Lal A, Karollus A, Gunsalus L, Garfield D, Nair S, Tseng AM, et al. Decoding Sequence Determinants of Gene Expression in Diverse Cellular and Disease States. Nature Methods. 2026;23:1138–51. doi:10.1038/s41592-026-03102-0.

25. Avsec Ž, Latysheva N, Cheng J, Novati G, Taylor KR, Ward T, et al. Advancing Regulatory Variant Effect Prediction with AlphaGenome. Nature. 2026;649(8099):1206–18. doi:10.1038/s41586-025-10014-0.

26. Wakefield J. Bayes Factors for Genome-Wide Association Studies: Comparison with P-Values. Genetic Epidemiology. 2009;33(1):79–86. doi:10.1002/gepi.20359.

27. Cui R, Elzur RA, Kanai M, Ulirsch JC, Weissbrod O, Daly MJ, et al. Improving Fine-Mapping by Modeling Infinitesimal Effects. Nature Genetics. 2024;56(1):162–9. doi:10.1038/s41588-023-01597-3.

28. McCreight A, Cho Y, Li R, Nachun D, Gan HY, Carbonetto P, et al. SuSiE 2.0: Improved Methods and Implementations for Genetic Fine-Mapping and Phenotype Prediction; 2025. BioRxiv preprint. doi:10.1101/2025.11.25.690514.

29. Yao Y, Vehtari A, Simpson D, Gelman A. Using Stacking to Average Bayesian Predictive Distributions. Bayesian Analysis. 2018;13(3):917–1007. doi:10.1214/17-BA1091.

30. Yao Y, Vehtari A, Gelman A. Stacking for Non-Mixing Bayesian Computations: The Curse and Blessing of Multimodal Posteriors. Journal of Machine Learning Research. 2022;23:1–45.

31. 1000 Genomes Project Consortium. A Global Reference for Human Genetic Variation. Nature. 2015;526(7571):68–74. doi:10.1038/nature15393.

32. Denault WRP, Carbonetto P, Li R, Alzheimer’s Disease Functional Genomics Consortium, Wang G, Stephens M. Accounting for Uncertainty in Residual Variances Improves Calibration for Fine-Mapping with Small Sample Sizes; 2025. BioRxiv preprint. doi:10.1101/2025.05.16.654543.

33. Guo F, Wang X, Fan K, Broderick T, Dunson DB. Boosting Variational Inference; 2016. ArXiv:1611.05559. Available from: https://arxiv.org/abs/1611.05559. doi:10.48550/arXiv.1611.05559.

34. Miller AC, Foti NJ, Adams RP. Variational Boosting: Iteratively Refining Posterior Approximations. In: Proceedings of the 34th International Conference on Machine Learning. vol. 70 of Proceedings of Machine Learning Research. PMLR; 2017. p. 2420–9. Available from: https://proceedings.mlr.press/v70/miller17a.html.

35. Zou Y, Carbonetto P, Xie D, Wang G, Stephens M. Fast and Flexible Joint Fine-Mapping of Multiple Traits via the Sum of Single Effects Model. Nature Genetics. 2026;58(2):454–62. doi:10.1038/s41588-025-02486-7.

36. Zhao S, Crouse W, Qian S, Luo K, Stephens M, He X. Adjusting for Genetic Confounders in Transcriptome-Wide Association Studies Improves Discovery of Risk Genes of Complex Traits. Nature Genetics. 2024;56:336–47. doi:10.1038/s41588-023-01648-9.

37. Zhu X, Duren Z, Wong WH. Modeling Regulatory Network Topology Improves Genome-Wide Analyses of Complex Human Traits. Nature Communications. 2021;12:2851. doi:10.1038/s41467-021-22588-0.

38. van de Wiel MA, Te Beest DE, Munch MM. Learning from a Lot: Empirical Bayes for High-Dimensional Model-Based Prediction. Scandinavian Journal of Statistics. 2019;46(1):2–25. doi:10.1111/sjos.12335.

39. Beal MJ. Variational Algorithms for Approximate Bayesian Inference [PhD thesis]. Gatsby Computational Neuroscience Unit, University College London; 2003.

40. Beal MJ, Ghahramani Z. Variational Bayesian Learning of Directed Graphical Models with Hidden Variables. Bayesian Analysis. 2006;1(4):793–831. doi:10.1214/06-BA126.

41. Zhang H. Simulated Data for 600,000 Subjects from Five Ancestries [dataset on the Internet]. Harvard Dataverse; 2022. Available from: 10.7910/DVN/COXHAP. Version 5.3.

42. GTEx Consortium. GTEx Portal, Version 10 (V10) Data Release [dataset on the Internet]; 2024. Available from: https://gtexportal.org. Lung cis-eQTL all-pairs association data; dbGaP accession phs000424.

