## Supplementary material for "SuSiNE: Genetic fine-mapping with signed functional priors and multi-basin ensembling": S1 Appendix

#### Theory, diagnostics, and LD-consistency tables

### Abstract

This appendix provides mathematical foundations for Sum of Single Non-central Effects (SuSiNE), an extension of the SuSiE fine-mapping model that incorporates signed functional priors. We establish how SuSiE's theoretical results extend to SuSiNE, derive the evidence lower bound (ELBO), characterize optimal hyperparameter settings under a stylized generative model, and identify limitations in single-locus hyperparameter estimation. The general optimal annotation scale is the causal-set regression slope of the true effect on the annotation; it equals the squared annotation–effect correlation only under the unit-loading generator used for the stylized analysis. We also provide conditions under which SuSiNE's potential benefits are realized, showing that sparse genetic architectures pose fundamental challenges for ranking-based metrics regardless of annotation quality. Additional empirical diagnostics and LD-consistency tables are included at the end of the appendix.

**9 Sparsity Throttles AUPRC Gains** **14**

**10 Convergence of IBSS for SuSiNE** **16**

**11 The Identification Problem** **17**

**14 Additional tables** **19**

### 1 Toy pathology constructions and complete numerical results

The four examples are deliberately pathological mechanism demonstrations. They are designed to distinguish ordinary LD ambiguity, the representational limitation of one factorized variational fit, and initialization-sensitive optimization. They are not intended to mimic the geometry of a typical eQTL locus. Each dataset has $n = 600$ individuals and $p = 50$ variants. Columns 1–6 carry all engineered structure. The causal support is $\{1, 3, 5\}$ and the decoy support is $\{2, 4, 6\}$. The remaining 44 columns are null. Each dataset is generated under SuSiE's own sparse linear Gaussian assumptions, without interactions or nonlinear effects. A failure in these examples therefore does not require an additional form of model misspecification.

For every scenario, we fit SuSiE with $L = 5$ with initialized parameters from the package default, a truth-warm start, and a decoy-warm start. The two components beyond the three causal signals leave SuSiE room to resolve additional structure if the data support it. Residual and prior variances use the package's default empirical-Bayes updates. The prior variance is initialized at `scaled_prior_variance` $= 0.2$ and is then updated rather than held fixed.

The pattern across starts is part of the diagnostic. Agreement across all starts indicates ambiguity that one factorized fit can express. Distinct configurations with nearly equal ELBOs indicate uncertainty that one factorized fit cannot retain jointly. Distinct configurations with a large ELBO gap indicate an optimizer barrier. We later carry these signatures into realistic simulations and real data, where they serve as diagnostic evidence rather than exact replicas of the toy geometry.

**Scenario 1 (control).** The 50 variants are approximately independent. Causal effects at columns 1, 3, and 5 are 0.8, $-0.8$, and 1.0, and target PVE is 0.3. There is no LD, representation, or optimizer barrier in this setting. All three initializations give the same fit. PIP is approximately one on each causal variant, the final ELBO is $-1392.79$, and fitted PVE is approximately 0.25.

**Scenario 2 (LD ambiguity and purity).** Each causal variant is paired with a decoy that has the same realized marginal association with the phenotype. The outer pairs $(1, 2)$ and $(5, 6)$ have $r \approx 0.98$. The middle pair $(3, 4)$ has only $r = 0.10$. The phenotype is built from three orthogonal signal components with target variance fractions 0.09, 0.12, and 0.05. After construction, the causal variants have realized marginal PVEs of 8.9%, 6.6%, and 5.0%, respectively, and each is matched by its decoy. These effect sizes fall in the strongest tier of the realistic oligogenic architecture used in the main simulations. The intended posterior structure is

$$\mathrm{AND}\{\mathrm{OR}(1, 2), \mathrm{OR}(3, 4), \mathrm{OR}(5, 6)\}.$$

All starts converge to the same fit at ELBO $-799.21$ and fitted PVE about 0.17. SuSiE splits PIP approximately $(0.50, 0.50)$ within every pair. The association evidence in this realized example does not distinguish the low-correlation middle pair at the resolution of its fitted single-effect component. That component explains about 3.5% of phenotypic variance even though its credible-set purity is 0.10. Because $L = 5$, two components remain idle beyond the three fitted signals. If the data contained evidence that separated the middle pair, the model would have spare components available to use it. The unfiltered 50-variant causal-recovery AUPRC is 0.50. Removing the two idle diffuse components leaves AUPRC unchanged. A purity-0.50 filter instead removes the real middle signal and lowers AUPRC to 0.35.

**Scenario 3 (factorized-representation barrier).** The causal triplet $\mathcal{A} = \{X_1, X_3, X_5\}$ is an orthonormal basis of a three-dimensional subspace. The decoy triplet $\mathcal{B} = \{X_2, X_4, X_6\}$ is obtained with the minimax rotation $\mathbf{Q} = \mathbf{I}_3 - (2/3)\mathbf{1}\mathbf{1}^\top$. Consequently, $\mathrm{span}(\mathcal{A}) = \mathrm{span}(\mathcal{B})$ while the maximum cross-set pairwise correlation is 2/3. The remaining 44 null columns are projected to be orthogonal to this shared subspace. Phenotypes use effects $(1, -1.1, 0.5)$ on $\mathcal{A}$ at target PVE 0.4.

The causal basis, the decoy basis, and mixed bases can reproduce nearly the same fitted mean. The posterior uncertainty is therefore an OR across coordinated AND configurations rather than a set of independent within-component choices. The default, truth-warm, and decoy-warm fits reach mixed, causal, and decoy configurations with ELBOs $-1256.68$, $-1256.82$, and $-1256.70$, respectively. Their ELBO-softmax weights are $(0.35, 0.31, 0.34)$. Each fit can be highly confident within one configuration, but one product-over-effects fit cannot retain the three configurations jointly.

**Scenario 4 (optimizer barrier).** Causal columns 1, 3, and 5 have mutual correlation about 0.98. Coefficients $(1.5, -2.2, 0.7)$ produce partial sign cancellation in their marginal associations. The decoys are progressively noisier proxies of the joint causal signal. Each decoy has a larger marginal correlation with the phenotype than any causal variant, reaching values up to 0.34 compared with at most 0.11 for the causal variants. Their joint explanatory power is nevertheless smaller. Target PVE is 0.98.

The default and decoy-warm starts reach the decoy basin at ELBO $-807.72$ and fitted PVE 0.18. The truth-warm start reaches the causal basin at ELBO 294.69 and fitted PVE 0.98. The truth start is not deployable because it uses unavailable causal information. Its role is to show that coordinate ascent can miss a much better basin when the first marginal screen favors jointly inferior decoys.

| Scenario | Intended diagnostic | Initialization behavior | Main numerical result |
|---|---|---|---|
| 1 | control | all starts agree | ELBO $-1392.79$ |
| 2 | LD ambiguity / purity | all starts agree | AUPRC 0.50 $\rightarrow$ 0.35 after purity filtering |
| 3 | representation barrier | three near-tied fits | ELBO range 0.14 nats |
| 4 | optimizer barrier | truth and decoy basins | fitted PVE 0.18 $\rightarrow$ 0.98 |

**Table A.** Complete toy-pathology summary. These examples isolate mechanisms and are not intended to mimic the geometry of a typical eQTL locus.

### 2 Annotation calibration and scale interpretation

The simulation annotation generator has two quality parameters. On causal variants, $\phi_a$ is the squared correlation between the annotation and the true effect. On null variants, $\nu_a$ controls the centered second moment relative to the causal annotation scale. The causal annotation is constructed as

$$a_{\mathcal{C}} = s_{\mathcal{C}} \left\{ \sqrt{\phi_a}\, u_\beta + \sqrt{1-\phi_a}\, u_\epsilon \right\},$$

where $u_\beta$ and $u_\epsilon$ are centered unit-RMS signal and orthogonal noise vectors, and $s_{\mathcal{C}}$ is the centered RMS of the causal effects. This construction fixes the centered RMS of the causal annotation block relative to the causal effects while $\phi_a$ changes its signal-to-noise mixture. It does not hold the total annotation RMS fixed after the null block is added. Null annotations are centered Gaussian noise whose second moment is scaled relative to the causal annotation block according to $\nu_a$. The two parameters therefore describe different aspects of annotation quality. The first measures directional accuracy on causal variants. The second measures how much annotation magnitude is assigned to variants that are truly null.

**Calibration target and published benchmarks.** Neither $\phi_a$ nor $\nu_a$ maps directly to one published AlphaGenome metric. We therefore use AlphaGenome's GTEx eQTL benchmarks to anchor a realistic simulation regime. We do not claim that this generator reproduces the full AlphaGenome error distribution.

AlphaGenome reports three benchmarks that are relevant here. Its eQTL-causality task distinguishes high-confidence fine-mapped variants with SuSiE PIP at least 0.9 from distance-matched controls with PIP below 0.01. The reported zero-shot auROC is 0.71. Its eQTL-coefficient task compares AlphaGenome scores with SuSiE posterior effect sizes across 17,675 fine-mapped `(variant, gene, tissue)` tuples. The reported signed main-scatter Spearman correlation is 0.50, the unsigned correlation is 0.10, and the tissue-weighted mean signed performance is 0.49. We use the causality auROC and signed coefficient correlation as anchors. The unsigned coefficient result is retained as a diagnostic because the signed prior mean uses information that an unsigned prior-inclusion channel discards.

**Simulator analogues.** We evaluate 600 phenotype datasets derived from 150 genotype matrices over an $11 \times 11$ grid with $(\phi_a, \nu_a) \in \{0, 0.1, \ldots, 1\}^2$. The causality analogue treats the three largest true causal effects per dataset as positives and true null variants as negatives. AlphaGenome instead uses distance-matched low-PIP controls, so the simulator reproduces the scoring idea rather than the exact label construction. The coefficient analogues use all 23 true causal variants. Entries are pooled across datasets before scoring.

The sparse-versus-null AUPRC has prevalence approximately $3/(3+977)$. Its numerical value is therefore not directly comparable with an auROC from a balanced evaluation. We use an equal-variance Gaussian score model with $S_0 \sim N(0,1)$ and $S_1 \sim N(d',1)$. At the empirical prevalence, each $d'$ induces one precision-recall curve and one AUPRC. We numerically invert this mapping and report the balanced-case equivalent

$$\text{auROC}_{\text{eq}} = \Phi(d'/\sqrt{2}).$$

Round-trip checks at balanced auROCs 0.50, 0.60, 0.70, and 0.80 recover their input values to floating-point tolerance.

**Calibration outcome.** The equivalent causality auROC is governed mainly by $\nu_a$ because reducing null annotation magnitude separates causal and null variants. Signed coefficient correlation is governed mainly by $\phi_a$ because it directly controls the signal-to-noise mixture on causal variants. Matching the two anchor benchmarks selects $\nu_a \approx 0.95$ for a causality auROC near 0.71 and $\phi_a \approx 0.3$ for signed $\rho \approx 0.49$. The baseline screen uses the nearest coarse-grid point $(0.3, 0.9)$. The ensemble study fixes $\nu_a = 0.95$.

The generator cannot match the signed and unsigned coefficient benchmarks simultaneously. Matching the signed correlation produces more unsigned rank information than AlphaGenome reports. This may favor functional-$\pi$, which uses $|\boldsymbol{a}|$. The signed-Gaussian construction may also favor the prior-mean channel because its error model resembles the model's own prior. We therefore treat the calibration as a bridge to a realistic annotation regime, not as a claim that the comparison is conservative or that the simulator recreates AlphaGenome predictions exactly.

**Why an empirical grid value near $c = 0.64$ is not the theoretical value 0.3.** At the baseline working point, the best saved single-fit grid value is $c = 0.642857\ldots$. This coefficient multiplies the simulator's explicitly scaled annotation. In the real-data workflow, the units differ again: bounded AlphaGenome quantiles are inverse-normal transformed, clipped to $[-2.5, 2.5]$, and divided by their within-locus RMS without mean-centering. A locus-specific base scale then anchors the prior mean to standardized marginal-effect units.

The general per-SNP optimum derived in Section 8 is

$$c^\star = \frac{\text{Cov}(\beta, a \mid \mathcal{C})}{\text{Var}(a \mid \mathcal{C})},$$

which is a unit-dependent regression slope. Rescaling $a$ by a positive factor $s$ changes the equivalent coefficient to $c/s$ while leaving $c\boldsymbol{a}$ and the scale-free correlation $\phi_a$ unchanged. The equality $c^\star = \phi_a$ holds only for the unit-loading generator $a = \beta + \epsilon$. Annotation rescaling is therefore the primary reason an empirical grid value near 0.64 need not equal $\phi_a = 0.3$. PIP softmax coupling, LD competition, optimizing AUPRC rather than expected per-SNP log Bayes factor, and interaction with $\sigma_0^2$ are additional differences between the theorem and the empirical grid.

### 3 Preliminaries: From SuSiE to SuSiNE

This section establishes how the theoretical foundations of SuSiE [1] extend to SuSiNE. We state the key SuSiE results and indicate which carry directly to SuSiNE and which require modification.

#### 3.1 The SuSiE Model and Algorithm

The SuSiE model represents the regression coefficient vector $\mathbf{b} \in \mathbb{R}^p$ as a sum of $L$ "single-effect" vectors:

$$\mathbf{y} = \mathbf{X}\mathbf{b} + \mathbf{e}, \quad \mathbf{e} \sim N_n(\mathbf{0}, \sigma^2 \mathbf{I}_n), \tag{1}$$

$$\mathbf{b} = \sum_{l=1}^{L} \mathbf{b}_l, \tag{2}$$

$$\mathbf{b}_l = \boldsymbol{\gamma}_l \odot b_l, \quad \boldsymbol{\gamma}_l \sim \text{Mult}(1, \boldsymbol{\pi}), \quad b_l \sim N(0, \sigma_0^2), \tag{3}$$

where $\boldsymbol{\gamma}_l$ is a one-hot vector selecting which SNP carries effect $l$, and $\boldsymbol{\pi} = (\pi_1, \ldots, \pi_p)$ is the prior inclusion probability vector.

> **Key Insight:** Each $\mathbf{b}_l$ has exactly one non-zero entry, making the single-effect regression (SER) model analytically tractable. SuSiE exploits this by iteratively fitting SER models to residuals.

#### 3.2 SuSiNE: The Non-Central Extension

SuSiNE modifies the prior on $b_l$ to have non-zero mean:

$$b_l | \gamma_{lj} = 1 \sim N(\mu_{0j}, \sigma_0^2), \tag{4}$$

where $\mu_{0j} = c \cdot a_j$ incorporates directional information from annotations $\mathbf{a} \in \mathbb{R}^p$ scaled by parameter $c \geq 0$.

> **Takeaway:** When $c = 0$, SuSiNE reduces to SuSiE. The parameter $c$ controls how strongly we weight the directional information from annotations.

#### 3.3 Theoretical Results: What Carries Over

We now state how each major SuSiE result extends to SuSiNE.

**Proposition 3.1** (Variational Factorization—Unchanged)**.** *SuSiNE uses the same variational family as SuSiE:*

$$q(\mathbf{b}_1, \ldots, \mathbf{b}_L) = \prod_{l=1}^{L} q_l(\mathbf{b}_l). \tag{5}$$

*This factorization assumption is unchanged by the non-zero prior mean.*

**Proposition 3.2** (Coordinate Ascent Structure—Unchanged)**.** *The optimal $q_l$ given $\{q_{l'}\}_{l' \neq l}$ is obtained by fitting a single-effect regression model to the expected residuals $\bar{\mathbf{r}}_l = \mathbf{y} - \mathbf{X} \sum_{l' \neq l} \bar{\mathbf{b}}_{l'}$.*

*This is Proposition 1 of Wang et al. (2020). The proof depends only on the additive structure of the model, not on the prior mean, and thus carries directly to SuSiNE.*

**Proposition 3.3** (IBSS as Coordinate Ascent—Unchanged)**.** *The Iterative Bayesian Stepwise Selection (IBSS) algorithm is a coordinate ascent algorithm for maximizing the ELBO. This is Corollary 1 of Wang et al. (2020) and follows immediately from Proposition 3.2.*

**Proposition 3.4** (Monotonicity and stationary accumulation points)**.** *For fixed $0 < \sigma, \sigma_0 < \infty$ and $\pi_j > 0$ for all $j$, each exact IBSS coordinate update for SuSiNE does not decrease the ELBO. Consequently, the ELBO values converge to a finite limit. Under the regularity conditions invoked in Section 10, every accumulation point of the IBSS iterates is stationary for the ELBO.*

*Proof.* See Section 10 for the complete proof. □

**Proposition 3.5** (Exchangeability—Modified)**.** *Wang et al. (2020, Proposition 3) show that for identical SNPs $\mathbf{x}_j = \mathbf{x}_k$ with identical priors $\pi_j = \pi_k$, the posterior is exchangeable in $j$ and $k$.*

*For SuSiNE, this extends to: exchangeability holds when $\mathbf{x}_j = \mathbf{x}_k$, $\pi_j = \pi_k$, **and** $\mu_{0j} = \mu_{0k}$. The additional condition reflects that different annotations break the symmetry between otherwise identical SNPs.*

**Proposition 3.6** (Prior Equivalence—Unchanged)**.** *Wang et al. (2020, Proposition A2) show that as $p \to \infty$ with $L$ fixed, the SuSiE prior converges to a standard BVSR prior with $L$ effects. This carries directly to SuSiNE, with the limiting prior being BVSR with non-zero prior means.*

##### 3.4 Posterior representations: AND-of-ORs and OR-of-ANDs

The main paper uses an informal pair of terms — *AND-of-ORs* and *OR-of-ANDs* — to describe how different fine-mapping methods represent posterior uncertainty over which configurations of variants are jointly causal. This subsection makes those terms precise.

**Definition 3.7** (Support and configuration)**.** A *configuration* is a subset $S \subseteq \{1, \ldots, p\}$ of variant indices, identified with its 0/1 indicator $\mathbf{1}_S \in \{0,1\}^p$. The set of all configurations is the Boolean lattice $2^{[p]}$.

The fine-mapping posterior of interest assigns a probability $\pi(S)$ to each configuration $S$. Different methods make different parametric assumptions about which distributions $\pi$ over $2^{[p]}$ they can represent. We characterize two representational classes that capture the essential dichotomy.

**Definition 3.8** (AND-of-ORs family)**.** The *AND-of-ORs* family with $L$ components consists of distributions over $2^{[p]}$ that admit a factorization

$$\pi(S) \;=\; \sum_{\substack{j_1,\ldots,j_L \in [p] \\ \{j_1,\ldots,j_L\} = S}} \prod_{\ell=1}^{L} \alpha_{\ell, j_\ell}, \tag{6}$$

for some collection of probability vectors $\boldsymbol{\alpha}_\ell = (\alpha_{\ell,1}, \ldots, \alpha_{\ell,p}) \in \Delta^{p-1}$, $\ell = 1, \ldots, L$. Equivalently, $\pi$ is the law of $\{J_1, \ldots, J_L\}$ where the $J_\ell$ are independent draws with $J_\ell \sim \boldsymbol{\alpha}_\ell$ (an "AND" across $L$ independent "OR" choices, allowing repeats which collapse to a smaller support).

**Definition 3.9** (OR-of-ANDs family)**.** The *OR-of-ANDs* family consists of arbitrary discrete distributions over $2^{[p]}$, viewed as mixtures over support sets:

$$\pi(S) \;=\; \sum_{T \in 2^{[p]}} w_T \, \mathbb{1}[S = T], \qquad w_T \geq 0, \; \sum_T w_T = 1. \tag{7}$$

The OR-of-ANDs family is the full simplex over $2^{[p]}$ and can represent any posterior; it is the representational target of explicit-enumeration methods such as FINEMAP, CAVIAR, and DAP-G. The AND-of-ORs family is the much smaller parametric family used by the SuSiE/SuSiNE variational posterior, in which the per-effect inclusion vectors $\boldsymbol{\alpha}_\ell$ are the natural parameters.

**Proposition 3.10** (Strict containment)**.** *For $p \geq 4$ and $L \geq 2$, the AND-of-ORs family with $L$ components is a strict subset of the OR-of-ANDs family. In particular, there exist distributions $\pi$ over $2^{[p]}$ that cannot be written in the form* (6) *for any choice of $\boldsymbol{\alpha}_1, \ldots, \boldsymbol{\alpha}_L$.*

*Sketch.* A counter-example with $p = 4$, $L = 2$ suffices. Consider $\pi$ that places mass $\frac{1}{2}$ on $S_1 = \{1, 3\}$ and $\frac{1}{2}$ on $S_2 = \{2, 4\}$, and mass 0 everywhere else. Under the AND-of-ORs form (6), the marginal inclusion probability of variant 1 is $\alpha_{1,1} + \alpha_{2,1} - \alpha_{1,1}\alpha_{2,1}$ (and so on). Marginalizing $\pi$ to its single-variant inclusion probabilities yields $(1/2, 1/2, 1/2, 1/2)$, which any AND-of-ORs distribution can reproduce by suitable choice of $\boldsymbol{\alpha}_\ell$. However, the AND-of-ORs distribution that achieves this also places mass on $\{1, 4\}$ and $\{2, 3\}$ (with $\alpha_{1,1}\alpha_{2,4}, \alpha_{1,2}\alpha_{2,3} > 0$ under the required marginals), contradicting $\pi(\{1,4\}) = \pi(\{2,3\}) = 0$. The two configurations $S_1, S_2$ are therefore correlated under $\pi$ in a way no product distribution can encode. □

**Remark 3.11** (Why the dichotomy matters for fine-mapping)**.** The example in Proposition 3.10 is the canonical *model-specification barrier* discussed in the main paper: two genuinely competing causal configurations that explain the data equally well cannot be co-represented by a SuSiE/SuSiNE single fit because the variational posterior is in the AND-of-ORs family. The remedy is not to enlarge the variational family (which would forfeit the tractability that makes SuSiE/SuSiNE useful) but to aggregate multiple fits, each landing in a different AND-of-ORs basin, into a mixture that lives in the OR-of-ANDs family. The cluster-weight aggregator (main paper, Methods: Ensembling and aggregation) is exactly such an OR-of-ANDs mixture over AND-of-ORs basins.

### 4 Notation and Setup

#### 4.1 Basic Notation

Throughout, we use the following notation:

- $n$: sample size
- $p$: number of SNPs
- $L$: number of single effects in the model
- $\mathbf{y} \in \mathbb{R}^n$: response vector (phenotype)
- $\mathbf{X} \in \mathbb{R}^{n \times p}$: genotype matrix (columns centered and scaled)
- $\boldsymbol{\beta} \in \mathbb{R}^p$: true effect sizes (sparse)
- $\sigma^2$: residual variance
- $\sigma_0^2$: prior variance on effect sizes
- $\boldsymbol{\mu}_0 \in \mathbb{R}^p$: prior mean vector (zero for SuSiE, non-zero for SuSiNE)
- $\mathbf{a} \in \mathbb{R}^p$: annotation vector from sequence-to-function model
- $c \geq 0$: annotation scale parameter (so $\mu_{0j} = c \cdot a_j$)
- $\boldsymbol{\pi} \in \mathbb{R}^p$: prior inclusion probability vector for the Multinomial

#### 4.2 Derived Quantities

For each SNP $j$:

- $\hat{b}_j = (\mathbf{x}_j^\top \mathbf{x}_j)^{-1} \mathbf{x}_j^\top \mathbf{y}$: univariate least squares estimate
- $s_j^2 = \sigma^2 / (\mathbf{x}_j^\top \mathbf{x}_j)$: sampling variance of $\hat{b}_j$
- $z_j = \hat{b}_j / s_j$: $z$-score
- $\tau_j^2 = \sigma_0^2 / s_j^2$: prior-to-noise variance ratio
- $\kappa_j = \tau_j^2 / (1 + \tau_j^2)$: shrinkage factor (weight on data in posterior mean)
- $\tilde{a}_j = a_j / s_j$: annotation in $z$-score units
- $\tilde{\mu}_{0j} = \mu_{0j} / s_j = c \cdot \tilde{a}_j$: prior mean in $z$-score units

**Remark 4.1** (Interpretation of $\kappa_j$)**.** The quantity $\kappa_j \in (0, 1)$ measures how much the posterior mean weights the data versus the prior. When $\kappa_j \approx 1$ (large $\tau_j^2$), the posterior is dominated by the data. When $\kappa_j \approx 0$ (small $\tau_j^2$), the posterior is dominated by the prior.

**Remark 4.2** (Homogeneity on standardized data)**.** When genotypes are standardized (each column of $\mathbf{X}$ has unit variance), we have $\mathbf{x}_j^\top \mathbf{x}_j = n$ for all $j$, giving $s_j^2 = \sigma^2 / n$ uniformly. Consequently, $\tau_j^2$ and $\kappa_j$ are constant across SNPs. This homogeneity assumption, which holds in standard fine-mapping practice, simplifies several results below and is explicitly invoked where needed.

#### 4.3 Causal Architecture

**Definition 4.3** (Causal Set)**.** The causal set is $\mathcal{C} = \{j : \beta_j \neq 0\}$ with cardinality $|\mathcal{C}| = p^*$.

**Definition 4.4** (Causal Fraction)**.** The causal fraction is $\rho = p^*/p \in (0, 1)$.

**Definition 4.5** (Annotation Strength)**.** We assume annotations are *pre-oriented.* Thus,

$$\mathrm{Corr}(a_j, \beta_j \mid j \in \mathcal{C}) \geq 0.$$

The annotation strength is:

$$\phi_a = \mathrm{Corr}(a_j, \beta_j \mid j \in \mathcal{C})^2 \in [0, 1]. \tag{8}$$

When $\phi_a = 1$, annotations perfectly predict effect sizes among causal SNPs; when $\phi_a = 0$, annotations are uninformative.

The squared-correlation definition is dimensionless and scale-invariant in both $\boldsymbol{a}$ and $\boldsymbol{\beta}$, so $\phi_a$ is well-defined regardless of the units in which the annotation is reported. The additive-noise generative model used in Section 7, $a_j = \beta_j + \epsilon_j^{(1)}$ for $j \in \mathcal{C}$, implicitly fixes a common scale for $\boldsymbol{a}$ and $\boldsymbol{\beta}$. If the annotation is reported on another scale, as is typical for sequence-to-function model outputs, it can be rescaled by any positive constant without changing $\phi_a$, while the per-locus scalar $c$ in $\boldsymbol{\mu}_0 = c\boldsymbol{a}$ absorbs the rescaling. The real-data analysis makes this explicit by RMS-normalizing $\boldsymbol{a}$ within each locus and anchoring $c$ to the locus's standardized marginal effect scale.

**Remark 4.6** (Pre-orientation assumption)**.** The pre-orientation assumption ensures that $c \geq 0$ is the natural parameter space. In practice, if sequence-to-function model predictions are systematically anti-correlated with true effects (e.g., due to predicting the wrong strand), the annotation vector should be negated before applying SuSiNE. The quantity $\phi_a$ measures the *strength* of the annotation-effect relationship, not its direction.

> **Takeaway:** Annotation strength $\phi_a$ is defined only over causal SNPs and measures how well annotations predict true effect magnitudes (given correct orientation). This is the key quantity determining SuSiNE's potential benefit over SuSiE.

### 5 Derivation of the ELBO

This section derives the evidence lower bound for SuSiNE. The derivation follows Wang et al. (2020) exactly—the non-zero prior mean affects only the specific form of the single-effect Bayes factors, not the ELBO structure.

#### 5.1 The Additive Effects Model

The additive effects model underlying SuSiE/SuSiNE is:

$$\mathbf{y} = \sum_{l=1}^{L} \boldsymbol{\mu}_l + \mathbf{e}, \quad \mathbf{e} \sim N_n(\mathbf{0}, \sigma^2 \mathbf{I}_n), \tag{9}$$

where $\boldsymbol{\mu}_l \in \mathbb{R}^n$ are additive effect vectors with independent priors $\boldsymbol{\mu}_l \sim g_l$.

For SuSiNE, $\boldsymbol{\mu}_l = \mathbf{X}\mathbf{b}_l$ where $\mathbf{b}_l$ follows a single-effect regression prior with non-zero mean.

#### 5.2 ELBO Decomposition

**Lemma 5.1** (ELBO Form)**.** *Under the factorized variational approximation*

$$q(\boldsymbol{\mu}_1, \ldots, \boldsymbol{\mu}_L) = \prod_l q_l(\boldsymbol{\mu}_l),$$

*the ELBO equals:*

$$F(q, g, \sigma^2; \mathbf{y}) = -\frac{n}{2} \log(2\pi\sigma^2) - \frac{1}{2\sigma^2} ERSS + \sum_{l=1}^{L} \mathbb{E}_{q_l} \left[ \log \frac{g_l(\boldsymbol{\mu}_l)}{q_l(\boldsymbol{\mu}_l)} \right], \tag{10}$$

*where the expected residual sum of squares is:*

$$ERSS = \left\| \mathbf{y} - \sum_{l=1}^{L} \bar{\boldsymbol{\mu}}_l \right\|^2 + \sum_{l=1}^{L} \sum_{i=1}^{n} Var[\mu_{li}], \tag{11}$$

*with $\bar{\boldsymbol{\mu}}_l = \mathbb{E}_{q_l}[\boldsymbol{\mu}_l]$.*

*Proof.* This is equation (B.6) in Wang et al. (2020, Supplement). The derivation uses only the Gaussian likelihood and independence across $l$ under $q$, neither of which depends on the prior mean. See the original supplement for details. □

> **Takeaway:** The ELBO structure is identical for SuSiE and SuSiNE. The non-zero prior mean affects the model only through the Bayes factors that appear in the third term.

### 6 Single Effect Regression with Non-Zero Prior Mean

This section derives the posterior distribution and Bayes factor for the SuSiNE single-effect model. These are the key quantities that differ from SuSiE.

#### 6.1 The SuSiNE Single Effect Model

For effect $l$ with expected residual $\bar{\mathbf{r}}_l = \mathbf{y} - \sum_{l' \neq l} \bar{\boldsymbol{\mu}}_{l'}$, the SuSiNE single effect model is:

$$\bar{\mathbf{r}}_l = \mathbf{X}\mathbf{b}_l + \mathbf{e}, \quad \mathbf{e} \sim N_n(\mathbf{0}, \sigma^2 \mathbf{I}_n), \tag{12}$$

$$\mathbf{b}_l = \boldsymbol{\gamma}_l \odot b_l \quad \text{(elementwise)}, \tag{13}$$

$$\boldsymbol{\gamma}_l \sim \text{Mult}(1, \boldsymbol{\pi}), \tag{14}$$

$$b_l | \gamma_{lj} = 1 \sim N(\mu_{0j}, \sigma_0^2), \tag{15}$$

where $\boldsymbol{\gamma}_l$ is a one-hot vector selecting which SNP carries the effect.

#### 6.2 Posterior Distribution

**Lemma 6.1** (SuSiNE Posterior)**.** *Conditioning on $\gamma_{lj} = 1$ (SNP $j$ carries the effect), the posterior for the effect size $b$ is $N(\mu_{1j}, \sigma_{1j}^2)$ where:*

$$\sigma_{1j}^2 = \frac{s_j^2 \sigma_0^2}{s_j^2 + \sigma_0^2}, \tag{16}$$

$$\mu_{1j} = \kappa_j \hat{b}_j + (1 - \kappa_j)\mu_{0j}. \tag{17}$$

*Proof.* Standard conjugate Gaussian analysis. The likelihood contribution from SNP $j$ is proportional to $\exp\{-(b - \hat{b}_j)^2/(2s_j^2)\}$. The prior is proportional to $\exp\{-(b - \mu_{0j})^2/(2\sigma_0^2)\}$. Multiplying and completing the square yields the result. □

> **Key Insight:** Equation (17) shows the posterior mean is a precision-weighted average of the data ($\hat{b}_j$) and the prior mean ($\mu_{0j}$), with weight $\kappa_j$ on the data. This is the standard Bayesian updating formula for Gaussian models.

#### 6.3 Bayes Factor

The Bayes factor is the key quantity determining posterior inclusion probabilities.

**Lemma 6.2** (SuSiNE Bayes Factor)**.** *The Bayes factor for SNP $j$ comparing $\gamma_{lj} = 1$ to $b_j = 0$ is:*

$$\log BF_j = -\frac{1}{2}\log(1 + \tau_j^2) + \frac{\kappa_j z_j^2}{2} + (1 - \kappa_j) z_j \tilde{\mu}_{0j} - \frac{(1 - \kappa_j)\tilde{\mu}_{0j}^2}{2}. \tag{18}$$

*Proof.* The Bayes factor compares the marginal likelihood under $H_1 : b \sim N(\mu_{0j}, \sigma_0^2)$ to $H_0 : b = 0$.

Under $H_1$: $\hat{b}_j | b \sim N(b, s_j^2)$ and $b \sim N(\mu_{0j}, \sigma_0^2)$, so by convolution of Gaussians, $\hat{b}_j \sim N(\mu_{0j}, s_j^2 + \sigma_0^2)$.

Under $H_0$: $\hat{b}_j \sim N(0, s_j^2)$.

The log Bayes factor is:

$$\log \mathrm{BF}_j = \log \frac{N(\hat{b}_j; \mu_{0j}, s_j^2 + \sigma_0^2)}{N(\hat{b}_j; 0, s_j^2)} \tag{19}$$

$$= -\frac{1}{2} \log \left( \frac{s_j^2 + \sigma_0^2}{s_j^2} \right) - \frac{(\hat{b}_j - \mu_{0j})^2}{2(s_j^2 + \sigma_0^2)} + \frac{\hat{b}_j^2}{2s_j^2}. \tag{20}$$

Substituting $\hat{b}_j = z_j s_j$, $\mu_{0j} = \tilde{\mu}_{0j} s_j$, and $\tau_j^2 = \sigma_0^2 / s_j^2$:

$$\log \mathrm{BF}_j = -\frac{1}{2} \log(1 + \tau_j^2) - \frac{(z_j - \tilde{\mu}_{0j})^2}{2(1 + \tau_j^2)} + \frac{z_j^2}{2}. \tag{21}$$

Expanding $(z_j - \tilde{\mu}_{0j})^2 = z_j^2 - 2z_j\tilde{\mu}_{0j} + \tilde{\mu}_{0j}^2$:

$$\log \mathrm{BF}_j = -\frac{1}{2} \log(1 + \tau_j^2) + \frac{z_j^2}{2} \left( 1 - \frac{1}{1 + \tau_j^2} \right) + \frac{z_j \tilde{\mu}_{0j}}{1 + \tau_j^2} - \frac{\tilde{\mu}_{0j}^2}{2(1 + \tau_j^2)}. \tag{22}$$

Using $1 - 1/(1 + \tau_j^2) = \kappa_j$ and $1/(1 + \tau_j^2) = 1 - \kappa_j$ yields (18). □

**Lemma 6.3** (Bayes Factor Decomposition)**.** *The log Bayes factor decomposes as:*

$$\log BF_j = \log BF_j^{SuSiE} + \underbrace{c(1 - \kappa_j) z_j \tilde{a}_j}_{alignment} - \underbrace{\frac{c^2(1 - \kappa_j)\tilde{a}_j^2}{2}}_{regularization}, \tag{23}$$

*where* $\log BF_j^{SuSiE} = -\frac{1}{2} \log(1 + \tau_j^2) + \frac{\kappa_j z_j^2}{2}$ *is the standard SuSiE Bayes factor.*

*Proof.* Substitute $\tilde{\mu}_{0j} = c\tilde{a}_j$ into (18) and group terms. □

**Key Insight:** The decomposition (23) reveals two effects of the directional prior:

1. **Alignment term** $c(1 - \kappa_j) z_j \tilde{a}_j$: Increases BF when $z_j$ and $\tilde{a}_j$ have the same sign (data agrees with annotation), decreases BF when they disagree.
2. **Regularization term** $-\frac{c^2(1-\kappa_j)\tilde{a}_j^2}{2}$: Always negative, penalizing non-zero prior means. This prevents unbounded BF inflation.

The factor $(1 - \kappa_j)$ weights these terms by how much the posterior relies on the prior.

**Remark 6.4** (Sign of Regularization Term)**.** The regularization term is *always negative* for any $\tau_j^2 > 0$ and $c > 0$. This provides automatic protection against over-reliance on annotations.

### 7 Expected Log Bayes Factor Analysis

This section analyzes the expected Bayes factor under a generative model for effects and annotations. The goal is to understand when SuSiNE systematically outperforms SuSiE *in expectation.*

**Remark 7.1** (Scope of this analysis)**.** The results in Sections 7–9 analyze *expected* quantities under a stylized generative model with key simplifying assumptions: (i) weak LD / approximately independent SNPs, (ii) homogeneous $\kappa_j$ and $s_j^2$ across SNPs, and (iii) per-SNP analysis that ignores softmax coupling across PIPs. These assumptions are explicitly stated and should be borne in mind when interpreting results. In particular, ranking-based metrics like AUPRC depend on distribution *tails*, not just means, and the coupling structure in real fine-mapping can cause behavior that deviates from these per-SNP predictions.

#### 7.1 Data-Generating Process

**Assumption 7.2** (Observation Model)**.** The $z$-scores satisfy:

$$z_j = \frac{\beta_j}{s_j} + \xi_j, \quad \xi_j \overset{\text{iid}}{\sim} N(0,1), \tag{24}$$

where the noise terms $\xi_j$ are independent across SNPs (weak LD approximation).

**Assumption 7.3** (Annotation Model)**.** The annotations satisfy:

$$a_j = \begin{cases} \beta_j + \epsilon_j^{(1)}, & j \in \mathcal{C}, \quad \epsilon_j^{(1)} \sim N(0, \sigma_{a,1}^2), \\ \epsilon_j^{(0)}, & j \notin \mathcal{C}, \quad \epsilon_j^{(0)} \sim N(0, \sigma_{a,0}^2), \end{cases} \tag{25}$$

where $\epsilon_j^{(1)}$, $\epsilon_j^{(0)}$, and $\xi_j$ are mutually independent, and $\beta_j \overset{\text{iid}}{\sim} N(0, \sigma_\beta^2)$ for $j \in \mathcal{C}$.

**Remark 7.4.** Under Assumption 7.3, the annotation strength is:

$$\phi_a = \frac{\sigma_\beta^2}{\sigma_\beta^2 + \sigma_{a,1}^2}. \tag{26}$$

Perfect annotations ($\sigma_{a,1}^2 = 0$) give $\phi_a = 1$; pure noise ($\sigma_{a,1}^2 \to \infty$) gives $\phi_a = 0$.

#### 7.2 Expected Bayes Factor Components

**Lemma 7.5** (Expected Alignment and Regularization)**.** *Under Assumptions 7.2–7.3 with the homogeneity approximation $s_j \approx \bar{s}$:*
***(a) For causal SNPs ($j \in \mathcal{C}$):***

$$\mathbb{E}[z_j \tilde{a}_j \mid j \in \mathcal{C}] = \frac{\sigma_\beta^2}{\bar{s}^2}, \tag{27}$$

$$\mathbb{E}[\tilde{a}_j^2 \mid j \in \mathcal{C}] = \frac{\sigma_\beta^2 + \sigma_{a,1}^2}{\bar{s}^2}. \tag{28}$$

***(b) For non-causal SNPs ($j \notin \mathcal{C}$):***

$$\mathbb{E}[z_j \tilde{a}_j \mid j \notin \mathcal{C}] = 0, \tag{29}$$

$$\mathbb{E}[\tilde{a}_j^2 \mid j \notin \mathcal{C}] = \frac{\sigma_{a,0}^2}{\bar{s}^2}. \tag{30}$$

*Proof.* **Part (a):** For $j \in \mathcal{C}$:

$$\mathbb{E}[z_j \tilde{a}_j] = \mathbb{E}\left[\left(\frac{\beta_j}{\bar{s}} + \xi_j\right) \cdot \frac{\beta_j + \epsilon_j^{(1)}}{\bar{s}}\right] \tag{31}$$

$$= \frac{1}{\bar{s}^2}\mathbb{E}[\beta_j^2 + \beta_j \epsilon_j^{(1)} + \bar{s}\xi_j\beta_j + \bar{s}\xi_j\epsilon_j^{(1)}] \tag{32}$$

$$= \frac{\sigma_\beta^2}{\bar{s}^2}, \tag{33}$$

since all cross terms vanish by independence. Similarly, $\mathbb{E}[a_j^2] = \mathbb{E}[(\beta_j + \epsilon_j^{(1)})^2] = \sigma_\beta^2 + \sigma_{a,1}^2$.

**Part (b):** For $j \notin \mathcal{C}$: $z_j = \xi_j$ and $a_j = \epsilon_j^{(0)}$ are independent mean-zero, so $\mathbb{E}[z_j \tilde{a}_j] = 0$. ☐

**Takeaway:** When annotations track true effects, the alignment term has positive expectation for causal SNPs and zero expectation for non-causal SNPs. This asymmetry is the source of SuSiNE's potential advantage.

**Lemma 7.6** (Expected Log Bayes Factor Difference)**.** *Define* $\Delta_j(c) = \mathbb{E}[\log BF_j(c)] - \mathbb{E}[\log BF_j(0)]$ *as the expected gain from using SuSiNE with scale c versus SuSiE. Under homogeneity* ($\kappa_j \approx \bar{\kappa}$, $s_j \approx \bar{s}$)*:*
***(a) For causal SNPs:***

$$\bar{\Delta}^{(\mathcal{C})}(c) = c(1-\bar{\kappa})\frac{\sigma_\beta^2}{\bar{s}^2} - \frac{c^2(1-\bar{\kappa})(\sigma_\beta^2 + \sigma_{a,1}^2)}{2\bar{s}^2}. \tag{34}$$

***(b) For non-causal SNPs:***

$$\bar{\Delta}^{(\mathcal{N})}(c) = -\frac{c^2(1-\bar{\kappa})\sigma_{a,0}^2}{2\bar{s}^2}. \tag{35}$$

*Proof.* Substitute the expectations from Lemma 7.5 into the decomposition (23) and note that $\mathbb{E}[\log \mathrm{BF}_j^{\mathrm{SuSiE}}]$ does not depend on $c$. □

> **Key Insight:** Equation (34) shows that causal SNPs can gain from SuSiNE (when $c$ is chosen appropriately), while equation (35) shows that non-causal SNPs are always penalized in expectation. This is the desired behavior: SuSiNE should boost causal signals without inflating null signals *on average*.

### 8 Optimal Annotation Scaling

This section derives the optimal annotation scale $c^\star$ and makes explicit its dependence on annotation units. The unit-loading generator of Assumption 7.3 yields a particularly simple special case.

#### 8.1 Single-SNP Optimum

**Proposition 8.1** (Optimal annotation scale for expected causal-SNP Bayes factor)**.** *Assume that* $(\beta_j, a_j) \mid j \in \mathcal{C}$ *has finite second moments, is centered, and that the shrinkage factor is constant over causal variants (or is independent of* $(\beta_j, a_j)$*). If* $\mathrm{Var}(a_j \mid j \in \mathcal{C}) > 0$*, the annotation scale that maximizes* $\mathbb{E}[\log \mathrm{BF}_j \mid j \in \mathcal{C}]$ *is*

$$c^\star = \frac{\mathrm{Cov}(\beta_j, a_j \mid j \in \mathcal{C})}{\mathrm{Var}(a_j \mid j \in \mathcal{C})}. \tag{36}$$

*Under Assumption 7.3,* $a_j = \beta_j + \epsilon_j^{(1)}$ *with independent mean-zero noise, this reduces to* $c^\star = \sigma_\beta^2/(\sigma_\beta^2 + \sigma_{a,1}^2) = \phi_a$.

*Proof.* Up to the positive constant $(1-\bar{\kappa})/\bar{s}^2$, the $c$-dependent part of the expected log Bayes factor is

$$g(c) = c\,\mathrm{Cov}(\beta_j, a_j \mid j \in \mathcal{C}) - \tfrac{1}{2}c^2\,\mathrm{Var}(a_j \mid j \in \mathcal{C}).$$

Hence $g'(c) = \mathrm{Cov}(\beta_j, a_j \mid \mathcal{C}) - c\,\mathrm{Var}(a_j \mid \mathcal{C})$ and $g''(c) = -\mathrm{Var}(a_j \mid \mathcal{C}) < 0$. Substitution of the unit-loading generator gives the stated special case. If the parameter space is restricted to $c \geq 0$ without pre-orienting the annotation, the constrained optimum is the positive part of Eq. (36). □

> **Key Insight:** The optimal numerical scale is a regression slope and therefore depends on annotation units. Annotation strength $\phi_a$ is scale-invariant; it equals $c^\star$ only under the unit-loading normalization of the stylized generator.

**Remark 8.2** (Independence from $\tau_j^2$)**.** Under the homogeneity or independence conditions in Proposition 8.1, the optimal $c^\star$ does not depend on $\tau_j^2$ or $\kappa_j$. These parameters affect the *magnitude* of the gain from using SuSiNE, but not the maximizing regression slope.

#### 8.2 Maximum Expected Gain

**Corollary 8.3** (Maximum Gain for Causal SNPs)**.** *Under the unit-loading generator of Assumption 7.3, at the special-case optimum $c^\star = \phi_a$, the maximum expected gain for a causal SNP is:*

$$\bar{\Delta}^{(\mathcal{C})}(c^\star) = \frac{(1-\bar{\kappa})\sigma_\beta^2\phi_a}{2\bar{s}^2} = \frac{(1-\bar{\kappa})\sigma_\beta^4}{2\bar{s}^2(\sigma_\beta^2+\sigma_{a,1}^2)}. \tag{37}$$

*Proof.* Substitute $c^\star = \phi_a$ into (34) and simplify. □

**Takeaway:** The maximum expected gain is proportional to $\phi_a$ (annotation strength) and $(1-\bar{\kappa})$ (weight on prior). When annotations are perfect ($\phi_a = 1$), the gain is maximized. When the data dominate ($\bar{\kappa} \to 1$), the prior matters less and the gain diminishes.

#### 8.3 Regret from Suboptimal $c$

Under the unit-loading generator, $c^\star = \phi_a$ depends on the latent causal set. We quantify the resulting loss from a suboptimal scale within that stylized model.

**Proposition 8.4** (Regret Bound for Causal SNPs)**.** *For a causal SNP $j \in \mathcal{C}$, the regret (loss in expected log BF) from using $\hat{c}$ instead of $c^\star = \phi_a$ is:*

$$\mathit{Regret}^{(\mathcal{C})}(\hat{c}) := \bar{\Delta}^{(\mathcal{C})}(c^\star) - \bar{\Delta}^{(\mathcal{C})}(\hat{c}) = \frac{(1-\bar{\kappa})(\sigma_\beta^2+\sigma_{a,1}^2)}{2\bar{s}^2}\cdot(\phi_a-\hat{c})^2. \tag{38}$$

*Proof.* Define $A = (1-\bar{\kappa})\sigma_\beta^2/\bar{s}^2$ and $B = (1-\bar{\kappa})(\sigma_\beta^2+\sigma_{a,1}^2)/(2\bar{s}^2)$. Then from (34):

$$\bar{\Delta}^{(\mathcal{C})}(c) = Ac - Bc^2. \tag{39}$$

This is a downward parabola maximized at $c^\star = A/(2B) = \phi_a$, with maximum value $\bar{\Delta}^{(\mathcal{C})}(c^\star) = A^2/(4B)$. For any $\hat{c}$:

$$\mathrm{Regret}^{(\mathcal{C})}(\hat{c}) = \frac{A^2}{4B} - (A\hat{c} - B\hat{c}^2) = B(\phi_a-\hat{c})^2. \tag{40}$$

□

**Key Insight:** The regret is *quadratic* in the estimation error $\phi_a - \hat{c}$: it is flat near the optimum and symmetric in over- and under-estimation, so a coarse grid over $c$ is comparatively forgiving in the neighborhood of $\phi_a$. The absolute penalty still scales with the locus-specific curvature constant $B$, which depends on unknown nuisance parameters, so the claim is relative (per unit $B$) rather than absolute.

### 9 Sparsity Throttles AUPRC Gains

SuSiNE's benefit from directional priors depends on genetic architecture through the causal fraction $\rho = p^*/p$. The main-text simulations demonstrate AUPRC gains in an oligogenic regime; the analysis below shows why those gains are increasingly throttled as the architecture becomes sparser, to the point where a directional prior need not help at all. The earlier sections analyzed expected ELBO changes aggregated over all SNPs, an objective dominated by the $(p-p^*)$ null SNPs that does not directly correspond to ranking-based metrics such as AUPRC; here we work directly with the precision–recall relationship.

The starting point is that higher-strength annotations increase the *expected causal–null separation* in per-SNP evidence scores. By Lemma 7.5, the directional increment $\Delta_j(c) = c(1-\bar{\kappa})z_j\tilde{a}_j - \frac{1}{2}c^2(1-\bar{\kappa})\tilde{a}_j^2$ has positive expectation for causal SNPs and non-positive expectation for null SNPs, so—provided $c$ is not so large that the quadratic regularization dominates—causal scores shift up relative to null scores, and this gap is nondecreasing in annotation strength $\phi_a$. The difficulty is that AUPRC depends on the *tails* of the null score distribution, not on this mean separation: the alignment term $z_j \tilde{a}_j$ has zero mean but positive variance for null SNPs, so some nulls receive spurious boosts by chance, and in sparse regimes these rare events dominate precision.

AUPRC depends on the causal fraction $\rho = p^*/p$ through the precision–recall relationship. Let $S_j$ denote any continuous ranking score (e.g., $\log \alpha_j$ or $\log \mathrm{BF}_j$). Let $F_C$ and $F_N$ be the CDFs of $S_j$ under causal and null SNPs respectively.

For a threshold $t$, define:

$$R(t) := \Pr(S_j \geq t \mid j \in \mathcal{C}) = 1 - F_C(t)$$

(recall at threshold $t$) and

$$U(t) := \Pr(S_j \geq t \mid j \notin \mathcal{C}) = 1 - F_N(t)$$

(null tail probability at threshold $t$).

**Proposition 9.1** (Sparsity Throttles Precision and AUPRC)**.** *For any threshold $t$, precision is:*

$$\mathit{Prec}(t) = \Pr(j \in \mathcal{C} \mid S_j \geq t) = \frac{\rho \cdot R(t)}{\rho \cdot R(t) + (1-\rho) \cdot U(t)}. \tag{41}$$

*Equivalently, parameterizing by recall $r \in (0,1)$ with $t(r) = F_C^{-1}(1-r)$ and $u(r) = U(t(r))$:*

$$\mathit{Prec}(r) = \frac{1}{1 + \frac{1-\rho}{\rho} \cdot \frac{u(r)}{r}}. \tag{42}$$

*Consequently:*

1. *For any fixed pair $(F_C, F_N)$, Prec$(r)$ and AUPRC are increasing in $\rho$.*
2. *To achieve precision at least $\eta \in (0,1)$ at recall $r$, it is necessary that:*

$$u(r) \leq \frac{\rho}{1-\rho} \cdot \frac{1-\eta}{\eta} \cdot r. \tag{43}$$

*In sparse regimes ($\rho \ll 1$), achieving nontrivial precision requires $u(r) = O(\rho)$.*

*Proof.* Bayes' rule gives (41): among all SNPs exceeding threshold $t$, the expected fraction causal is proportional to $\rho$ and $(1-\rho)$ multiplied by their respective tail probabilities. Reparameterization by recall yields (42). Monotonicity in $\rho$ follows since (42) is decreasing in $(1-\rho)/\rho$. Finally, rearranging gives (43). □

**Remark 9.2** (Why expected-value gains need not translate to AUPRC in sparse regimes)**.** Combining the two ingredients: better annotations increase expected causal–null separation, which tends to reduce $u(r)$ at a given recall, but Proposition 9.1 shows that when $\rho \ll 1$ even modest null tail probabilities dominate the precision denominator and suppress AUPRC. The theory therefore predicts *when gains should be largest* (moderate $\rho$, high $\phi_a$) rather than guaranteeing gains in the sparsest regimes, where softmax coupling across PIPs—ignored in this per-SNP analysis—can further let a spuriously boosted null steal mass from nearby causals. This is consistent with the main-text sparse-architecture sensitivity check: the C-CS AUPRC gain remained positive across the tested sparse grid but was smaller than in the central oligogenic benchmark, with high-precision recall often nearly unchanged.

**Corollary 9.3** (Gaussian Separation Scaling)**.** *Consider a stylized model where $S \mid N \sim N(0,1)$ and $S \mid C \sim N(\mu, 1)$. The null tail at recall $r$ is $u(r) = 1 - \Phi(\mu + \Phi^{-1}(1-r))$. To achieve precision $\eta$ at recall $r$,* (43) *requires:*

$$\mu \gtrsim \Phi^{-1}\left(1 - \frac{\rho}{1-\rho} \cdot \frac{1-\eta}{\eta} \cdot r\right) - \Phi^{-1}(1-r),$$

*which grows on the order of $\sqrt{2\log(1/\rho)}$ as $\rho \to 0$ (by Gaussian tail inversion). Thus, increasingly sparse settings require substantially larger causal–null separation to obtain comparable AUPRC.*

### 10 Convergence of IBSS for SuSiNE

This section verifies monotone ELBO ascent and characterizes stationary accumulation points for SuSiNE IBSS, following the coordinate-ascent argument of Wang et al. (2020, Proposition 2).

**Proposition 10.1** (Monotonicity and stationary accumulation points of SuSiNE IBSS)**.** *For fixed $0 < \sigma, \sigma_0 < \infty$ and $\pi_j > 0$ for all $j = 1, \dots, p$, exact cyclic IBSS updates produce a non-decreasing sequence of ELBO values that converges to a finite limit. Under the regularity conditions of the cyclic block-coordinate-ascent result used below, every accumulation point of the iterate sequence is stationary for the ELBO $F$.*

*Proof.* We follow the argument of Wang et al. (2020, Supplement B.5.2), which establishes convergence of IBSS for SuSiE, and verify that the non-zero prior mean preserves each condition it requires. The argument has two ingredients: IBSS is monotone coordinate ascent on an objective bounded above, and the optimal update for each coordinate block is a uniquely determined variational factor.

**Step 1: IBSS is monotone ascent on a bounded objective.** By Proposition 3.2, updating $q_l$ while holding $\{q_{l'}\}_{l' \neq l}$ fixed maximizes the ELBO $F$ over the factor $q_l$, and is equivalent to fitting an SER model with non-zero prior mean to the expected residuals $\bar{\mathbf{r}}_l$. Each update therefore does not decrease $F$, so the sequence $\{F(q^{(t)})\}_t$ is non-decreasing. Because $F$ is a lower bound on the (finite) log marginal likelihood, it is bounded above, and hence $\{F(q^{(t)})\}_t$ converges.

**Step 2: The optimal factor $q_l$ is uniquely determined.** The maximizer of $F$ over $q_l$ is the exact posterior under the SER model with prior $b|\gamma_j = 1 \sim N(\mu_{0j}, \sigma_0^2)$ and likelihood $\hat{b}_j | b \sim N(b, s_j^2)$, namely

$$\sigma_{1j}^2 = \left( \frac{1}{s_j^2} + \frac{1}{\sigma_0^2} \right)^{-1}, \tag{44}$$

$$\mu_{1j} = \sigma_{1j}^2 \left( \frac{\hat{b}_j}{s_j^2} + \frac{\mu_{0j}}{\sigma_0^2} \right), \tag{45}$$

$$\alpha_j \propto \pi_j \cdot \mathrm{BF}_j, \tag{46}$$

with $\mathrm{BF}_j$ given by Lemma 6.2. For $0 < \sigma, \sigma_0 < \infty$ and $\pi_j > 0$, these moments and inclusion weights are uniquely determined by $(\bar{\mathbf{r}}_l, \sigma^2, \sigma_0^2, \boldsymbol{\mu}_0, \boldsymbol{\pi})$, so the optimal $q_l$ is a unique distribution. We emphasize that the uniqueness required here is of the variational *factor* $q_l$, not of which variant carries the effect: when two variants are statistically exchangeable the optimal factor splits inclusion mass between them ($\alpha_j = \alpha_k$), but this is still a single, uniquely determined distribution and hence a single maximizer of $F$ over $q_l$. The familiar SuSiE behavior of spreading PIP mass across an LD-equivalent set is a property of that unique factor, not a violation of uniqueness.

**Step 3: Stationarity of accumulation points.** Subject to the continuity and compact-level-set regularity conditions of Proposition 2.7.1 of Bertsekas (1999), the unique exact block maximizers imply that every accumulation point of the iterate sequence is stationary for $F$. This argument establishes convergence of the ELBO values and stationarity of accumulation points; it does not by itself establish convergence of the entire iterate sequence to a unique point. The non-zero prior mean $\mu_{0j}$ enters only through the SER posterior moments in Step 2 and leaves the monotonicity, boundedness, and factor-uniqueness arguments otherwise unchanged. □

**Remark 10.2** (Comparison with SuSiE)**.** The proof structure follows Wang et al. (2020, Supplement B.5.2). The non-zero prior mean $\mu_{0j}$ shifts the SER posterior mean but changes neither the uniqueness of the optimal variational factor nor the monotone-ascent and boundedness properties on which convergence rests. In particular, the factor-level uniqueness in Step 2 is what the convergence argument requires; ambiguity over *which* of several exchangeable variants is credited is a property of that unique factor and does not create multiple coordinate maximizers.

### 11 The Identification Problem

The previous sections showed that the optimal $c^\star$ is a causal-set regression slope, with $c^\star = \phi_a$ under the unit-loading generator. This section states the information limitation involved in learning that target at a single locus.

#### 11.1 Fundamental Non-Identifiability

**Proposition 11.1** (Single-locus identification limitation)**.** *Without labeled causal variants, auxiliary restrictions on the distributions of causal and null effects, or information shared across loci, observed data from one locus do not in general identify the causal-set regression slope* $\mathrm{Cov}(\beta, a \mid \mathcal{C})/\,\mathrm{Var}(a \mid \mathcal{C})$*, and hence do not identify its unit-loading special case* $\phi_a$*.*

*Proof.* The target is conditional on the latent causal set. By Definition 4.5,

$$\phi_a = \mathrm{Corr}(a_j, \beta_j \mid j \in \mathcal{C})^2. \tag{47}$$

This requires knowing which SNPs are causal. The observable correlation $r = \mathrm{Corr}(a_j, z_j)$ computed over all SNPs conflates:

1. True signal correlation among $j \in \mathcal{C}$
2. Noise correlation among $j \notin \mathcal{C}$ (zero in expectation but with variance)
3. The unknown mixing proportion $\rho = p^*/p$

Without the restrictions listed in the proposition, different latent allocations of variants to causal and null groups and different annotation–effect relationships can produce the same observable mixture. The all-variant correlation therefore does not in general disentangle these components or recover the causal-set target. □

> **Key Insight:** This creates a circularity: we need $\phi_a$ to set $c^\star$, but we need to know which SNPs are causal to estimate $\phi_a$. Yet identifying causal SNPs is precisely the goal of fine-mapping.

**Remark 11.2** (Pooling across loci)**.** While $\phi_a$ is not identifiable from a single locus, pooling information across many loci with shared annotation quality may enable estimation of a global $\phi_a$ (or equivalently, $c^\star$). This is an avenue for future work but is not addressed in this paper.

#### 11.2 Circularity in Empirical Bayes Estimation

**Remark 11.3** (Circular Dependence)**.** Consider estimating $c$ by maximizing the ELBO:

$$\hat{c} = \arg\max_c F(q(c), c, \sigma_0^2; \mathbf{y}). \tag{48}$$

The optimal $q(c)$ depends on $c$ through the PIPs $\alpha_j(c)$, which determine which SNPs receive weight. But the "correct" $c$ depends on $\phi_a$, which depends on knowing $\mathcal{C}$—precisely what $\alpha_j$ attempts to estimate.

Formally, the EB estimate implicitly solves the fixed-point problem:

$$\hat{c} = c^\star(\hat{\phi}_a), \quad \hat{\phi}_a = \phi_a(\mathcal{C}(\hat{q})), \quad \hat{q} = q(\hat{c}), \tag{49}$$

where $\mathcal{C}(q) = \{j : \alpha_j > \theta\}$ for some threshold $\theta$. This circular dependence makes single-fit EB estimates unreliable.

### 12 Extension to Summary Statistics

SuSiNE extends naturally to summary statistics following the approach of SuSiE-RSS [2].

**Remark 12.1** (SuSiNE-RSS)**.** The SuSiE-RSS likelihood replaces the individual-level data likelihood with an equivalent form based on sufficient statistics: $z$-scores $\hat{\mathbf{z}}$, an LD matrix $\mathbf{R}$, and sample size $n$. For SuSiNE, the key observation is that the Bayes factor formula (Lemma 6.2) depends on data only through $z_j$ and the variance ratio $\tau_j^2$. Both quantities are available from summary statistics:

- $z_j = \hat{z}_j$ directly from the summary statistics
- $\tau_j^2 = n\sigma_0^2$ under standardization (since $s_j^2 = \sigma^2/n$ with standardized genotypes)

The prior mean $\mu_{0j} = c \cdot a_j$ enters identically. Therefore, SuSiNE-RSS is implemented by applying the modified Bayes factor (18) within the IBSS-ss algorithm of Zou et al. (2022), with no additional derivation required.

All theoretical results in this appendix apply equally to the summary statistics setting, with the understanding that LD matrix misspecification introduces additional approximation error as characterized in Zou et al. (2022).

### 13 Additional figures

#### 13.1 Effect-drift and truth-basis-divergence analysis (high-quality regime)

This section gives the full methods and figures for the high-quality-regime effect-drift analysis summarized in the main text (Simulation study results — baseline). Because the $L$ component indices are arbitrary and effects migrate across fits, we compare effect quality across specifications at the level of the *distribution* of $(K_\ell, A_\ell)$ — the effect-diffuseness and accuracy metrics defined in the main text — rather than by tracking individually paired components. Each fitted effect contributes one point on the $(K_\ell, A_\ell)$ plane, weighted by its fraction of fitted-$\hat{\mathbf{y}}$ variance explained; for a given pair of specifications we estimate the variance-weighted two-dimensional density of effects under each and report the absolute difference (destination minus source), so that regions where effects accumulate or vacate are read directly off the plane. Comparisons use the high-annotation-quality condition $(\phi_a, \nu_a) = (0.5, 0.9)$ to maximize contrast in effect behavior. To isolate optimizer-barrier from model-specification-barrier contributions, we run a 5-arm refit chain: cold SuSiE-vanilla; cold SuSiNE-functional-$\mu$ at $c = 0.6$; SuSiE-vanilla warm-started from the converged $\mu$ fit; cold SuSiE-functional-$\pi$ at $\tau = 0.464$; and SuSiE-vanilla warm-started from the converged $\pi$ fit. Here "cold" means fitting from the default initialization; the two warm refits remove the annotation prior and use the annotated posterior approximation only as the initialization, asking whether the basin discovered under the prior is durable under the annotation-free objective.

Whereas $K_\ell$ and $A_\ell$ grade effect quality within a single fit, the truth-basis divergence asks whether a treatment moves the fitted explanatory basis toward the known data-generating basis. For a fitted model $F$, denote by $\hat{\mathbf{y}}_\ell^F$ the fitted-$\hat{\mathbf{y}}$ contribution of component $\ell$ and by $v_\ell^F$ its fraction of variance explained. We construct a synthetic truth basis from the simulated causal effects by ranking true causal variants by $|\boldsymbol{\beta}|$ and retaining the top $K^\star = 23$ variants, one component per causal: $\hat{\mathbf{y}}_k^\star = \beta_k \mathbf{X}_{\cdot,k}$, with explained-variance fraction $v_k^\star = \beta_k^2 \operatorname{Var}(\mathbf{X}_{\cdot,k})/\operatorname{Var}(\mathbf{y})$. Fitted and truth components are paired via the maximum-weight one-to-one assignment on the matrix of Pearson correlations $r_{\ell k} = \operatorname{Corr}(\hat{\mathbf{y}}_\ell^F, \hat{\mathbf{y}}_k^\star)$, solved exactly by the Hungarian algorithm with zero-variance padding as needed. The truth-basis divergence is the variance-weighted $1 - r^2$ drift across matched fit-truth pairs:

$$D_{\text{truth}}(F) = \frac{\sum_{\ell=1}^{L} w_\ell \left(1 - r_{\ell,\sigma(\ell)}^2\right)}{\sum_{\ell=1}^{L} w_\ell}, \qquad w_\ell = \min\Big(v_\ell^F,\, v_{\sigma(\ell)}^\star\Big), \tag{50}$$

where $\sigma(\ell)$ is the truth component matched to fitted component $\ell$. The term $1 - r^2$ is the fraction of variance in one matched component left unexplained by the other and ranges from 0 at $r = \pm 1$ to 1 at $r = 0$. The min-variance weighting suppresses pairings involving inactive components or zero-variance padding, so $D_{\text{truth}}$ measures displacement among components carrying signal. The per-dataset *reduction* in truth-basis divergence delivered by a treatment, $D_{\text{truth}}(\text{COLD-VANILLA}) - D_{\text{truth}}(\text{ANNOTATED})$, is the mechanism-level analogue of per-dataset $\Delta$AUPRC.

The pooled precision-recall curves (Fig. BA) show that the annotated $\boldsymbol{\mu}_0$ arm gives the largest practical gain. Pooled AUPRC rises from 0.25 for cold SuSiE to 0.35 for annotated $\boldsymbol{\mu}_0$, versus 0.29 for annotated $\boldsymbol{\pi}$, with warm refits retaining only small gains (0.27 from $\boldsymbol{\mu}_0$, 0.26 from $\boldsymbol{\pi}$). These drift-chain AUPRCs use continuous-threshold integration rather than the pooled confusion bins of the baseline-screen and ensemble headline numbers, so they should be compared within this figure rather than treated as interchangeable with the headline benchmarks. At a fixed 50% precision, cold SuSiE recovers 19% of the top-eight causal variants, annotated $\boldsymbol{\mu}_0$ recovers 31%, and annotated $\boldsymbol{\pi}$ recovers 26%. Per-effect density shifts (Fig. BB) move mass toward the high-accuracy, small-$K_\ell$ corner for both channels, more strongly for $\boldsymbol{\mu}_0$. The cold-to-warm-refit panels show weaker shifts that primarily re-center effects onto causal variants, isolating the durable between-basin operation.

Truth-basis divergence quantifies this between-basin movement at the model level (Fig. C). Cold SuSiE AUPRC declines nonlinearly as the fitted $\widehat{\mathbf{y}}$ lies farther from $\mathbf{X}\boldsymbol{\beta}$ (df-5 spline $R^2 = 0.216$; panel A), and per-dataset reductions in basis divergence track $\Delta$AUPRC (panel B). The cold annotated fits have positive $\Delta$AUPRC intercepts even at zero basis movement (0.087 for $\boldsymbol{\mu}_0$, 0.041 for $\boldsymbol{\pi}$), which is the within-basin sharpening mode. The warm-refit intercepts are approximately zero (0.0014 and $-0.00064$), with consistently positive slopes. Direct $\mu$-versus-$\pi$ comparisons (panels C, D) show $\boldsymbol{\mu}_0$ giving the larger AUPRC gain on 371 datasets before prior removal (tied on 152, smaller on 77), attenuating to 73 / 478 / 49 after warm refit, while its truth-basis-divergence advantage persists (76 then 49 datasets, versus 25 then 16 for $\boldsymbol{\pi}$). Decomposing the mean per-dataset gain (panels E, F), movement along the baseline divergence–AUPRC curve explains about 17% of the cold-annotated gain for either channel (0.016/0.097 for $\boldsymbol{\mu}_0$, 0.0075/0.045 for $\boldsymbol{\pi}$), with the remainder in the positive intercept. For the much smaller warm-refit gains (0.0081 and 0.0031), the basis-predicted component accounts for essentially all of the improvement. Thus the annotation prior resolves both within-basin confusion (erased by prior removal) and between-basin confusion (the optimizer-barrier analogue, durable after prior removal), with $\boldsymbol{\mu}_0$ more effective than $\boldsymbol{\pi}$ at both.

### 14 Additional tables

### Acknowledgments

This appendix supports the article “SuSiNE: Genetic fine-mapping with signed functional priors and multi-basin ensembling” by M. G. Callahan and X. Zhu. The analyses reported here used data from the Genotype-Tissue Expression (GTEx) Project and the 1000 Genomes Project, signed variant-effect annotations generated with AlphaGenome under API access provided by the AlphaGenome team at Google DeepMind, and computing resources on the Roar Collab cluster at The Pennsylvania State University. Competing interests and the full data- and code-availability statement are given in the main article.

### References

1. Wang G, Sarkar A, Carbonetto P, Stephens M. A Simple New Approach to Variable Selection in Regression, with Application to Genetic Fine Mapping. Journal of the Royal Statistical Society: Series B. 2020;82(5):1273-300. doi:10.1111/rssb.12388.

2. Zou Y, Carbonetto P, Wang G, Stephens M. Fine-Mapping from Summary Data with the “Sum of Single Effects” Model. PLoS Genetics. 2022;18(7):e1010299. doi:10.1371/journal.pgen.1010299.

| Gene | Variant (pos, alleles) | $z$ | $z$-dev | log LR | Flag | Strand | $a$ | $PIP_{anch}$ | $PIP_{ens}$ | $PIP_{warm}$ |
|---|---|---|---|---|---|---|---|---|---|---|
| *ARSA* | 50716030 T/C | −0.77 | −10.7 | 1.6 | N | N | −0.82 | 0.156 | 0.883 | 1.000 |
| *ARSA* | 50625049 T/C | −12.33 | −8.0 | −29.9 | N | N | −3.07 | 1.000 | 1.000 | 1.000 |
| *ARSA* | 49778206 G/A | 2.23 | 7.3 | 4.8 | Y | N | −0.42 | 0.006 | 0.001 | 1.000 |
| *ARSA* | 49728364 G/C | −2.64 | −4.8 | −2.8 | N | Y | −0.67 | 0.033 | 0.094 | 0.875 |
| *ARSA* | 49732867 G/A | −2.64 | −4.8 | −2.8 | N | N | −0.60 | 0.033 | 0.126 | 0.875 |
| *ARSA* | 50628562 C/T | −5.59 | 1.7 | −19.3 | N | N | −0.18 | 0.020 | 0.857 | 0.997 |
| *ARSA* | 50617472 C/T | 1.97 | 1.4 | −7.4 | N | N | 0.97 | 0.160 | 0.791 | 0.521 |
| *ARSA* | 50718304 G/A | 3.88 | 0.7 | −13.2 | N | N | −0.35 | 0.002 | 0.002 | 1.000 |
| *ARSA* | 50725687 C/T | 4.32 | 0.2 | −15.4 | N | N | 0.80 | 0.004 | 0.811 | 0.000 |
| *RRP7A* | 42500030 G/A | 2.39 | 6.2 | 4.0 | Y | N | 2.49 | 1.000 | 1.000 | 1.000 |
| *RRP7A* | 42548713 A/G | −3.77 | −5.6 | −4.6 | N | N | 0.11 | 1.000 | 1.000 | 1.000 |
| *RRP7A* | 42539806 A/G | 1.73 | −4.7 | −10.1 | N | N | 1.16 | 1.000 | 1.000 | 1.000 |
| *RRP7A* | 42545697 A/G | −3.51 | −4.5 | −6.1 | N | N | 0.58 | 1.000 | 0.000 | 0.000 |
| *RRP7A* | 42629188 C/G | 0.96 | 4.1 | 4.6 | N | Y | −0.62 | 1.000 | 0.000 | 0.000 |
| *RRP7A* | 42509764 G/A | −11.67 | −3.8 | −36.2 | N | N | −1.16 | 0.833 | 0.984 | 0.770 |
| *RRP7A* | 42547475 G/A | 9.65 | 2.9 | −35.8 | N | N | −0.69 | 1.000 | 1.000 | 1.000 |
| *RRP7A* | 42553742 C/G | 7.08 | 2.6 | −33.9 | N | Y | 2.18 | 1.000 | 0.000 | 0.000 |
| *YDJC* | 21715668 G/A | 0.11 | 7.9 | 0.8 | N | N | 1.00 | 0.050 | 1.000 | 1.000 |
| *YDJC* | 22362135 A/G | 1.34 | 3.3 | 0.3 | N | N | −1.07 | 0.004 | 0.986 | 0.961 |
| *YDJC* | 21629564 G/A | −6.19 | −3.2 | −26.7 | N | N | −3.41 | 0.997 | 0.000 | 0.983 |
| *YDJC* | 21715468 T/C | −3.61 | −1.1 | −17.3 | N | N | −0.28 | 0.001 | 1.000 | 1.000 |
| *LGALS9* | 27631065 A/G | −10.68 | −4.2 | −35.7 | N | N | −2.80 | 1.000 | 0.997 | 0.998 |
| *LGALS9* | 27630899 C/T | 7.43 | 3.4 | −33.6 | N | N | 2.80 | 0.001 | 0.264 | 0.535 |
| *LGALS9* | 27635397 G/C | −1.05 | 2.7 | −4.9 | N | Y | −0.70 | 0.999 | 0.319 | 0.010 |

**Table B.** Reference-LD consistency of the high-PIP movers at the four annotation-responsive loci (GTEx Lung, 1000 Genomes EUR reference panel). Positions are GRCh38 on the gene's chromosome (*ARSA*, *RRP7A*, *YDJC*: chr22; *LGALS9*: chr17). $z$ is the observed marginal $z$-score; *$z$-dev* is the standardised deviation of $z$ from the value predicted by its LD neighbours under `kriging_rss` (large $|z\text{-dev}|$ indicates the observed $z$ is inconsistent with the reference LD); $\log \text{LR}$ is the `kriging_rss` outlier log-likelihood ratio. *Flag* is the `susieR` allele-flip flag, which fires only when $\log \text{LR} > 2$ *and* $|z| > 2$; note that the most damaging cases (e.g. *YDJC* 21715668 and *ARSA* 50716030) are unflagged solely because $|z| < 2$, so the continuous $z$-dev is the more informative quantity. *Strand* marks strand-ambiguous (A/T or C/G) SNPs. $a$ is the processed signed AlphaGenome annotation. $PIP_{anch}$, $PIP_{ens}$, and $PIP_{warm}$ are the posterior inclusion probabilities under the annotation-agnostic SuSiE anchor, the SuSiNE ensemble aggregate, and the annotation-free warm refit, respectively.

| | $n$ loci | median $s$ | max $s$ | median $n$-ratio |
|---|---|---|---|---|
| Non-responder | 16 | 0.074 | 0.212 | 11.46 |
| Responder | 4 | 0.138 | 0.191 | 11.65 |

**Table C.** `estimate_s_rss` consistency parameter, responders versus non-responders. Larger $s$ indicates greater $z$–LD inconsistency. Values are computed on the top-$|z|$ subset of each locus ($\leq 1500$ variants), so $s$ is a screen rather than a calibrated per-locus value, and the responder/non-responder gap is suggestive rather than decisive. The $n$-ratio is the within-locus ratio of maximum to minimum effective per-variant sample size, reported because heterogeneous $n_j$ is one further source of $(z, R, n)$ inconsistency. The method-of-moments residual-variance and Cholesky columns from the source run are omitted: they returned uniform NA/failure across all loci, an artifact of the unregularised LD object used in the one-off validation rather than a property of the loci.

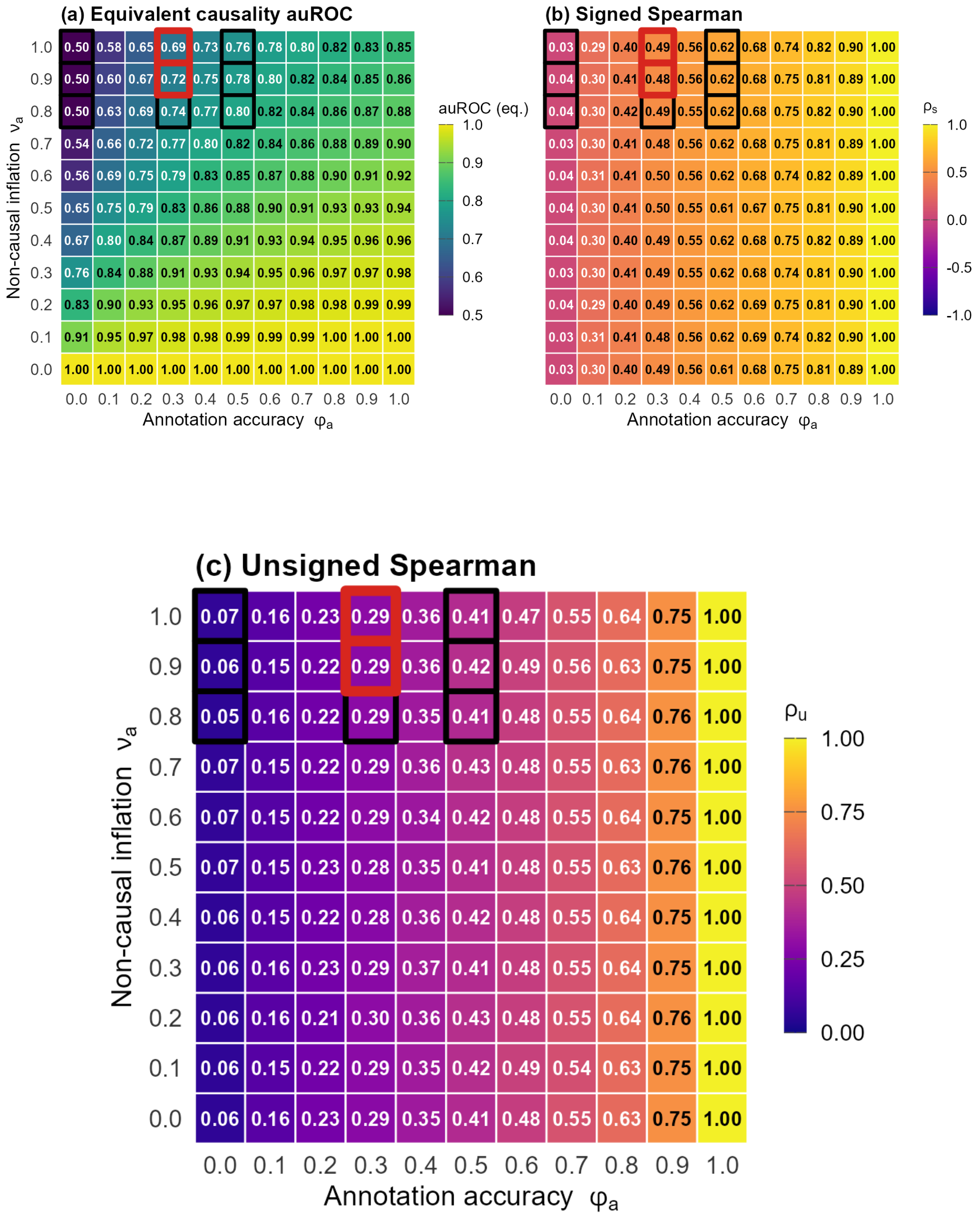

**Fig A.** Annotation calibration over the $11 \times 11$ $(\phi_a, \nu_a)$ grid. (a) Equivalent causality auROC from pooled sparse-versus-null AUPRC. (b) Signed Spearman correlation between annotation and effect over the 23 causal variants. (c) Unsigned Spearman correlation over the same variants. Black outlines mark the baseline $3 \times 3$ working grid; red outlines bracket the $\nu_a \approx 0.95$ calibration at $\phi_a = 0.3$.

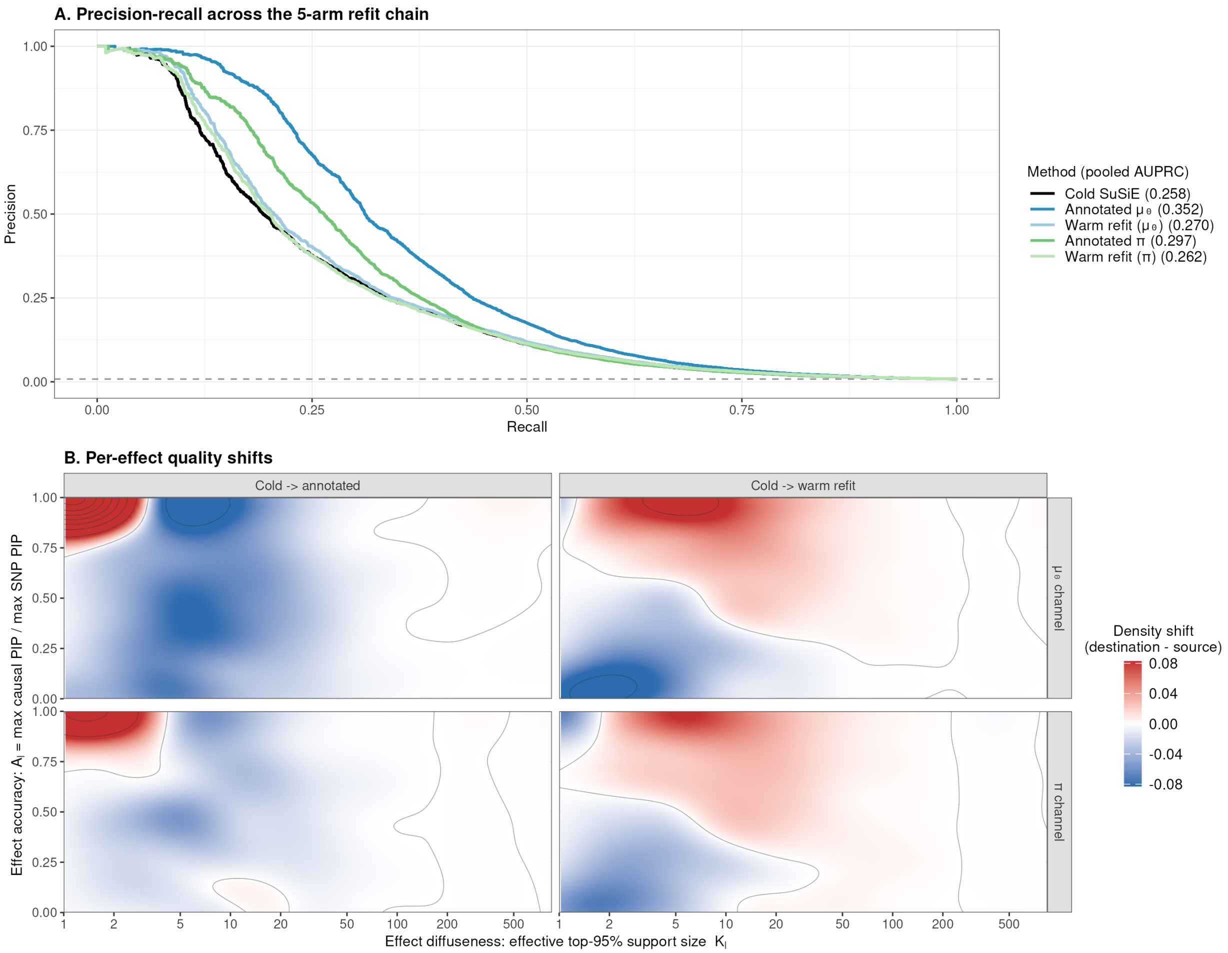

**Fig B.** Pooled performance and per-effect quality shifts for the 5-arm refit chain in the high-quality annotation regime $(\phi_a, \nu_a) = (0.5, 0.9)$ on the 600 oligogenic-architecture simulations. The chain is cold SuSiE-vanilla, cold SuSiNE-functional-$\mu$ at $c = 0.6$, SuSiE-vanilla warm refit from $\mu$, cold SuSiE-functional-$\pi$ at $\tau = 0.464$, and SuSiE-vanilla warm refit from $\pi$; all five arms use $\sigma_0^2 = 0.01$, with $c$ and $\tau$ the best-AUPRC values within this regime from the baseline screen. **(A)** Pooled precision-recall curves across the five arms; positives are the top-8 causal variants per dataset by $|\boldsymbol{\beta}|$ and negatives are the true non-causal variants; lower-effect polygenic-tier causals are excluded from scoring, matching the baseline-screen convention. Per-method pooled AUPRC is in the legend; the dashed line marks the random-classifier baseline at the pooled prevalence. **(B)** Per-effect joint-density shifts on the (effect diffuseness $K_\ell$, effect accuracy $A_\ell$) plane, pooled across effects and weighted by per-effect explained-variance fraction. Red shading marks regions where effects accumulate in the destination fit relative to the source; blue marks regions effects vacate. Columns compare cold vanilla $\to$ cold annotated and cold vanilla $\to$ warm-vanilla refit; rows show the $\mu$ and $\pi$ chains. Fill scale is shared across panels; the diffuseness axis is on $\log K_\ell$.

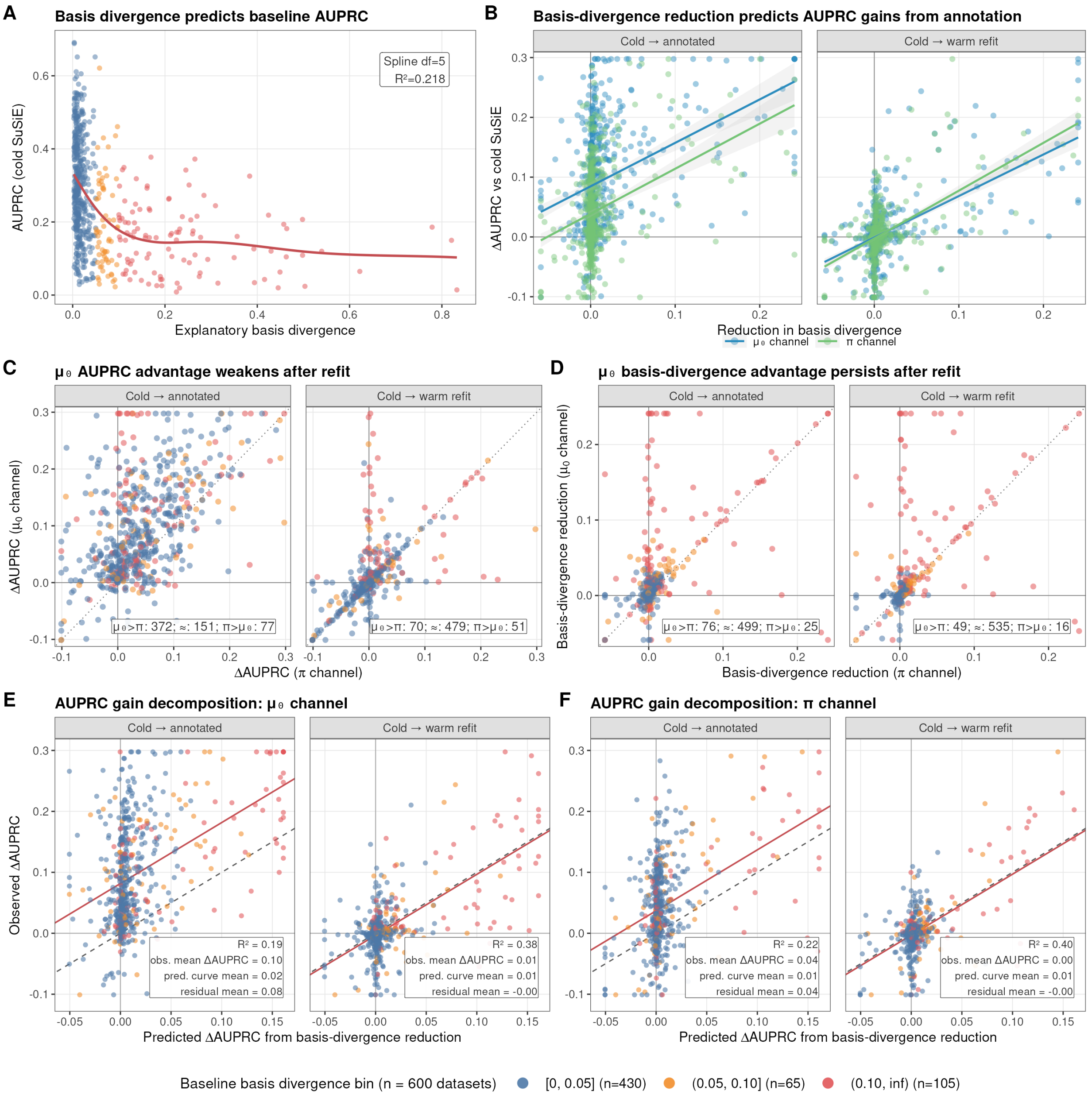

**Fig C.** Truth-basis divergence as a mechanism-level diagnostic on the same 5-arm refit chain as Fig. B. Divergence is the variance-weighted $1 - r^2$ distance (Eq. 50) between matched fitted components and the synthetic top-23-causal truth basis. **(A)** Cold SuSiE AUPRC versus baseline truth-basis divergence; the df-5 spline explains $R^2 = 0.216$. **(B)** Per-dataset AUPRC gain versus reduction in truth-basis divergence, split by channel and by whether the annotation prior remains active. **(C,D)** Direct $\mu$-versus-$\pi$ comparisons for AUPRC gains and divergence reductions. **(E,F)** Decomposition of AUPRC gain into the component predicted by movement along the baseline divergence–AUPRC spline plus a residual component. Warm-refit gains are smaller but more directly tied to basis movement, whereas cold annotated gains also include prior-dependent within-basin reweighting.

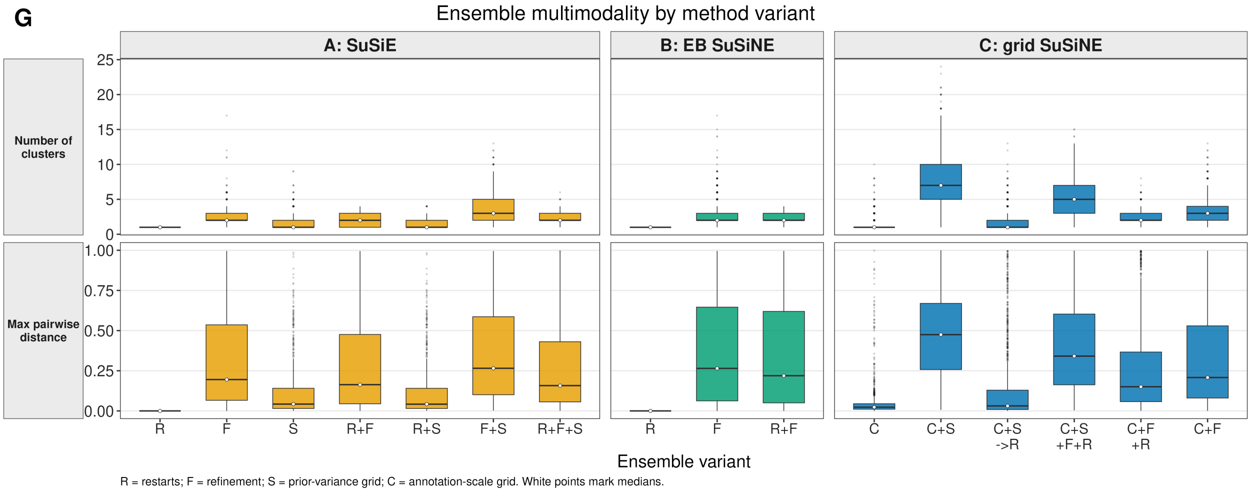

**Fig D.** Ensemble multimodality by method variant, across the 600 oligogenic phenotype datasets derived from 150 genotype matrices. Each box is the per-dataset distribution of a multimodality metric (rows) for a given ensemble construction strategy (within-panel $x$-axis: exploration-axis suffix), grouped by model family into columns (A: SuSiE; B: EB SuSiNE; C: grid SuSiNE). Lower values indicate less posterior multimodality across ensemble members; white points mark medians. Suffixes: R restarts, F refinement, S prior-variance grid, C annotation-scale grid. C-CSR is the exception: it is not an additional restart axis, but the C-CS functional-prior search followed by default zero-prior refits from each source fit's exact per-effect posterior moments, so its multimodality summarizes the annotation-discovered basins after prior removal.

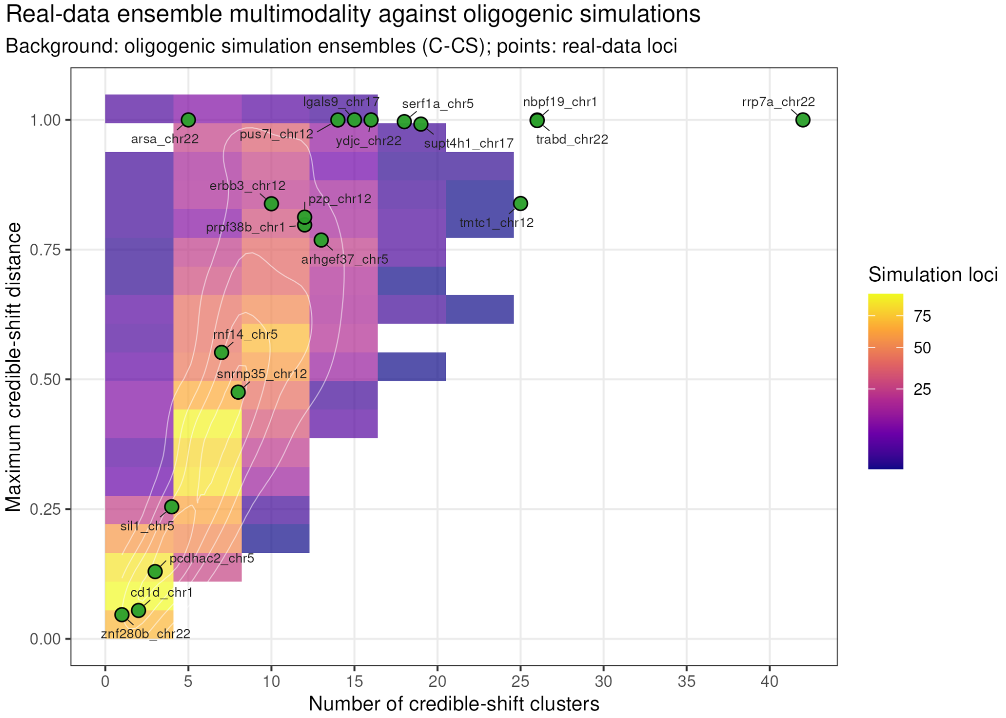

**Fig E.** Real-data ensemble multimodality benchmarked against oligogenic C-CS simulation ensembles. The simulation background uses the same credible-shift diagnostics as cluster-weight aggregation: number of clusters at distance threshold 0.05 and maximum pairwise credible-shift distance. Green points are the 20 GTEx Lung loci. Several real loci fall in the high-cluster or high-distance tail, supporting the use of cluster-aware aggregation in the real-data study.

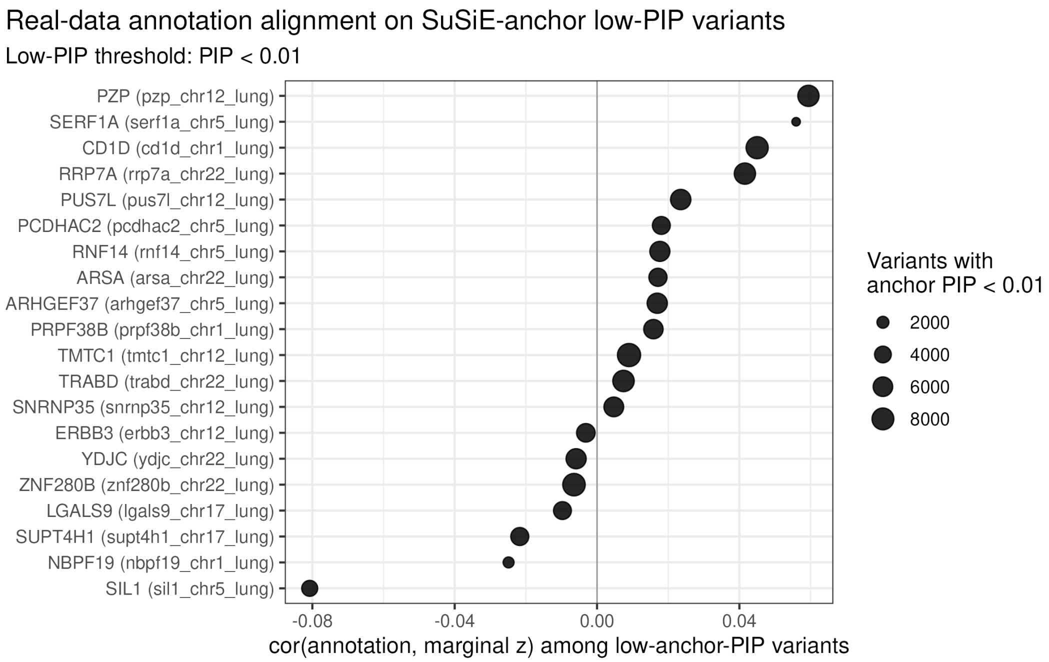

**Fig F.** Real-data internal null-alignment diagnostic. For each of the 20 GTEx Lung loci, variants with annotation-free SuSiE anchor PIP $< 0.01$ define a low-baseline-evidence subset, and the plotted value is the correlation between AlphaGenome annotation $a_j$ and marginal association statistic $z_j$ within that subset. The summary values were near zero across the panel: median correlation 0.0124, median absolute correlation 0.0174, and maximum absolute correlation 0.0807. This argues against strong null annotation–association alignment in this panel but is not an external negative control.

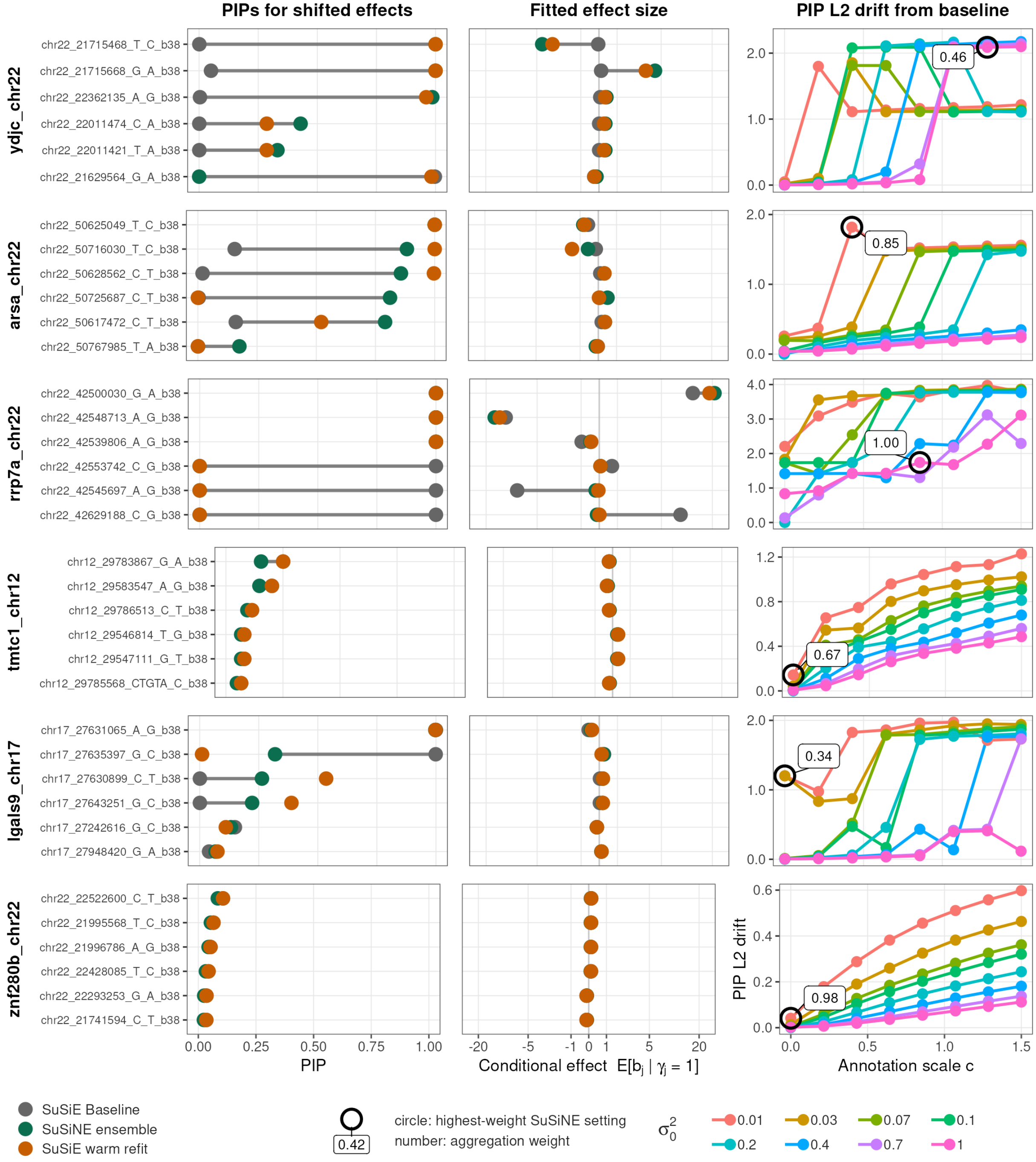

**Fig G.** Zoom on six GTEx Lung loci spanning the annotation-influence range: *YDJC*, *ARSA*, *RRP7A*, *TMTC1*, *LGALS9*, and *ZNF280B*. For each locus, the six displayed variants have the largest absolute anchor-to-ensemble posterior-mean shifts. Left: PIPs for the SuSiE anchor, SuSiNE ensemble, and warm-vanilla refit. Middle: corresponding conditional effects. Right: PIP L2 drift from the anchor for all 64 grid runs, plotted against prior-mean scale $c$ and colored by $\sigma_0^2$. The open circle marks the highest-weight source run. Banded or discontinuous L2 curves indicate discrete basin changes rather than smooth within-basin reweighting.
